# Metabolic regulation of copper homeostasis governs the Sec61-dependent protein translocation process in *Saccharomyces cerevisiae*

**DOI:** 10.1101/2025.08.03.668358

**Authors:** Vandana Anjana, Smriti Anand, Prateeksha Thakur, Rajshree Pal, Santoshi Acharjee, Sugandh Sharma, Sharayu Subhash Awachat, Ritika Manjhi, Devika Rejidev, Ranu Singh, Raghuvir Singh Tomar

## Abstract

Concentration of cellular labile pool of copper must be strictly regulated because disruption in copper homeostasis results in diseases. In *Saccharomyces cerevisiae*, elevated levels of labile copper impair cell viability by inhibiting Sec61-mediated protein translocation into the endoplasmic reticulum. We investigated how metabolic pathways, specifically mitochondrial respiration and autophagy, contribute to copper homeostasis and the translocation of secretory proteins. We show that copper selectively inhibits protein translocation in yeast cells grown in minimal medium but not in rich medium, highlighting a critical role of nutrients in modulating copper toxicity. Supplementation of specific amino acids suppresses the copper-induced defects in protein translocation and cell death, identifying amino acids suppressors of the copper toxicity. Using a panel of gene deletion mutants affecting mitochondrial functions, autophagy, peroxisomes, and lipid droplets, we demonstrate that metabolic pathways regulate sub-cellular distribution of copper and translocation of secretory proteins. Further, disruption of redox and pH homeostasis, and pharmacological inhibition of respiration, reveals that correct subcellular trafficking of copper is essential to prevent inhibitory effects on protein translocation process. Together, our findings provide mechanistic insights into how metabolic status influences cellular copper homeostasis and the secretory pathway of proteins, with broader implications for understanding diseases of copper metabolism.

## Introduction

Micronutrients such as copper, iron, and zinc play crucial roles in regulating a variety of evolutionarily conserved biological processes across living organisms (1–3). Both deficiency and excess conditions of these metals are associated with a range of physiological and pathological conditions (4,5). The genetic, epigenetic, and metabolic factors contribute to maintaining the homeostasis of metals, including copper (6,7). As metal homeostasis is essential to regulate the cellular functions and organismal health, identification and functional characterization of factors that regulate cellular metal concentrations can provide insights into various diseases and potential therapeutic approaches. For example, genetic factors regulate copper concentration, which is essential for normal biological processes and minimizes the potential toxicity. Disorders such as Wilson’s and Menkes diseases are linked to mutations in the ATP7B and ATP7A genes, respectively (8). Wilson’s disease results in copper accumulation due to impaired copper efflux, leading to toxicity, while Menkes disease is characterized by copper deficiency, which occurs due to a defect in the intestinal absorption of copper. Similarly, epigenetic mechanisms, including histone modifications, influence copper homeostasis by regulating the expression of genes involved in copper transport and sequestration. Yeast cells regulate cellular copper levels through transcription factors and chaperones such as Mac1p, which activate transcription of genes involved in copper uptake, such as *CTR1*, *CTR3*, and *FRE1*, in response to copper deficiency, and Ace1p induces the expression of *CUP1* and *CRS5*, encoding metallothionein proteins that sequester excess copper, protecting from toxicity (9). These regulatory mechanisms maintain copper levels within the optimal ranges, supporting cellular functions while preventing toxicity.

The interplay between molecular and cellular factors that regulate genetics, epigenetic modifications, and metabolic states determines overall metal homeostasis within cells (10). Disruptions in any of these biological processes can lead to imbalances in metal concentrations, resulting in disease conditions. For example, in yeast cells, mutations in histone proteins have been shown to increase the labile copper pool, leading to defects in Sec61-mediated protein translocation processes and cell death (11). In addition, the activity of the copper-dependent enzymes, including those that work inside mitochondria, is dependent on the cuprous (Cu1^1+^) state, which is regulated by copper reductase activity of the histone H3-H4 tetramer. Excess labile copper can disrupt the protein maturation process in the secretory pathway by inhibiting the Sec61 translocon. Higher labile copper levels lead to the accumulation of unprocessed forms of proteins, such as Gas1p and CPY, in the cytosol, suggesting that copper homeostasis must be tightly regulated for proper protein processing and cellular functions (12). Furthermore, studies suggest that the cellular concentrations of metabolic intermediates, such as amino acids and glutathione, play crucial roles in the regulation of copper homeostasis. Glutathione, a small tripeptide consisting of glutamate, cysteine, and glycine, serves as one of the primary intracellular agents that mitigate copper toxicity. Glutathione binds to metals like copper, facilitating their sequestration and preventing oxidative damage and copper induced protein translocation defects. In addition, the stoichiometric balance among metals may also contribute to cellular metal homeostasis, as some metals may compete for binding with proteins and amino acids, although with varying affinities. The cellular pool of amino acids must be maintained through the nutrients present in the growth medium, mitochondrial respiration, and autophagy, impacting the metabolic state of the cells, because the disruption of amino acid metabolism can cause diseases in humans (13–16). The process of autophagy has also been shown to regulate CLS in yeast, suggesting that aging-associated changes in the metabolism of metals, including copper, in eukaryotic organisms can affect protein homeostasis by modulating the translocation process of proteins. In yeast, *Saccharomyces cerevisiae*, there are additional mechanisms, such as import, export, and compartmentalization, that play an important role in the regulation of the cellular pool of amino acids (17,18). On the other hand, the cellular labile pool of essential metals is mainly regulated by the expression of genes that encode metal-specific transcription factors, membrane transporters, cytoplasmic metallothionein and chaperone proteins, and vacuole membrane transporters (19–21). As a protective mechanism, the excess quantity of many of the metals in yeast cells is sequestered inside vacuoles. In addition, according to a study in yeast, the alkaline pH of the growth medium decreases the availability of copper and iron due to their low solubility, leading to a slow-growth phenotype of the cells (22). Since the pool of amino acids and metals inside cells is very critical for life processes, the concentration of both must be regulated to avoid the toxic effects of metals. However, mechanisms that regulate the biochemical or genetic interactions between essential metals and amino acids are not completely understood.

Mitochondria are one of the major copper-utilizing cell organelles (23–26). Many of the mitochondrial proteins require copper to regulate their activities. For example, cytochrome c oxidase (Cox1p), as part of the mitochondrial electron transport chain (ETC), is essential for aerobic respiration. Cu, Zn superoxide dismutase (Sod1p) detoxifies superoxide radicals by converting them into hydrogen peroxide and molecular oxygen. Ccs1p is a copper chaperone responsible for transporting copper to mitochondrial Sod1p. Cox17p is another copper chaperone that conveys copper to the mitochondria and works alongside Sco1p and Cox11p to facilitate the incorporation of copper into cytochrome c oxidase inside the mitochondria. These studies indicate that activity and the concentration of copper-requiring proteins, trafficking of copper between organelles, and stoichiometry among essential metals are the important factors that determine the labile pool of copper inside cells. In addition, the fusion and fission processes that regulate the biogenesis of mitochondria, vacuoles, and other cell organelles are additional mechanisms that may impact the availability of labile copper and other copper competing metals in the cytoplasm. The mitochondrial number and size of the vacuoles in yeast cells are regulated in response to external and internal factors. For example, the availability of glucose and nitrogen in the growth medium determines the rate of glycolysis, mitochondrial respiration, and autophagy (27,28). Superoxide radicals are generated inside mitochondria as part of the metabolic reactions and upon treatment with DNA-damaging agents. To protect from the toxic effects of superoxide radicals and redox imbalance, the demand for copper inside mitochondria may increase to enhance the activity of the copper-dependent enzymes such as Sod1p and Cox1p. For example, studies have shown that the addition of copper protects the *SOD1* deleted yeast cells from the toxic effects of MMS (methyl methane sulfonate), a DNA-damaging agent (29). Another study has demonstrated that Sod1 binds to promoters in the nucleus to regulate the expression of antioxidant and repair genes (30). In addition, mutations at several locations in Sod1 have been identified that cause amyotrophic lateral sclerosis disease (31).

This comprehensive study was conducted to identify the metabolic factors regulating copper homeostasis and the protein translocation process. Previously, by using yeast cells, we demonstrated that an increase in the labile copper concentrations in the cytoplasm of histone mutants inhibits the Sec61-mediated translocation of secretory proteins. We also observed copper-induced protein translocation defect in wild-type yeast cells, specifically during the exponential growth phase, indicating that the balance between glycolysis and mitochondrial respiration can regulate the protein translocation process. Therefore, we hypothesized that an intricate balance between respiration and autophagy mechanisms may play a crucial role in maintaining the labile copper concentration. We started investigations by supplementation of amino acids one at a time in the growth medium to identify the amino acid suppressors of copper-induced defects in the protein translocation process and slow growth phenotype. These experiments led to the identification of a few of the amino acids suppressing the copper-induced defects in the protein translocation process, but to different extents; poor, moderate and strong. Supplementation of some of the amino acids did not suppress the toxic effects of copper. Screening of gene deletion mutants defective in mitochondrial respiration, vacuoles, autophagy, peroxisomes, lipid droplets, redox, and pH balance led to the identification of conserved cellular factors regulating the copper homeostasis and protein translocation process. In addition, results upon the pharmacological inhibition of mitochondrial respiration and V-ATPases, as well as minimal and rich growth medium, further confirm that the cellular pool of amino acids, redox, and pH balance are the key mechanisms to regulate the copper homeostasis; disruption in these mechanisms affects the protein translocation process. Together, our findings provide evidence that metabolic regulation of copper homeostasis is necessary to prevent the defects in protein translocation process.

## Results

### Growth medium composition modulates copper homeostasis and protein translocation process in yeast cells

*Saccharomyces cerevisiae* yeast model organism is extensively used to identify the factors of different metabolic pathways because it is a simple single cellular eukaryotic organism, with easy genetic manipulations and rapid growth. To study different biological processes, a variety of growth media are utilized to examine the growth of yeast cells. In general, two major types of growth media are used for the cultivation of yeast cells, YPD (Yeast Extract, Peptone, Dextrose) rich medium contains complete set of nutrients which is in general used for vigorous growth and minimal media also known as synthetic complete or SC medium is utilized for specific experimental applications such as to identify the nutritional requirements. In addition, selective growth media are used for the selection of specific yeast strains. Since the nutrient composition of YPD and minimal growth media is distinct, the metabolic profiles of the yeast cells grown in these growth media will be different. YPD, a complex medium, contains readily available nutrients, leading to faster growth of the cells. In contrast, yeast cells in the SC medium need to synthesize their own amino acids from glucose, leading to a different metabolic profiling. Availability of nutrients can influence the rate of metabolic pathways and growth. Rich growth medium promotes faster growth, whereas nutrient-deficient conditions may lead to cell cycle arrest, increased chronological life span, and activate autophagy. Thus, the composition of nutrients in the growth medium impacts almost all the fundamental biological processes, such as respiration, autophagy, and the cell cycle. However, the significance of nutrients of the growth medium in the copper homeostasis remains poorly understood. The minimal growth media should have a smaller or lower concentration of free amino acids in comparison to the YPD rich medium. Since a few of the amino acids, such as cysteine, histidine, and reduced glutathione, can directly bind with copper, the concentration of labile copper will be higher in cells grown in minimal media than in cells grown in rich medium. As we know that an increase in labile copper concentration in the cytoplasm can inhibit the Sec61-mediated protein translocation process, we hypothesize that copper-induced defects in the protein translocation process will be much higher in the cells grown in minimal media than in the cells grown in rich growth media (Figure 1D).

**Figure 1:**
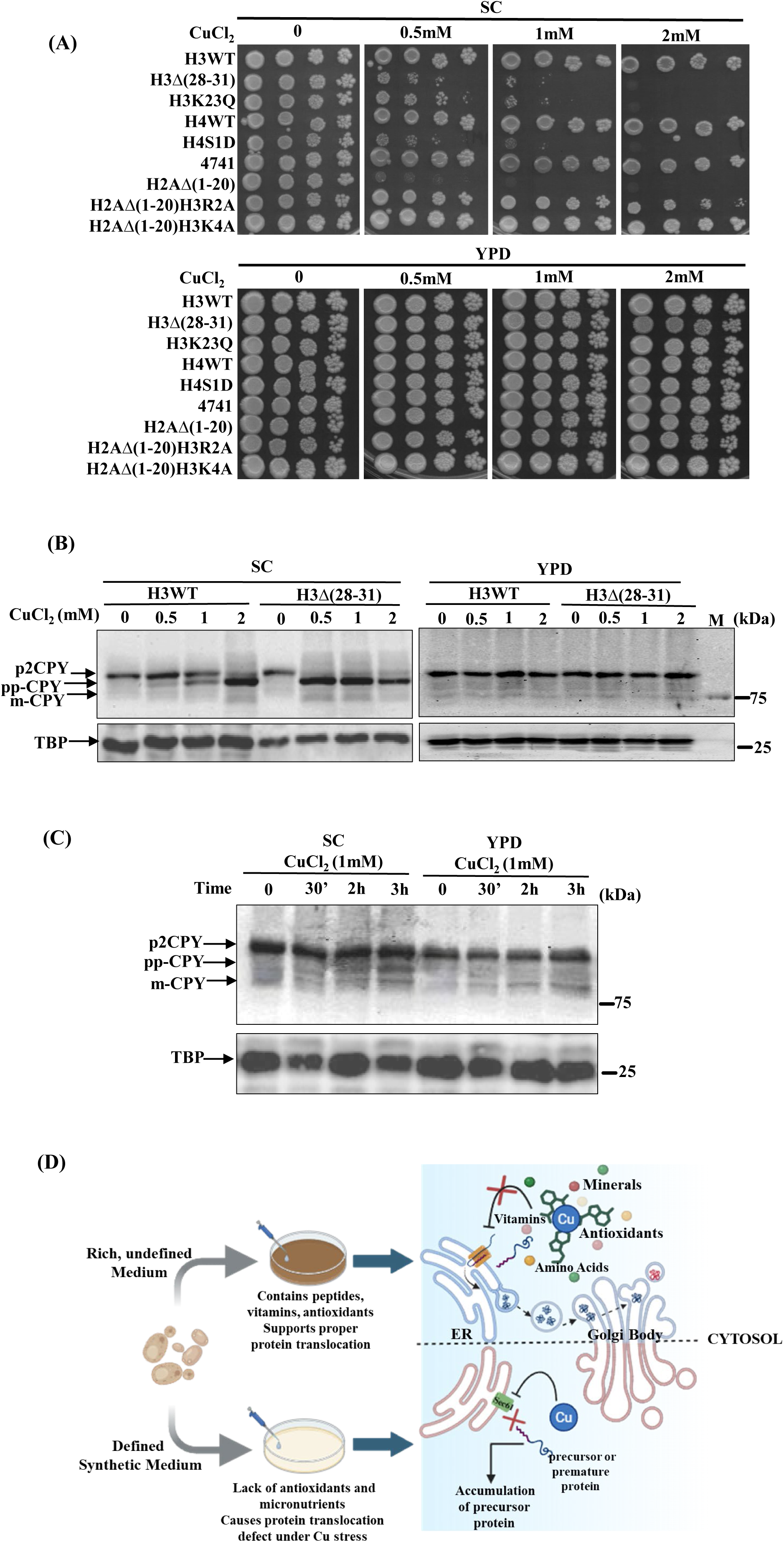
The growth medium composition regulates copper homeostasis and protein translocation in yeast. A. Wild type and a few of the copper-sensitive histone H3 mutants were grown overnight in SC medium, and cells were adjusted to an OD_600_ of 1. A 10-fold serial dilution was prepared: 1:10, 1:100, 1:1000. 3-4 microliters of cells from each dilution were spotted on SC agar or YPD agar solid growth medium with and without CuCl_2_ (concentrations as indicated in the figure). Growth was recorded after 72 hrs of incubation. B. Overnight grown cultures of wild type and a histone H3 mutant expressing myc-tagged CPY were seeded at 0.2 OD_600_ in SC and YPD, grown till 1OD6_00_, untreated and treated with copper chloride (0.5, 1.0, and 2.0 mM) for 2 hours, harvested, whole extracts were prepared, and the translocation of CPY (myc-tagged) was analyzed by western blotting. C. Translocation of CPY was analyzed by western blotting in wild type cells grown in SC and YPD growth medium in same manner as in ‘B’, exponentially growing cells were treated with 1 mM of copper chloride for 30 minutes, 2 hours and 3 hours and extracts were prepared for the western blotting of CPY. D. Hypothesis predicting the role of nutrients of the growth medium in the regulation of copper homeostasis mediated protein translocation process.

To understand the impact of minimal and rich growth media on the protein translocation process, we utilized wild type and a few of the copper-sensitive and non-sensitive histone H3, H4, and H2A mutants of *Saccharomyces cerevisiae*. Following strains were utilized; H3 wild type, H4 wild type, BY4741 wild type, H3 mutants [H3K23Q, and H3Δ(28–31)], H4 mutants, H4 (H4S1D) and H2A mutants [H2AΔ(1–20), H2AΔ(1–20)H3R2A and H2AΔ(1–20)H3K4A]. Among H2A mutants, only one mutant, H2AΔ(1–20) is copper sensitive and other mutants, H2AΔ(1–20)H3R2A and H2AΔ(1–20)H3K4A are copper non-sensitive, their growth in presence of copper is same as that of wild type cells. All these strains were grown overnight in SC growth media. 1OD_600_ cells of each yeast strain were serially diluted and spotted onto SC and YPD growth media with and without copper chloride. The growth of cells was recorded after 72 hours. We observed almost no difference in the growth pattern of all the strains in SC and YPD media in copper-untreated cells. However, in the presence of copper (0.5 mM, 1.0 mM, and 2.0 mM), the growth of copper-sensitive histone mutants, as reported earlier, was significantly reduced only in the SC media, but we did not observe any noticeable effect on the growth in YPD media (Figure 1A and S1A). As the nutrients in SC and YPD media are different, the labile concentration of copper in the cells grown in these two media conditions will be different, which may affect the protein translocation process to different degrees. So, to examine the effect of SC and YPD media on the protein translocation process in copper-untreated and copper-treated conditions, we utilized wild type and only one of the copper-sensitive histone H3 mutants, H3Δ(28–31). We used only one copper-sensitive histone mutant just to reduce the number of samples for the western blotting. The overnight-grown cultures of myc-tagged CPY-expressing cells were seeded in SC and YPD media at 0.2OD_600_ and grown until 1.0OD_600._ Subsequently, cells were allowed to grow for two hours in the respective media in untreated and copper chloride-treated conditions. Cells were harvested, whole cell extracts were prepared as outlined in the materials and methods section, and western blot analysis was conducted to detect the different forms: cytosolic, Golgi, and vacuolar of carboxypeptidase Y (CPY), an example of the secretory pathway of proteins. Interestingly, we observed a very strong band of pp-CPY (cytosolic non-glycosylated form) in copper-treated conditions, which was absent in copper-untreated conditions in the cells grown in SC media. We also observed a faster migrating protein band of CPY in copper treated conditions which we have labelled as m-CPY (mature CPY in vacuole) based on the molecular weight. On the other hand, we observed only the Golgi glycosylated form of CPY (p2CPY) in untreated as well as copper-treated conditions in the cells grown in YPD media (Figure 1B, 1C, and S1B, S1C, and S1D). The above results clearly demonstrate that the availability of nutrients in the growth media controls the copper homeostasis-mediated protein translocation process. Nutrients can impact the rate of metabolic pathways, including respiration and autophagy. Therefore, we further focused our investigations to identify the role of amino acids, mitochondrial respiration, autophagy, peroxisome, redox, pH balance, and lipid droplets in regulating the complex mechanism of copper homeostasis and protein translocation process.

### Screening identifies specific amino acid suppressors of copper-mediated defects in protein translocation process

Regulation of sub-cellular distribution and concentration of the amino acid pool is essential for proper cellular functions. Multiple cellular mechanisms are involved in maintaining amino acid homeostasis. Import, synthesis, sequestration, utilization, mitochondrial respiration, and vacuolar pH are important mechanisms to maintain amino acid homeostasis. As amino acids are the primary building blocks for living systems, all 20 amino acids must be concomitantly available for the biosynthesis of proteins. In addition to protein synthesis, the role of all 20 amino acids in other biological functions, such as signalling mechanisms and metal homeostasis, is not completely understood. Many of the amino acids also play important roles in maintaining the redox balance (32). Some amino acids, like histidine and cysteine, can directly bind with copper and zinc, impacting cellular processes. It is quite challenging to establish the functional relationship between metabolic pathways of amino acids and copper homeostasis because the biosynthesis pathways of amino acids involve a complicated network (33). The metabolic intermediates of the biosynthesis pathway of one amino acid can merge into the pathway of another amino acid. Similarly, defects in the metabolic pathway of one amino acid can limit or promote the rate of another amino acids metabolic pathway. As an increase in labile copper concentration inhibits Sec61-mediated translocation of secretory proteins, we hypothesize that the cellular amino acid pool can modulate the copper homeostasis. In other words, the presence or absence of certain amino acids in growth media can affect the copper homeostasis and protein translocation process. For example, previously we observed complete rescue of the yeast cells from the copper-induced cell death and defects in the protein translocation process upon supplementation of cysteine and histidine in the growth media. Studies suggest that cysteine can reduce copper (II) to copper (I), and histidine has a strong chelation property. Due to these unique properties, we found these two amino acids to be strong suppressors of copper-mediated toxicity and defects in the protein translocation process. However, the significance of remaining amino acids in copper homeostasis and the protein translocation process is yet to be explored. Therefore, we hypothesized that supplementation of remaining amino acids individually in the growth media may be helpful to identify the amino acids that can suppress copper toxicity and inhibit defects in the protein translocation process (Figure 2F).

**Figure 2:**
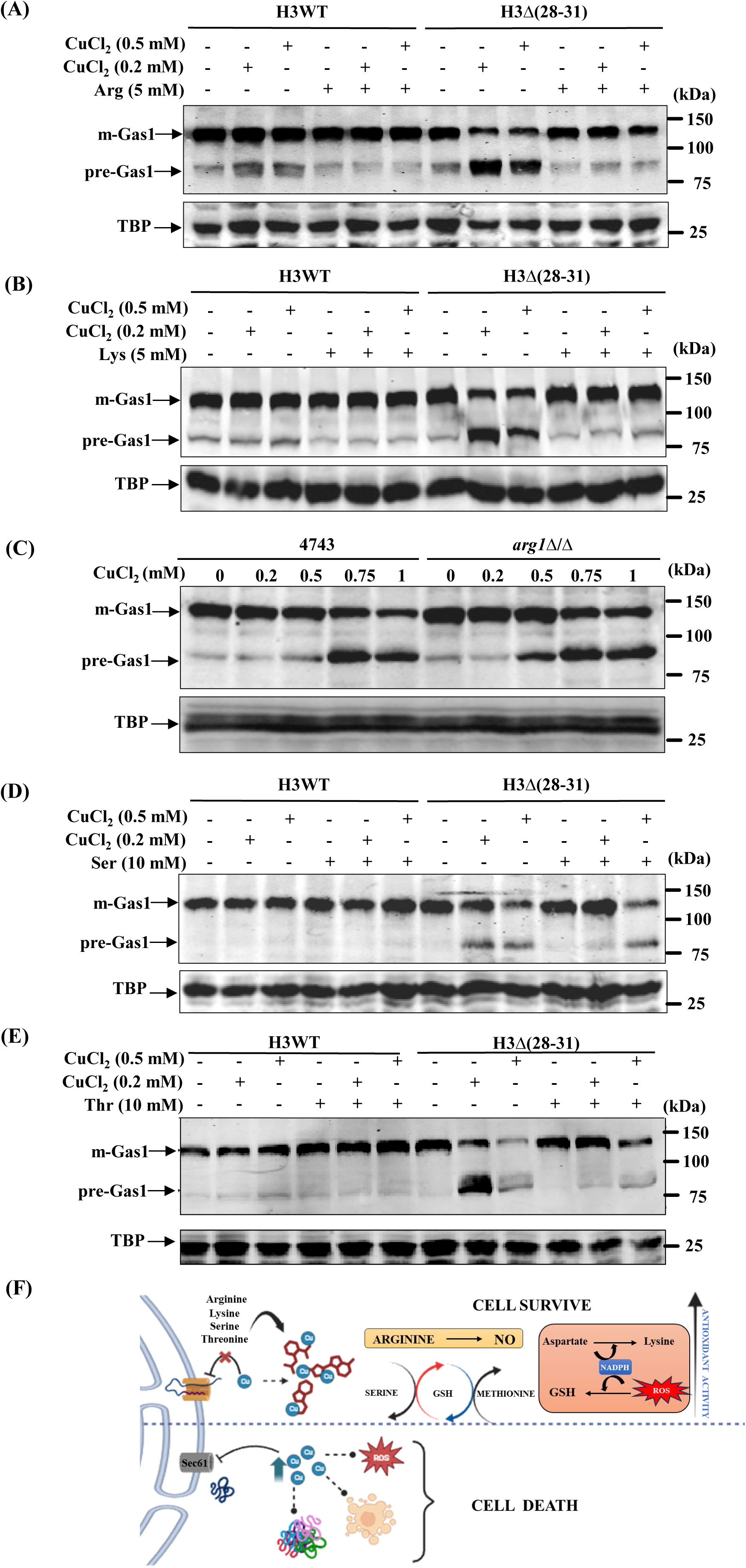
Screening identifies amino acids that mitigate copper-induced defects in protein translocation. Overnight grown cultures of Gas1-GFP plasmid transformed wild type, and a histone mutant were grown seeded at 0.2OD6_00_ in SC growth medium and grown till exponential phase, subsequently treated and untreated as indicated, extracts were prepared, and protein translocation was examined by western blotting of Gas1 using anti-GFP primary antibody. A. As indicated in the figure, exponentially growing cultures of wild type and a histone mutant were cultured in the presence and absence of 0.2 mM of copper, 0.5 mM of copper, 5 mM of arginine, and cotreatment with 0.2 mM/0.5 mM of copper plus 5 mM of arginine. B. Same as ‘A’ but supplemented with 5 mM of lysine. C. Gas1 transformed wild type and *arg1*Δ*/*Δ deleted cells were seeded at 0.2OD_600_ and grown till 1OD_600_, subsequently cultured for two hours in the absence or in the presence of copper (0.2, 0.5, 0.75, and 1.0 mM). D. As indicated in the figure, exponentially growing cultures (1OD_600_) of wild type and a histone mutant were grown for 2 hours in untreated and treated conditions, 0.2 mM, 0.5 mM of copper, 10 mM of serine, and cotreatment with 0.2 mM/0.5 mM of copper plus 10 mM of serine. E. Exponentially growing cultures (1OD_600_) of wild type and a histone mutant were grown for 2 hours in untreated and treated conditions, 0.2 mM, 0.5 mM of copper, 10 mM of threonine, and cotreatment with 0.2 mM/0.5 mM of copper plus 10 mM of threonine. F. Hypothesis indicating the role of amino acids in regulating the labile copper concentration.

Further to identify the amino acid suppressors of the copper-induced defects in growth and protein translocation process via directly or indirectly modulating copper metabolism, we employed a few very basic experimental approaches. First, tested the growth of yeast cells in absence (drop out) or upon exogenous supplementation of each amino acids one at a time in the growth medium, second, we tested the growth of a few gene deletion mutants of amino acid biosynthesis pathways and third, the protein translocation process was examined upon supplementation of only those amino acids which suppressed the toxic effects of copper on the growth of cells. We focused our investigations on gene deletion mutant cells defective in biosynthesis pathways of amino acids (leucine, lysine, cysteine, histidine, methionine, arginine, and glutathione) by testing their growth in SC media in the absence and presence of copper chloride. In addition, we also tested the growth of a few gene deletion mutants defective in amino acid permeases, transcription factors, S-adenosyl synthetase, membrane transporters, cystathionine to homocysteine converter, peroxisomal cystathionine, amino acid starvation sensor, and heavy metals transporter. We detected a slow growth phenotype in a few of the gene deletion mutants defective in amino acid metabolism upon the addition of copper. For example, gene deletion mutants of glutathione (GSH2), lysine (Lys7/Ccs1 and Lys14), and arginine (Arg1, Arg2 and Can1) were found to be slow growing in SC media in the presence of copper chloride, and the *lys12*Δ*/*Δ and *skn1*Δ*/*Δ gene deletion mutants were slightly copper resistant (Figure S2A). These observations suggest that amino acid metabolism plays a crucial role in copper homeostasis. Further, we examined the growth of wild-type and a few of the histone mutant cells in the absence and presence of copper upon supplementation of specific concentrations of individual amino acids that showed rescue in the SC media (Figure S2B, S2C, and S2D). According to the difference in the degree of suppression of the copper induced defect on the growth of cells upon supplementation of individual amino acids, we have divided amino acids into three groups; very good suppressors (lysine and arginine), good suppressors (serine and threonine) and weak suppressors (glycine, aspartic acid and glutamine). Remaining amino acids, valine, leucine, isoleucine, proline, tryptophan, phenylalanine, tyrosine, and glutamic acid did not show any effect on the growth of cells in the presence of copper. To gain more insight, we also tested the effect of copper on the growth of cells in amino acid dropout growth media in which serine, lysine, arginine, or glycine were omitted individually. However, the effect of copper on the growth of cells in the absence or presence of these amino acids was found to be almost the same as that of complete SC media, suggesting that probably crosstalk between the metabolic pathways of amino acids compensates for the dropouts (Figure S2E). The above results indicate that the concentration of some of the amino acids is very critical to maintain the copper homeostasis, suppresses the copper-induced defects in growth, probably by suppressing the defects in the protein translocation process. Thus, we went ahead and tested the impact of a few amino acids that suppresses the copper toxicity; lysine and arginine (very good suppressors), serine and threonine (good suppressors) and glycine (weak suppressor) on the protein translocation process in absence and presence of copper by western blotting using two yeast strains, wild type and a copper sensitive histone mutant, H3Δ(28–31). Interestingly, we observed that the supplementation of lysine and arginine suppresses the copper-induced defects in the protein translocation process. We observe two bands of a Gas1 secretory protein in cells upon copper treatment, larger molecular weight top band (mature glycosylated) and smaller molecular weight bottom band (immature cytosolic non-glycosylated) whereas in copper untreated condition, we detected only one major top band of Gas1 which is a mature glycosylated form (Figure 2A, 2B and S2G, S2H, S2L, S2M). On the other hand, we observed only the mature form of Gas1 in cells co-treated with copper and amino acid suppressors of copper toxicity (lysine and arginine). These results suggest that supplementation of lysine or arginine suppresses the toxic effect of copper on the growth of cells by preventing the inhibitory effect of copper on the protein translocation process. Further to confirm the role of these amino acids in the rescue of cells from copper-induced defects in protein translocation, we tested the translocation process in the absence and presence of copper in the *arg1*Δ*/*Δ gene deletion mutant and compared it with the wild-type cells. We observed about 20-30% more defects in the *arg1*Δ*/*Δ gene deletion mutant at 0.5 mM copper chloride treatment in comparison to wild-type cells, suggesting that arginine plays a crucial role in suppressing the copper-induced defects in the protein translocation process (Figure 2C and S2I, S2N). We also observed that supplementation of serine, threonine, and glycine in the growth media suppresses the copper induced defects in the protein translocation process, but less than lysine and arginine supplementation (Figure 2D, 2E, and S2J, S2K, S2O, S2P, S2F). These results indicate that the cellular pool of amino acids can regulate the copper homeostasis-mediated protein translocation process.

### Mitochondrial activities regulate copper homeostasis and the protein translocation process

Mitochondria are the essential cell organelles that, in addition to ATP synthesis, regulate cellular metabolic pathways by providing precursors for the metabolic biosynthetic pathways (34). Numerous mitochondrial diseases exist, primarily genetic disorders resulting from mutations in both nuclear DNA (nDNA) and mitochondrial DNA (mtDNA) genes, which encodes proteins necessary for mitochondrial functions, resulting in defects in oxidative phosphorylation. Mitochondria also play essential roles in the redox balance, biosynthesis of amino acids, fatty acids, nucleotides, signalling pathways, and detoxification of reactive oxygen species (35). In addition, mitochondria regulate the metal homeostasis, such as copper, zinc, and iron, as these metals are utilized by the mitochondrial enzymes conducting the metabolic reactions. Mitochondrial membrane transporters and chaperones contribute to the copper and zinc homeostasis essential for the activity of superoxide dismutase 1 (Sod1) (36). Since many of the mitochondrial functions are heavily dependent on copper, it becomes important to maintain the physiological concentration because labile copper can inhibit the enzymes of the TCA cycle (37). Similarly, an increase in the labile concentration of copper in the cytoplasm can inhibit the Sec61-mediated translocation of secretory proteins. Therefore, we hypothesize that mitochondrial activities such as respiration, fermentation, fission, and fusion processes play an essential role in the regulation of sub-cellular concentration as well as trafficking of labile copper between cytoplasm and mitochondria (Figure 3H). It is necessary to understand the role of mitochondrial activities in the regulation of cellular copper homeostasis mediated protein translocation process because the protein translocation regulates multiple biological functions such as protein secretion, biogenesis of cell organelles, and immune response. A defect in the protein translocation process correlates with pathological conditions, including cancer and infections. To identify the role of mitochondrial activities in copper homeostasis and the protein translocation process, we performed a couple of experiments. We decided to examine effect of copper on the growth of wild-type cells in non-fermentable carbon source medium and in SC medium, copper response mutants such as *sod1*Δ*/*Δ*, ccs1*Δ*/*Δ*, atx1*Δ*/*Δ*, ccc2*Δ*/*Δ*, cup2*Δ*/*Δ*, mac1*Δ*/*Δ, petite cells, and on the antimycin-treated cells. In addition, we also tested the effect of copper on the growth of a few gene deletion mutants defective in respiration, fission, and fusion processes. First, we tested the growth of wild type along with a few of the copper-sensitive histone mutants in the absence and presence of copper in SC media containing glucose or non-fermentable carbon sources. We observed that the effect of copper is decreased on the growth of cells grown in non-fermentable carbon source medium in comparison to cells grown in normal SC glucose medium (Figure S3A). This observation suggests that the demand for copper utilization increases to support the mitochondrial respiration activities in cells grown in non-fermentable carbon source medium in comparison to cells grown in normal SC medium. To understand the effect of non-fermentable carbon sources on mitochondrial activities, we measured the expression of genes, *CAT8* and *FBP1,* which are essential to support the growth of yeast cells in non-fermentable carbon source medium. We observed significant de-repression of these genes in cells grown in non-fermentable carbon source medium (Figure 3C). As the demand for copper inside mitochondria increases during the growth of cells in a non-fermentable carbon source medium, the trafficking of copper transport towards mitochondria may lead to a decrease in the cytosolic labile copper pool. If the labile copper pool decreases in the cytoplasm of cells grown in non-fermentable carbon source medium, copper-induced defects in the protein translocation process should be less than those in cells grown in normal SC medium. To examine this hypothesis, we tested the translocation of CPY protein, an example of the secretory pathway, in the wild type and a copper-sensitive histone H3 mutant. According to the hypothesis, we observed only one band of CPY, p2CPY, in the wild-type cells grown in media containing a non-fermentable carbon source (SCEG) in untreated and in 1 mM copper-treated conditions. p2CPY form of CPY is the glycosylated form that resides in the Golgi, indicative of ‘no defect’ in the translocation process. On the other hand, we observed two major bands of CPY protein, p2CPY and ppCPY (cytosolic non-glycosylated form) upon treatment of wild-type cells with 1 mM of copper chloride grown in SC medium (Figure 3A and S3B, S3C, and S3D). Furthermore, only one major band was detected of CPY, ppCPY in the H3 mutant, H3Δ(28–31), upon treatment with 1 mM of copper chloride in SC medium, and two bands, p2CPY (Golgi form) and ppCPY (cytosolic form) in non-fermentable carbon source medium. As an H3 mutant, H3Δ(28–31) contains a higher concentration of labile copper pool, producing only one cytosolic form of CPY, ppCPY, in SC media and vacuole and cytosolic forms in non-fermentable carbon source medium upon copper treatment. Similar results were observed with Gas1, another example of a secretory protein; no defect was observed in the translocation of Gas1 in SCEG medium upon copper treatment (Figure 3B and S3E). These results suggest that an increase in the rate of mitochondrial respiration in non-fermentable carbon source medium suppresses the inhibitory effect of copper on the protein translocation process.

**Figure 3:**
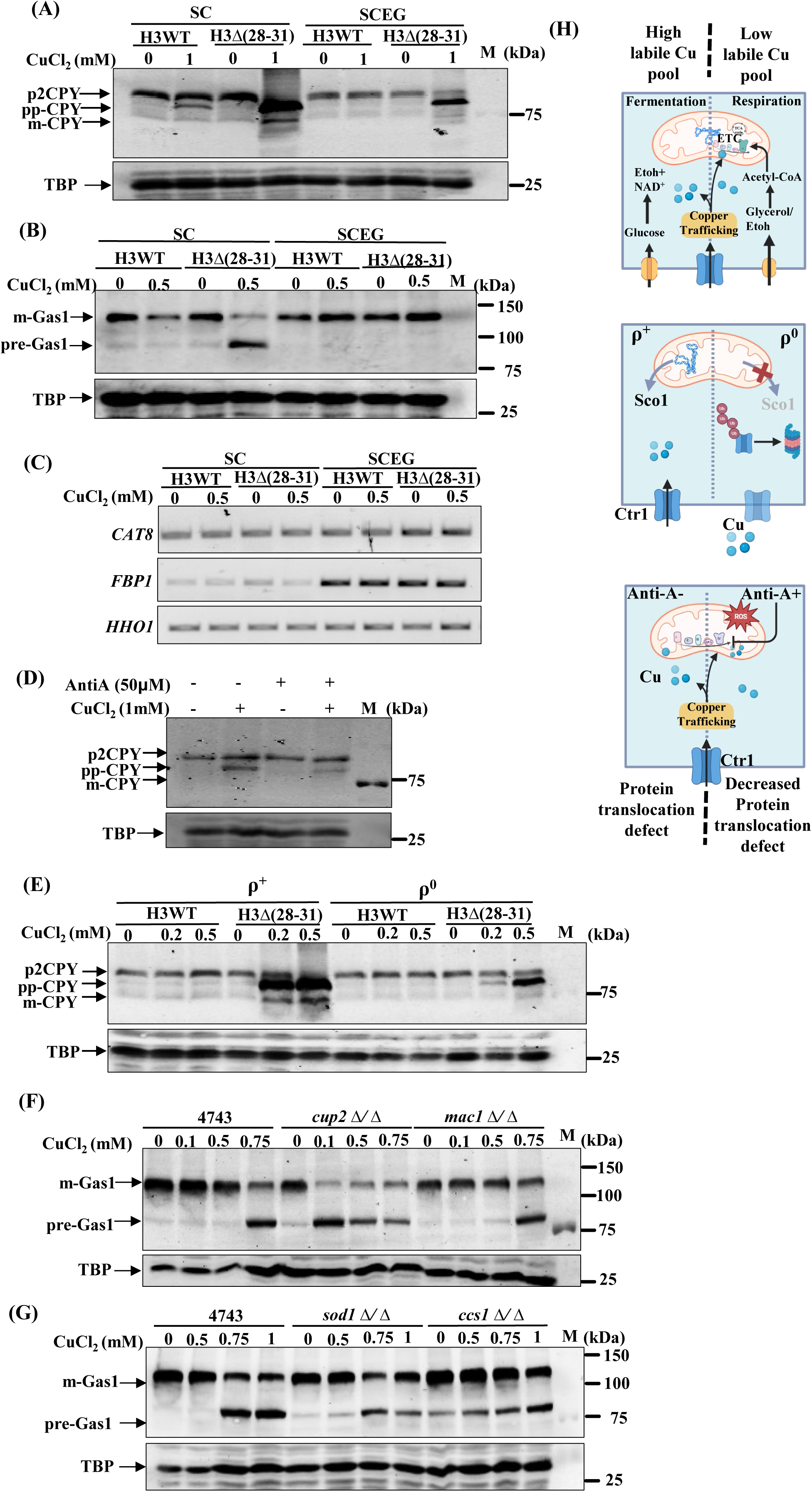
Mitochondrial functions regulate copper homeostasis and protein translocation. A. The overnight grown cultures of wild type, and a histone H3 mutant expressing myc-tagged CPY were seeded at 0.2OD_600_ in SC and SCEG growth medium and cultured till 0.1OD_600_, subsequently grown for 2 hours in the absence or presence of 1.0 mM of copper chloride, and protein translocation was examined by western blotting of CPY using anti-myc primary antibody. B. Exponentially growing cultures of Gas1-GFP transformed wild type, and a histone mutant in SC and SCEG growth medium were further cultured for two hours in the absence or presence of 0.5 mM of copper chloride, and protein translocation was examined by western blotting of Gas1 using anti-GFP primary antibody. C. Expression analysis of gluconeogenic genes through semi-qPCR, CAT8, and FBP1 in cells (wild type and a histone H3 mutant) grown in SC and SCEG growth medium. The expression of the HHO1 gene was taken as a control. D. Overnight grown cultures of myc-tagged CPY expressing wild type cells were seeded at 0.2OD_600_ in SC growth media and grown till 1OD_600_, untreated and treated in the presence of 1.0 mM of copper, 50 µM of Antimycin A, and cotreatment with 1.0 mM of copper plus 50 µM of Antimycin A. Whole cell extracts were subsequently prepared and protein translocation was examined by western blotting of CPY using anti-myc primary antibody. E. Overnight grown cultures of Rho plus and Rho zero of wild type and a histone H3 mutant cells, as indicated, were seeded at 0.2OD_600_ in SC growth media and grown till 1OD_600_, untreated and treated in the presence of 0.2 and 0.5 mM of copper chloride. Extracts were prepared, and protein translocation was examined by western blotting of CPY using an anti-myc primary antibody. F. Overnight grown cultures of Gas1-GFP transformed wild type and *cup2*Δ*/*Δ and *mac1*Δ*/*Δ deleted cells were seeded at 0.2OD_600_ in SC growth media and grown till 1OD_600_, untreated and treated in the presence of copper chloride (0.1, 0.5, and 0.75 mM). Extracts were prepared, and protein translocation was analyzed by western blotting using an anti-GFP primary antibody. G. Overnight grown cultures of Gas1-GFP transformed wild type and sod1Δ*/*Δ and ccs1Δ*/*Δ deleted cells were seeded at 0.2OD_600_ in SC growth media and grown till 1OD_600_, untreated and treated in the presence of copper chloride (0.5, 0.75, and 1.0 mM). Extracts were prepared, and protein translocation of Gas1 was analyzed by western blotting using an anti-GFP primary antibody. H. Hypothetical model to show the role of fermentation and respiration in the regulation of copper trafficking between mitochondria and the cytoplasm (top scheme). Rho^+^ and Rho^-^ model to study the role of mitochondria in copper trafficking (middle scheme). Model to investigate the copper homeostasis under respiration inhibition conditions, treatment with antimycin-A (bottom scheme).

Further to understand the role of mitochondrial respiration in the protein translocation process, we treated the wild-type cells with copper, antimycin A, an inhibitor of respiration, and co-treated them with copper and antimycin A. We observed both bands of CPY in equal intensity, p2CPY (Golgi form) and ppCPY (cytosolic form) upon copper treatment, and in co-treatment, both bands were observed, but with a higher intensity of p2CPY and a lower intensity of ppCPY (Figure 3D and S3I, S3G). A similar effect was observed with the translocation of Gas1 (Figure S3H). The growth defect upon 0.5 mM copper treatment of the H3 copper-sensitive mutant, H3Δ(28–31), which contains a higher concentration of labile copper, was suppressed upon cotreatment with antimycin, which further suggests that copper suppresses the growth by inhibiting the protein translocation process (Figure S3F ). These observations indicate that the transport of copper increases towards mitochondria, which is required for the activity of the antioxidant system to suppress the superoxide radicals produced upon antimycin A treatment. Next, we also examined the role of mitochondria by utilizing rho zero cells (lack of mitochondrial genome) upon copper treatments. We created rho zero of wild type and a copper-sensitive histone H3 mutant, H3Δ(28–31), and were subsequently grown in SC medium in the absence and presence of copper for the analysis of protein translocation by western blotting of CPY (Figure S3J, S3K, S3L, S3M). In copper-treated condition, we detected only one major band of CPY (ppCPY) in H3Δ(28–31) and two bands, p2CPY and ppCPY, in rho zero of H3Δ(28–31) mutant (Figure 3E and S3N). The copper supplementation partially supports the growth of rho zero cells created in the H3Δ(28–31) mutant, probably by suppressing the mitochondrial oxidative stress. Western blotting analysis with cells treated with copper indicates suppression of translocation defect, two forms of CPY, p2CPY (Golgi) and ppCPY (cytosolic) in the rho zero cells created in H3Δ(28–31) mutant in comparison to normal H3Δ(28–31) mutant wherein only one cytosolic form of CPY, ppCPY was detected. Since in this experiment we used only up to 0.5 mM of copper, we did not see any effect on CPY translocation in wild-type cells as well as in rho zero mutants. To further understand the sub-cellular distribution of copper, we utilized a few of the gene deletion mutants, mainly of CUP2/ACE1 (needed for Cup1 expression), SOD1 (suppresses superoxide radicals), CCS1 (encodes a copper chaperone for Sod1), and MAC1 (required for expression of copper transporter). We hypothesized that in the absence of proteins encoded by *CUP2, SOD1, CCS1*, and *MAC1*, the cellular homeostasis of copper will be perturbed, which may or may not affect the growth of mutant cells upon copper treatment. Interestingly, we observed slow growth in *sod1*Δ*/*Δ*, ccs1*Δ*/*Δ and *cup2*Δ*/*Δ mutants upon copper treatment and the growth of *mac1*Δ*/*Δ mutant was not affected (Figure S3O). In the absence of Cup1 (copper metallothionein) in the *cup2*Δ*/*Δ cells, the concentration of labile copper in the cells may increase, which can inhibit the protein translocation process. Indeed, in *cup2*Δ/Δ cells, we observed significantly higher defects in the protein translocation process than wild-type cells (Figure 3F, S3P and S3R). A similar effect on the protein translocation process was observed in *ccs1*Δ*/*Δ cells upon copper treatment (Figure 3G and S3Q, S3S). Since copper is required for the activity of the Sod1 enzyme present in the cytoplasm and in the mitochondria, we tested the growth and the protein translocation assay in *sod1*Δ*/*Δ cells and compared them with the wild-type cells. The growth of *sod1*Δ*/*Δ cells was found to be much better upon supplementation of 1 mM of copper, but growth was reduced at 2 mM of copper. However, the translocation defect in the presence of copper in *sod1*Δ*/*Δ cells was found to be the same as that of wild-type cells. As Sod1 is present in the cytoplasm and mitochondria, it is quite possible that in the absence of Sod1, the labile copper concentration between the cytoplasm and mitochondria may remain unchanged; hence, the effect of copper on the protein translocation process in *sod1*Δ*/*Δ and wild-type cells remains the same. Why *mac1*Δ*/*Δ cells did not show any effect on the protein translocation process upon copper treatment needs to be explored further. Further, we also observed defects in the translocation process (premature band of Gas1) in the *ccs1*Δ*/*Δ mutant in copper-untreated conditions (Figure 3G, S3Q), requires further investigation. Taken together, the above results suggest that copper trafficking between the cytoplasm and mitochondria controls the translocation of secretory proteins.

### Copper suppresses the growth of fis1 and mrs3 deleted cells but inhibit the protein translocation process only in fis1 deleted cells

So far, our experimental observations suggest that mitochondrial activities modulate the protein translocation process by regulating sub-cellular distribution of copper. Further, we hypothesized that mitochondrial dynamics, by the process of fission and fusion, may regulate the copper trafficking between mitochondria and cytoplasm and therefore protein translocation (Figure 4G). To identify the role of mitochondrial dynamics in copper homeostasis and protein translocation process, we screened the mutants defective in mitochondrial respiration, fission, fusion, and ERMES (Figure S4A) by testing their growth upon copper treatment and in co-treatment with copper plus iron (Figure S4B). Interestingly, we observed fis1 and mrs3 gene deletion mutants to be copper sensitive. Gene FIS1 encodes a protein essential for mitochondrial fission and has been shown to inhibit programmed cell death (38). The *MRS3* gene encodes a protein required for iron transport across the mitochondrial membrane (39). Similarly, we also detected a few of the gene deletion mutants to be copper resistant but at higher concentrations (data not shown), such as *mmm1*Δ*/*Δ*, mdm12*Δ*/*Δ, and *mgm1*Δ*/*Δ. The growth of several other gene deletion mutants, *mdv1*Δ*/*Δ*, caf4*Δ*/*Δ*, sdh6*Δ*/*Δ*, fzo1*Δ*/*Δ*, cit1*Δ*/*Δ, etc., was found to be the same as wild-type cells in the presence of copper chloride. Genes, *MMM1*, *MDM12*, and *MGM1*, encode proteins required to maintain mitochondrial morphology through the fusion process, and *CIT1* encodes citrate synthase. Next, we examined the protein translocation process upon copper treatment in gene deletion mutants, which are copper sensitive (fis1 and mrs3), and a few other gene deletion mutants were utilized as controls (*caf4*Δ*/*Δ*, mdv1*Δ*/*Δ*, fzo1*Δ*/*Δ*, cit1*Δ*/*Δ*, sdh6*Δ*/*Δ*, mgm1*Δ*/*Δ*, mmm1*Δ*/*Δ, and *mdm12*Δ*/*Δ). To our surprise, we found defect in protein translocation process only in the *fis1*Δ*/*Δ although both, *fis1*Δ*/*Δ and *mrs3*Δ*/*Δ mutants are hypersensitive to copper chloride. The intensity of premature (cytosolic) and mature (endoplasmic reticulum) forms of Gas1 in *mrs3*Δ*/*Δ mutants was found to be the same as wild-type cells upon copper treatment. However, we observed slightly more defects in protein translocation upon copper treatment only in the *fis1*Δ*/*Δ, requires further investigation. In contrast, mutants that showed slight resistance at higher concentrations of copper (*mmm1*Δ*/*Δ*, mdm12*Δ*/*Δ, and *mgm1*Δ*/*Δ) and *fzo1*Δ*/*Δ showed less defect in the protein translocation process than the wild-type cells upon copper treatment (Figure 4A-F and S4C, S4D, S4E, S4F, S4G, S4H, and S4I). Out of all these mutants, the inhibitory effect of copper on the protein translocation process in the *mgm1*Δ*/*Δ mutant was found to be almost nil in comparison to wild-type cells (Figure 4C). Since *MGM1* is required for the fusion process of mitochondria, in its absence it the defect in mitochondrial fusion may increase the fission process, which in turn may increase the trafficking of copper towards mitochondria (40). As the cellular pool of metabolites (concentration, composition, and sub-cellular distribution) may vary among these gene deletion mutants, which in turn may affect the sub-cellular trafficking of copper and other metals to different extents, leading to differential effects on the protein translocation process upon copper treatment. Alternatively, as *fis1*Δ*/*Δ and *mrs3*Δ*/*Δ gene deletion mutants are slightly resistant to iron, it is quite possible that stoichiometric imbalance between these metals may be responsible for their sensitivity to copper. To examine this notion, we tested the growth of *fis1*Δ*/*Δ and *mrs3*Δ*/*Δ mutant cells upon co-treatment (copper and iron) and found complete rescue of growth (Figure S4B). In addition, studies indicate that the deletion of *fis1* or copper exposure causes cell death due to accumulation of mitochondrial oxidative stress. Further investigations are required to identify the factors responsible for the copper sensitivity of *mrs3*Δ*/*Δ and *fis1*Δ*/*Δ gene deletion mutants. Altogether, the above results indicate that copper suppresses the growth of cells exhibiting defects in mitochondrial fission and iron transport across the mitochondrial membrane.

**Figure 4:**
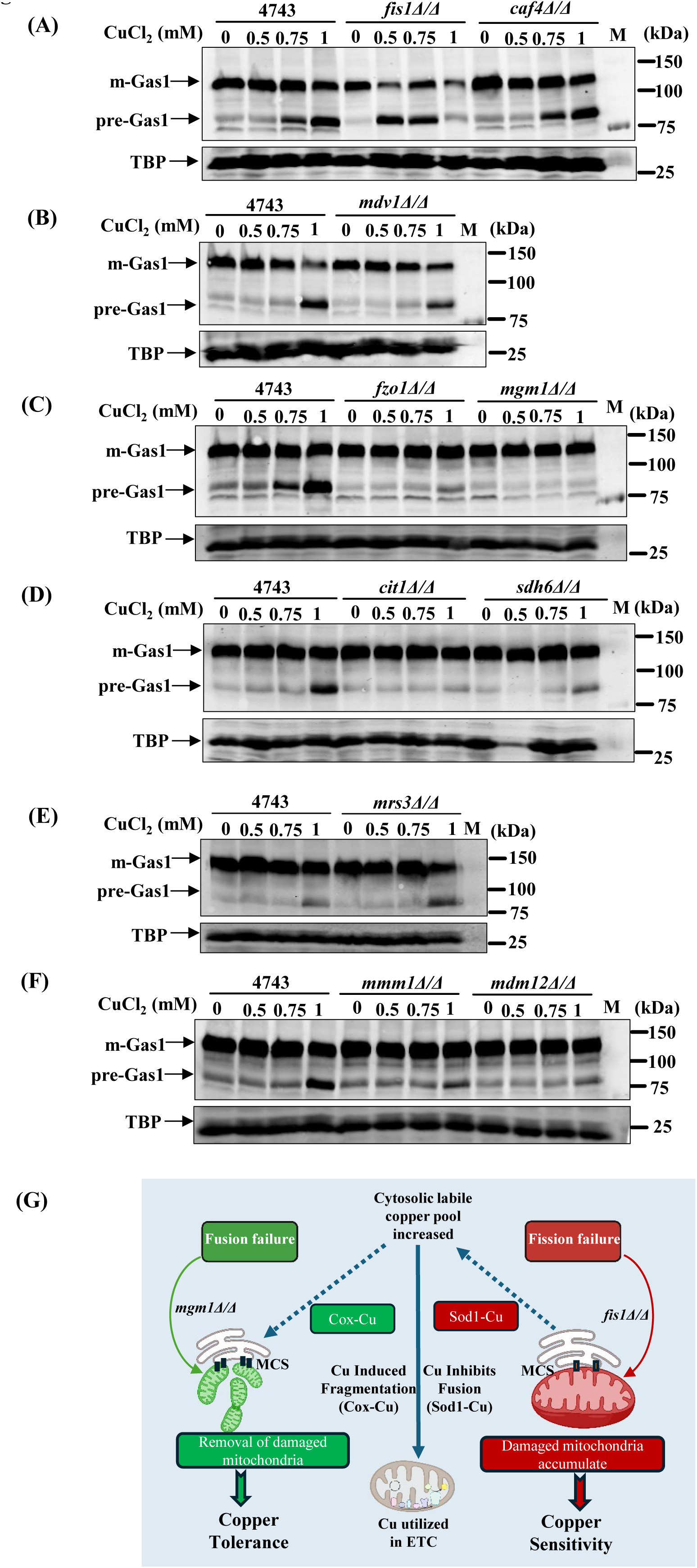
Copper inhibits the growth of *fis1*Δ*/*Δ and *mrs3*Δ*/*Δ mutants; however, only *fis1*Δ/Δ shows defect in the protein translocation. Deletion mutants of mitochondrial respiration, fission, fusion, and ERMES were screened in the presence of copper chloride. The growth of the only two deletion mutants, *fis1*Δ*/*Δ and *mrs3*Δ*/*Δ, was severely reduced upon copper chloride treatments; growth data are shown in the supplementary. The process of protein translocation was tested by western blotting of Gas1 in Gas1-GFP transformed wild type and the deletion mutants. A. *fis1*Δ*/*Δ and *caf4*Δ*/*Δ. B. *mdv1*Δ*/*Δ. C. *fzo1*Δ*/*Δ and *mgm1*Δ*/*Δ. D. *cit1*Δ*/*Δ and *sdh6*Δ*/*Δ. E. *mrs3*Δ*/*Δ. F. *mmm1*Δ*/*Δ and *mdm12*Δ*/*Δ. Overnight grown cultures of wild type and mutants were seeded at 0.2OD_600_ in SC growth media and grown till 1OD_600_, untreated and treated in the presence of increasing concentrations of copper (0.5, 0.75, and 1.0 mM). Extracts were prepared, and western blotting of Gas1 was conducted to assess the protein translocation process. G. Hypothetical scheme to investigate the role of mitochondrial fission and fusion in cellular copper metabolism. Probably the balance between fission and fusion processes regulates the sub-cellular trafficking and labile copper concentration.

### The autophagy and vacuolar activities regulate copper homeostasis and the protein translocation process

The evolutionary conserved metabolic pathways, including autophagy, are crucial to regulate a variety of biological functions, including protein trafficking, organelle homeostasis, and respiratory growth of yeast cells (27). For example, the autophagy process is an intracellular degradation machinery required to maintain the concentration of nutrients during starvation and other pathophysiological conditions (41). Metabolic pathways such as glycolysis, the TCA cycle, and oxidative phosphorylation produce energy. On the other hand, the autophagy process provides raw material for energy production by degradation of cytoplasmic components through autophagosomes formation and fusing them with the vacuoles during starvation conditions (42). As autophagy is required to regulate the concentration of nutrients essential for the growth of cells, it may also impact the metal homeostasis because the products of the autophagy process, such as certain amino acids, can bind with the metals. A few studies suggest that the metabolism of copper, zinc, and sulphur modulates the process of autophagy. However, our understanding of the role of autophagy in copper homeostasis-mediated translocation of proteins is not complete. There are several autophagy-related (Atg) proteins that are required for autophagosome formation (43,44). The process of autophagy regulates the homeostasis (concentration and composition) of the amino acid pool. However, the activation or inhibition of the autophagy process may lead to drastic changes in the cellular pool of amino acids. We hypothesize that the process of autophagy may play a crucial role in regulating the protein translocation process by modulating the amino acid pool-mediated alteration in copper homeostasis (Figure 5F). To examine the hypothesis, we first tested the growth of more than 40 different gene deletion mutants of the autophagy process (ATGs) in the presence of copper to identify the factors of the autophagy process that may be involved in the regulation of copper homeostasis. We found one of the Atg deletion mutants (*atg6*Δ*/*Δ, also known as *vps30*Δ*/*Δ) hypersensitive to copper (Figure S5A). We also tested the growth of one important gene deletion mutant, *vps38*Δ*/*Δ (part of the Atg6 complex), in the presence of copper. *VPS38* encodes a vacuole protein sorting factor that is involved in the autophagy process in yeast. Interestingly, we observed the *vps38*Δ*/*Δ mutant to be copper sensitive, the same as *vps30*Δ*/*Δ. We also found a few other Atg gene deletion mutants slightly resistant to copper. For example, the growth of the *atg15*Δ*/*Δ gene deletion mutant (slow growing) was found to be slightly resistant or better growing in copper treatment conditions in comparison to wild-type cells. These observations suggest that certain ATG proteins play important roles in copper homeostasis by modulating the process of autophagy. Further investigations are in progress to identify the mechanism.

**Figure 5:**
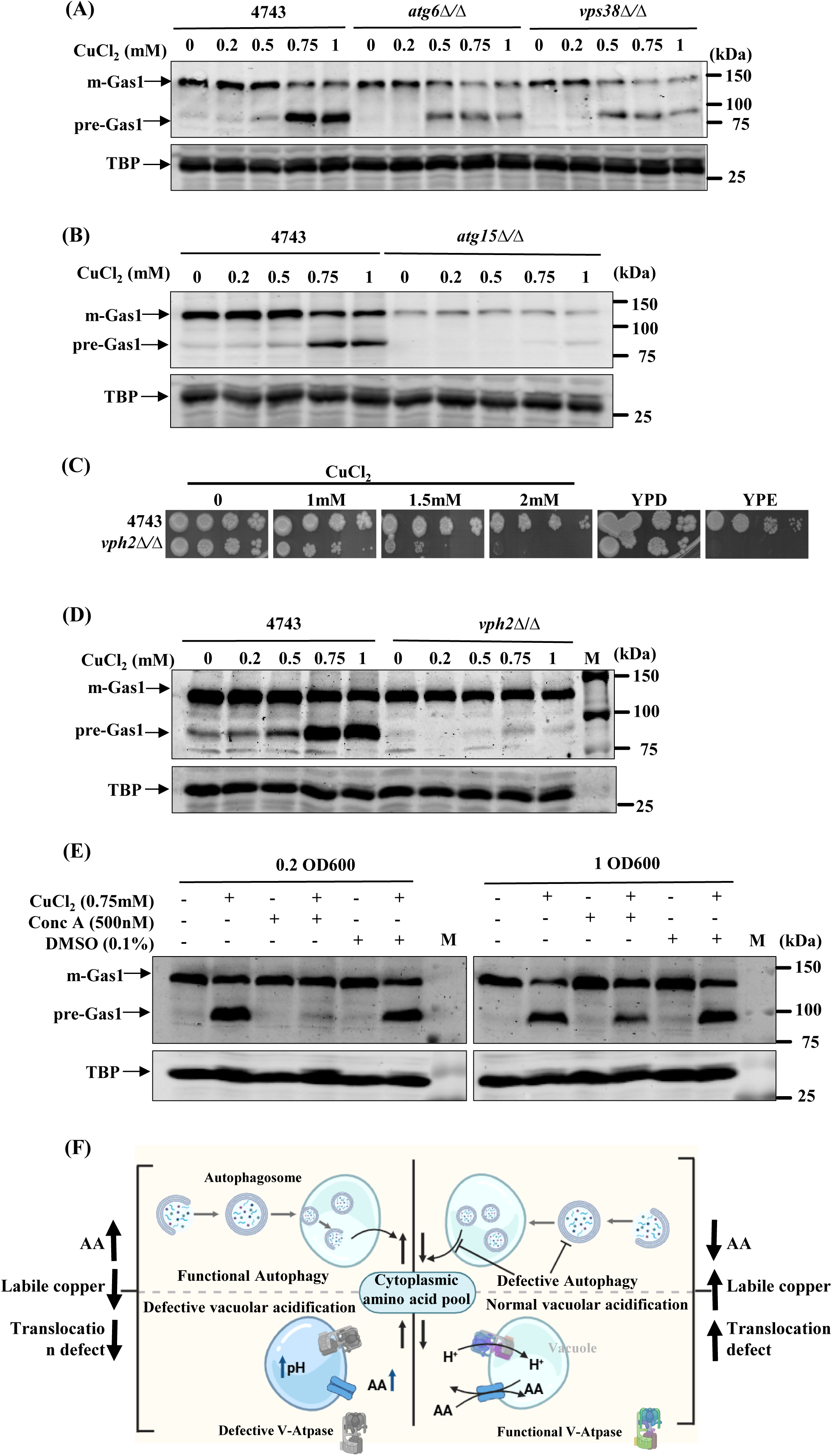
Autophagy and vacuole functions play key roles in regulating copper homeostasis and protein translocation. The growth of gene deletion mutants of autophagy (*atg*Δ*/*Δ) and a deletion mutant of vacuole acidification gene, vph2, was tested in the presence of copper chloride; the data are presented in the supplementary file. Protein translocation was examined in a few of the gene deletion mutants transformed with Gas1-GFP by western blotting of Gas1 using anti-GFP primary antibody. Overnight grown cultures of wild type and respective gene deletion mutants were seeded at 0.2OD_600_ in SC growth media and grown till 1OD_600_, untreated and treated in the presence of copper chloride, and extracts were prepared for the western blotting. A. *atg6*Δ*/*Δ and *vps38*Δ*/*Δ. B. *atg15*Δ*/*Δ. C. Growth of the *vph2*Δ*/*Δ deletion mutant was examined through spot assay in the absence and presence of copper (1.0, 1.5, and 2.0 mM) and compared with wild type. The growth of *vph2*Δ*/*Δ mutant was also assessed in YPD and YPE medium in the absence of copper treatment. D. Analysis of protein translocation process in Gas1-GFP transformed *vph2*Δ*/*Δ deletion mutant and compared with wild type cells. Exponentially growing cells were grown further in SC growth medium in the absence and presence of copper chloride (0.2, 0.5, 0.75, and 1.0 mM) for 2 hours, and western blotting of Gas1 was conducted using anti-GFP primary antibody. E. Analysis of protein translocation process in Gas1-GFP transformed wild-type cells in two different treatment conditions. First, overnight grown wild-type cells were seeded at 0.2OD_600_ in SC growth medium and grown until 1.0OD_600_. Subsequently they were grown for 2 hours upon treatments; 0.75 mM of copper, 500 nanomolar of concanamycin A, and co-treated (0.75 mM copper plus 500 nanomolar Concanamycin A). Second, overnight grown wild-type cells were seeded at 0.2OD_600_ in SC growth medium and 500 nanomolar of concanamycin A was added and grown until 1.0OD_600_. Subsequently 0.75 mM of copper was added and grown for 2 hours. Extracts were prepared, and western blotting of Gas1 was conducted using an anti-GFP primary antibody. F. Model to study the role of autophagy and vacuole acidification in copper homeostasis and protein translocation.

Next, we tested the role of some of the Atg and *vps38*Δ*/*Δ gene deletion mutants in the Sec61-mediated protein translocation process. The gene deletions (*atg6*Δ*/*Δ*, atg15*Δ*/*Δ*, and vps38*Δ*/*Δ) and wild-type cells were grown in the absence and presence of copper. The protein translocation process in these strains was tested by western blotting of Gas1. We detected more translocation defects (appearance of pre-Gas1) in copper-sensitive atg6Δ and *vps38*Δ*/*Δ deletion mutants in comparison to wild-type cells upon 0.5 mM copper treatment. On the other hand, fewer translocation defect (appearance of only mGas1) was detected in copper-resistant, *atg15*Δ*/*Δ mutants upon copper treatment. As *atg15*Δ*/*Δ mutant is slightly copper resistant, whereas *vps38*Δ*/*Δ is highly copper sensitive to copper, we believe that labile copper pool may be higher in *vps38*Δ*/*Δ mutant leading to more defect in translocation process and lower labile copper pool in *atg15*Δ*/*Δ correlates with less defect in protein translocation process (Figure 5A, 5B and S5B, S5E and S5F). The less effect on protein translocation process in *atg15*Δ*/*Δ upon copper treatment can also be due to degradation of lipid droplets which has been shown by a published study (45). The above observations indicate that Atg6, Atg15, and Vps38 are involved in the regulation of the protein translocation process, probably via regulating the metabolism of copper and amino acids. Since amino acid metabolism is mainly regulated by the activities in the vacuole, to gain better insight, we tested the growth of a *vph2*Δ*/*Δ deletion mutant in the presence of copper. The vph2 protein maintains the acidic environment inside the vacuole, which is essential to regulate the functions of vacuolar proteases and amino acid transporters localized on the vacuole membrane (46,47). We find the *vph2*Δ*/*Δ mutant to be copper sensitive, but it does not show any defect in the protein translocation process upon copper treatment in comparison to wild-type cells. Perhaps the alteration in pH homeostasis in *vph2*Δ*/*Δ mutant affects copper metabolism or results in trafficking of copper towards mitochondria, hence not available to inhibit the protein translocation process (Figure 5C, 5D, and S5C, S5G). Furthermore, it is also possible that in the absence of V-ATPase protein, Vph2 may lead to a decrease in the cytosolic pH because V-ATPase maintains proper pH balance within the cytoplasm and sub-cellular compartments. In the absence of V-ATPase, it may be difficult to regulate the proton (H^+)^ levels, which can result in pH imbalance and disruption in copper metabolism. Alternatively, the total amino acid pool increases in the cytoplasm in the absence of Vph2, leading to a decrease in labile copper concentration. In addition, a study suggest that cupric copper (Cu^2+^) can inhibit the vacuole fusion and V-ATPase function (48). To gain more insight, we also treated the cells with Concanamycin A, a V-ATPase-specific inhibitor, and found that inhibitor treatment suppresses the copper-mediated protein translocation defect, like that of the *vph2*Δ*/*Δ mutant (Figure 5E and S5D, S5H). These observations indicate that the acidic environment within the vacuoles plays an essential role in maintaining copper homeostasis.

### Lipid droplets are essential for the regulation of copper homeostasis

Lipid droplets are highly dynamic membrane-enclosed, evolutionarily conserved organelles that majorly store triacylglycerols (TAG), steryl esters (SE), and proteins (49). The large number of studies suggests that homeostasis of lipid droplets is required to regulate a variety of biological functions such as host-pathogen interactions, development, protein homeostasis, and signalling pathways (50). The metabolic states, number, size, and composition (lipids and proteins) of lipid droplets change under certain pathophysiological and stress conditions such as starvation and infections. The dynamic nature of lipid droplets may be essential for survival under different stress conditions. To maintain lipid homeostasis, lipid droplets fuse with vacuoles for their degradation by the process of autophagy. The cell organelles, such as peroxisomes, vacuoles, mitochondria, the endoplasmic reticulum, and lipid droplets, are involved in the process of lipid metabolism. In addition, the biogenesis of lipid droplets is also functionally linked with the accumulation of copper in yeast cells (51). However, our understanding of the lipid droplets in copper homeostasis and the protein translocation process is not complete. We hypothesize that a defect in lipid droplets may adversely affect the autophagy and organelle functions, leading to disruption in copper metabolism (Figure 6B, lower panel). Since lipid droplets regulate a variety of biological functions, including energy metabolism and membrane homeostasis, disruption in the homeostasis of lipid droplets results in pathological conditions. There are four major genes in yeast, *DGA1*, *LRO1*, *ARE1,* and *ARE2,* that encode enzymes for the synthesis of neutral lipids. *DGA1* encodes diacylglycerol acyltransferase 1, *LRO1* encodes lipid acyl transferase, and *ARE1* and *ARE2* encode acyl-CoA:sterol acyltransferases, which are involved in sterol ester synthesis. Cells in the absence of all four of these genes completely lack lipid droplets. To better understand the significance of lipid droplets in copper homeostasis, we created single, double, triple, and quadruple deletion mutants of these four genes and tested their growth on nonfermentable carbon source media and compared them with the normal SC growth media in the absence and presence of copper (Figure S6A). Growth of the following double and triple deletion mutants: *are2*Δ*dga1*Δ, *are1*Δ*are2*Δ*dga1*Δ, *are1*Δ*lro1*Δ*dga1*Δ, and *are2*Δ*lro1*Δ*dga1*Δ was reduced in the presence of copper in SC media. However, copper did not affect the growth of the quadruple mutant (H1246) lacking all four genes. On the other hand, the growth of *are1*Δ*lro1*Δ double deletion and *are1*Δ*are2*Δ*lro1*Δ triple deletion mutants was severely reduced in non-fermentable carbon source medium (YPEG), suggesting that the lipid droplets play an essential role in copper homeostasis by regulating mitochondrial functions. These results indicate that the composition of lipid droplets modulates the copper-induced cell death. Lipid droplets serve as storage sites for neutral lipids. In addition, peroxisomes are also necessary for fatty acid oxidation and lipid metabolism. The lipid droplets and peroxisome organelles are frequently present in proximity, indicating that their cellular functions may be co-regulated. As lipid droplets (LDs) and peroxisomes are functionally linked organelles and both are involved in lipid metabolism (52–54), we went ahead to test the growth of a few important gene deletion mutants of peroxisome (fatty acid oxidation, matrix and membrane proteins, membrane receptor, and biogenesis) in copper. Interestingly, a few of the gene deletion mutants of peroxisome were found to be copper sensitive (*rtn1*Δ*/*Δ) and a few resistant (*faa2*Δ*/*Δ, *pex31*Δ*/*Δ, and *pex13*Δ*/*Δ), indicating that the proteins encoded by these genes play a crucial role in the regulation of cellular copper homeostasis (Figure S6F, S6G). To gain more insight about the role of lipid droplets and peroxisomes in copper homeostasis, we examined the protein translocation process in a few of the copper-sensitive and resistant gene deletion mutants of lipid droplets and peroxisomes. First, we examined the protein translocation process in the lipid droplet mutants upon copper treatment by western blotting of Gas1. Wild type, SCY62 and the gene deletion mutants of lipid droplets, *are1*Δ*, are1*Δ*lro1*Δ*, are2*Δ*dga1*Δ and *are2*Δ*lro1*Δ*dga1*Δ were grown in SC growth medium and treated with different concentrations of copper chloride for 2 hours. We observed higher translocation defects in the *are1*Δ mutant upon copper treatment and much fewer defects in *are1*Δ*lro1*Δ*, are2*Δ*/dga1*Δ, and *are2*Δ*/lro1*Δ*/dga1*Δ in comparison to wild-type cells (Figure 6A, 6B, and S6B, S6C, S6D, and S6E). These observations indicate that lipid droplets play important roles in maintaining copper homeostasis. Next, as the *rtn1*Δ*/*Δ mutant is copper sensitive, and *faa2*Δ*/*Δ*, pex31*Δ*/*Δ, and *pex13*Δ*/*Δ mutants showed resistance to copper chloride treatment, we decided to examine the protein translocation process in these cells by taking the example of Gas1. We observed about 15-20 percent fewer defects in the protein translocation process upon treatment, but only in the *rtn1*Δ*/*Δ mutant in comparison to wild-type cells (Figure 6C, 6D, and S6H, S6I, S6J, and S6K). Since Rtn1 plays a role in maintaining ER membrane structure (52), it is quite possible that alteration in the ER surface area in the absence of Rtn1 may affect the translocation of proteins conducted through the Sec61 channel. Taken together, these results suggest that the metabolism of lipid droplets and peroxisome activities can regulate copper homeostasis by modulating the mitochondrial and ER functions.

**Figure 6:**
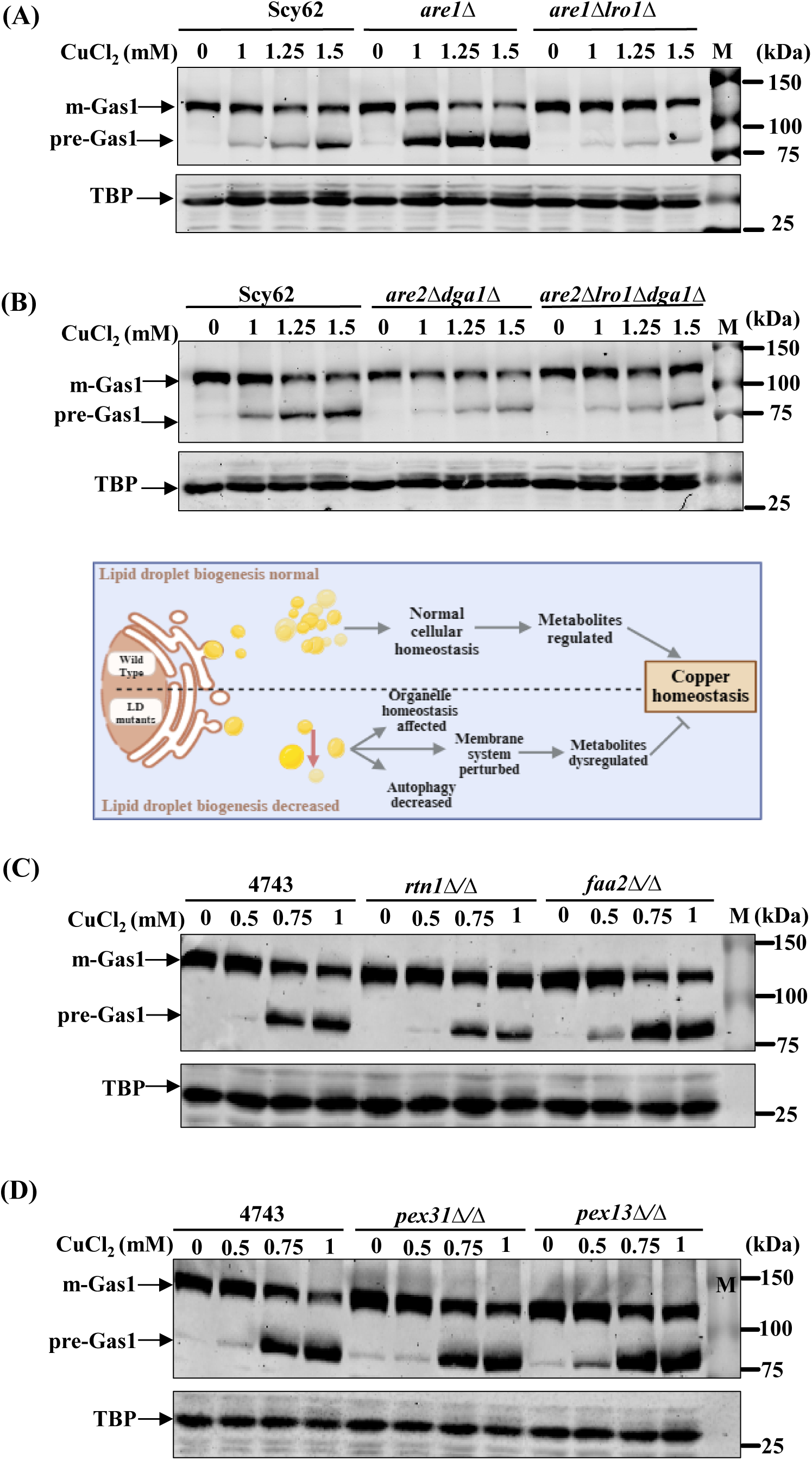
Lipid droplets are key regulators of copper homeostasis. The growth of gene deletion mutants of lipid droplets and peroxisome biogenesis was tested upon copper chloride treatments; data are available in the supplementary file. The protein translocation process was examined upon copper treatments by western blotting of Gas1 using anti-GFP primary antibody in gene deletion mutants of lipid droplets (*are1*Δ*, are1*Δ*lro1*Δ*, are2*Δ*dga1*Δ, and *are2*Δ*lro1*Δ*dga1*Δ) and of peroxisome (*rtn1*Δ*/*Δ*, faa2*Δ*/*Δ*, pex31*Δ*/*Δ, and *pex13*Δ*/*Δ) along with the wild type. Overnight grown cultures of Gas1-GFP transformed wild type, and mutants were seeded at 0.2OD_600_ in SC growth media and grown till 1OD_600_, untreated and treated in the presence of different concentrations of copper chloride as indicated, and protein extracts were prepared for analysis of Gas1 by western blotting. A. Western blotting analysis of Gas1 in wild type, SCY62, and deletion mutants, *are1*Δ and *are1*Δ*lro1*Δ. B. Western blotting analysis of Gas1 in wild type, SCY62 and mutants, *are2*Δ*dga1*Δ, *are2*Δ*lro1*Δ*dga1*Δ. Below this western image, a hypothetical model is given to understand the role of lipid droplets in copper homeostasis. C. Western blotting analysis of Gas1 in wild type, BY4743, and mutants, *rtn1*Δ*/*Δ and *faa2*Δ*/*Δ. D. Western blotting analysis of Gas1 in wild type, BY4743, and mutants, *pex31*Δ*/*Δ and *pex13*Δ*/*Δ.

### Redox and pH homeostasis are essential to protect from copper-mediated protein translocation defects

Redox mechanisms, consisting of oxidation and reduction reactions, which play important functions to regulate cellular processes, including metabolism, imbalance can cause a variety of pathophysiological diseases in humans (55). Living organisms produce reactive oxygen species (ROS) as byproducts during metabolic reactions such as the mitochondrial respiratory chain. In addition, certain stress conditions can also generate a large amount of ROS, which can damage the biomolecules. However, living organisms have evolved with systems that allow them to tolerate the ROS exposure. For example, the antioxidant systems include superoxide dismutase (SOD), catalase (CAT), glutathione peroxidase (GPx), and thioredoxin (Trx), in addition to several other antioxidants that function in a non-enzymatic manner to protect from oxidative damage (56). Understanding about the role of redox mechanisms in metal homeostasis, especially of copper and the cytosolic secretory pathway of proteins, is not complete. We hypothesize that redox imbalance may affect the solubility of copper and other metals inside the cellular environment (Figure 7D). Therefore, to understand the role of redox systems in the copper homeostasis and Sec61 mediated translocation of proteins, we employed majorly three experimental approaches; first, tested the effect of copper chloride on the growth of wild type cells at increasing pH and second, examined the growth of the gene deletion mutants of redox enzymes and alkaline pH sensitive mutants of Slt2 MAPK pathway upon copper treatment (57). To understand the impact of copper on the growth, we grew wild-type and a few copper-sensitive histone mutant cells in SC growth medium and spotted them on the SC agar solid growth medium at increasing pH in the absence and presence of copper. The suppression of the growth phenotype of wild-type cells upon copper treatment was abolished at higher pH above 6.5, which is because the solubility of copper decreases at the alkaline pH (Figure S7C). Next, we examined the effect of copper on the protein translocation process at increasing pH by utilizing Gas1-transformed wild-type cells. The wild-type cells were treated with copper (1 and 1.5 mM) for 2 hours at increasing pH (4.8, 5.5, 6.25, 6.85, and 7.5), and it was observed that copper inhibits the protein translocation process only in the acidic pH range. Above pH 5.5, copper treatment did not generate the immature form of the Gas1 protein, which is indicative of no defect in the protein translocation process (Figure 7B and S7D, S7H). Subsequently, we tested the growth of gene deletion mutants of redox systems and tested the protein translocation process in the absence and upon copper chloride treatment (Figure 7A and S7A, S7B, S7G). Analysis of growth suggests that copper slightly suppresses the growth of *taz1*Δ*/*Δ (Figure S7A), which correlates with slightly more defects in the translocation of Gas1 in *taz1*Δ*/*Δ mutant upon copper chloride treatment (Figure 7A and S7B). This observation suggests that the cellular pH balance is required to regulate copper homeostasis. Next, we tested the growth of gene deletion mutants of CWI pathway and found that an alkaline pH sensitive mutant, *bem2*Δ*/*Δ, is hypersensitive sensitive to copper chloride treatment at acidic pH (Figure S7E, however, did not show any significant defect in protein translocation process in comparison to wild type cells (Figure 7C and S7F, S7I). Yeast Bem2 is a Rho GTPase-activating protein (RhoGAP) that regulates the actin cytoskeleton and cell morphology. Although not known but we believe that Bem2 can indirectly influence vacuole morphology and acidification by affecting cytoskeleton formation and vesicle trafficking. It is quite possible that in the absence of bem2, a defect in the vacuole may disrupt the cytosolic amino acid pool, preventing copper trafficking towards Sec61, but can cause cell death due to oxidative stress. Alternatively, as many of the amino acids are required to maintain the redox balance (32), a defect in the vacuole may result in redox imbalance due to disruption in the amino acid pool. This observation further confirms that cellular pH can modulate the copper-mediated defects in the protein translocation process.

**Figure 7:**
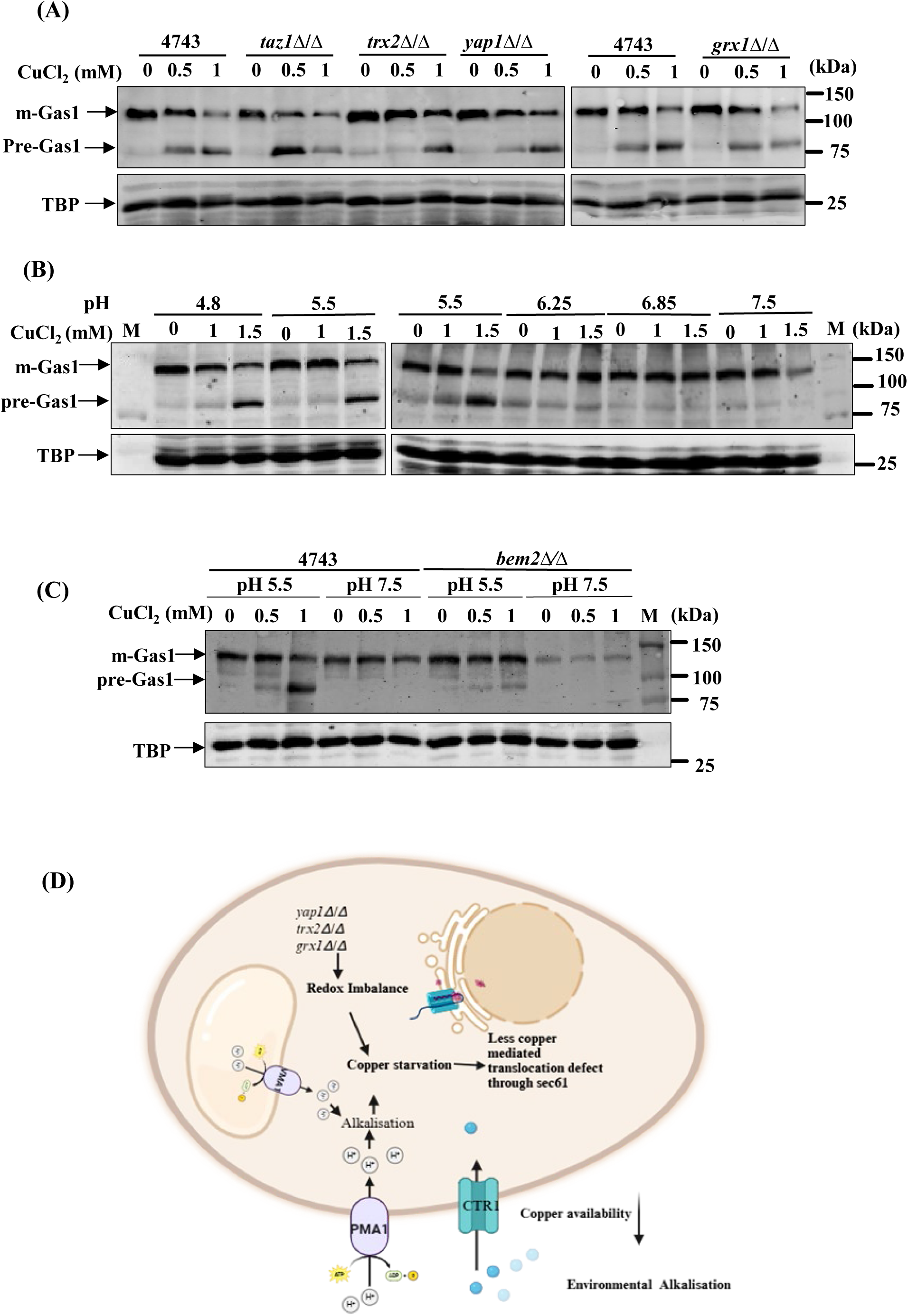
Redox and pH homeostasis regulates copper mediated protein translocation process. A. Overnight grown cultures of Gas1-GFP transformed wild type and gene deletion mutants of redox system (*taz1*Δ/Δ, *trx2*Δ/Δ, *yap1*Δ*/*Δ and *grx1*Δ/Δ) were seeded in SC growth medium at 0.2OD_600_ in SC growth media and grown till 1OD_600_, untreated and treated in the presence of 0.5 and 1.0 mM of copper chloride. Whole cell extracts were prepared, and the protein translocation process was examined by western blotting of Gas1 using an anti-GFP primary antibody. B. Overnight grown culture of a Gas1-GFP transformed wild type strain was seeded at 0.2OD_600_ in SC growth medium at different pH (4.8, 5.5, 6.25, 6.85 and 7.5) and cultured till 1OD_600_, treated and untreated with copper chloride (1.0 and 1.5 mM) for two hours, extracts were prepared and translocation of Gas1 was assessed by western blotting using anti-GFP primary antibody. C. Overnight grown culture of Gas1-GFP transformed wild type and a gene deletion mutant of CWI pathway, *bem2*Δ/Δ was seeded at 0.2OD_600_ in SC growth medium at different pH (5.5 and 7.5) and grown till 1OD_600_, untreated and treated in the presence of copper chloride (0.5 and 1.0 mM) for 2 hours, extracts were prepared and translocation of Gas1 was assessed by western blotting using anti-GFP primary antibody. D. Model scheme to understand the significance of redox homeostasis in copper metabolism, impacting the protein translocation process.

**Figure 8:**
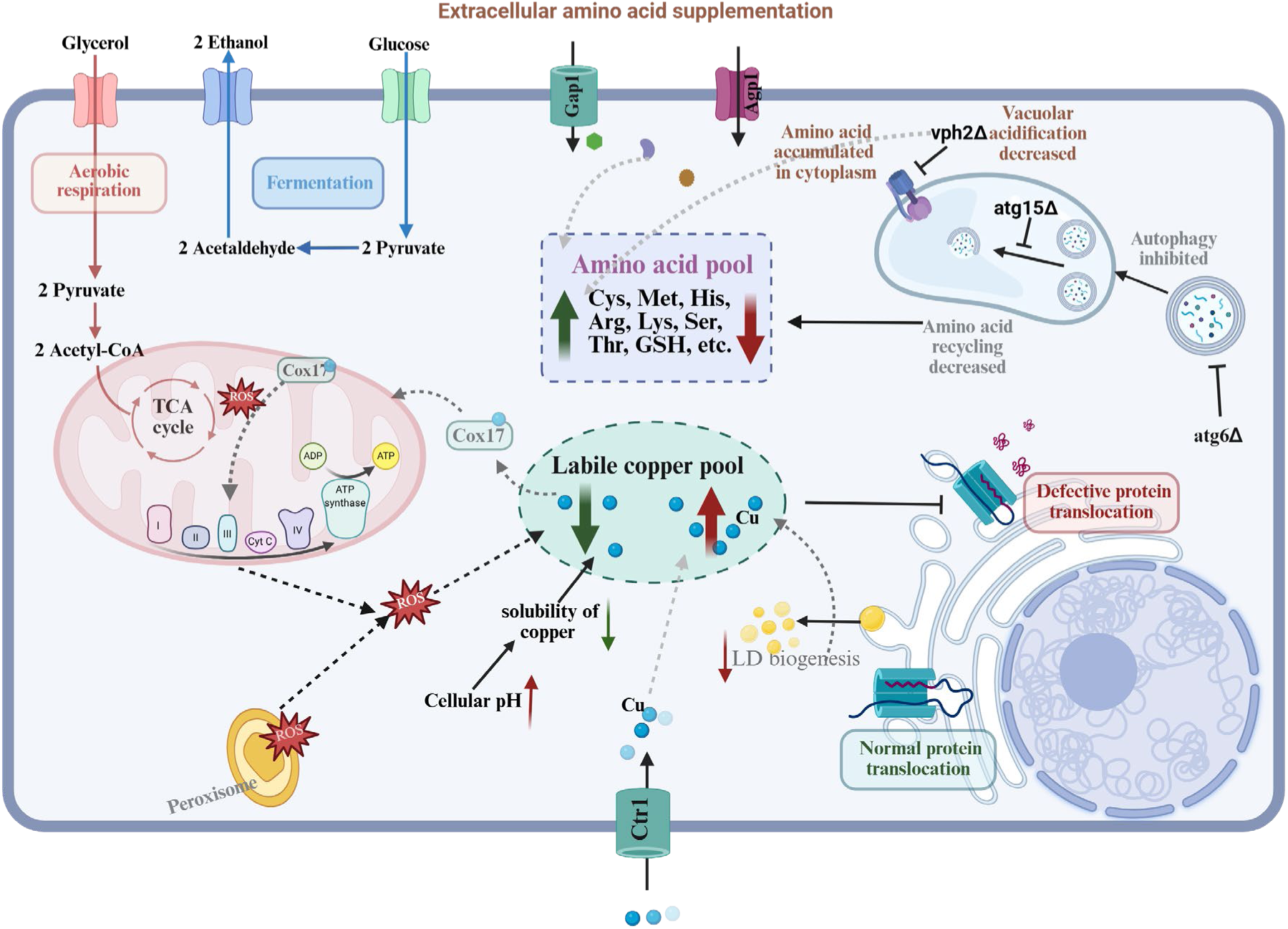
Schematic representation summarising the role of mitochondria, autophagy, vacuole, pH, and lipid droplets in copper homeostasis impacting the process of protein translocation. Trafficking of copper increases towards mitochondria in different conditions, such as growth of wild-type cells in the non-fermentable carbon source medium, and upon inhibition of respiration, in rho zero cells, and in sod1 deleted cells, suppressing the copper-induced defects in the protein translocation process due to a decrease in labile copper concentrations in the cytoplasm. Similarly copper induced defects in protein translocation also decreases in cells grown in YPD rich medium and in SC medium in presence of higher concentration of amino acids such as Cys, His, Met, Arg, Lys, Ser, Thr and GSH. Defects in vacuole acidification in absence of vph2 or upon treatment with concanamycin A causes sensitivity to copper but suppresses the copper induce defects in protein translocation process. Absence of certain genes of autophagy and lipid droplets modulates copper homeostasis and the protein translocation process. Above the normal pH of the growth medium suppresses the copper-induced protein translocation process.

## Discussion

Copper is one of the essential trace elements universally present across living organisms. A variety of external environmental and internal cellular factors can influence copper homeostasis. Many of the metalloenzymes of biological pathways utilize copper as a cofactor. Copper is also utilized in other applications such as pesticides, fungicides, anti-bacterial and anticancer therapies. For example, treatment of triple negative breast cancer (TNBC) patients with a copper chelator, tetrathiomolybdate, has been shown to suppress the metastasis, indicating that copper promotes the proliferation of cancer cells (58). Disruption in copper homeostasis, deficiency or excess concentration is associated with diseases in humans (25). Not only the concentration, the subcellular distribution, and the oxidation states of copper (cupric/cuprous) can also play a crucial role in the regulation of life processes. The cellular concentration of the labile pool of copper is mainly regulated through the membrane transporters, metallothionein, and chaperones. The ratio between cupric or cuprous forms of copper can be impacted by the activity of the copper-specific reductases, cellular redox state, and utilization. It is the cupric form of copper that is utilized by the enzymes to control their activities. As copper is an evolutionarily conserved essential element, it is quite possible that copper regulates the fundamental biological pathways such as mitochondrial respiration, autophagy, immune response, apoptosis, metabolism of proteins and amino acids, MAP kinase signaling pathways, etc (59–61). Like other metals, copper can also bind with a variety of cellular components with different affinities depending upon the chemical nature of binding sites. However, at present, our understanding of the factors regulating cellular homeostasis, as well as the targets of copper, is incomplete. This study was designed to identify the metabolic pathways regulating copper homeostasis because an increase in the labile copper pool can cause toxicity, inhibit the enzymes of the TCA cycle, and the protein translocation process.

Observation of the experiments conducted with gene deletion mutant strains of yeast cells defective in mitochondrial activities, autophagy, amino acid metabolism, vacuolar functions, lipid droplets, peroxisome functions, and redox balance, along with pharmacological inhibition of mitochondrial respiration and vacuole acidification indicates that the metabolic factors are essential for the regulation of copper homeostasis and protein translocation process. Our results indicate that the composition of growth media has a significant impact on the maintenance of the labile copper pool. No defects in the protein translocation process were observed upon copper treatment when cells were grown in rich medium. However, strong inhibition of the protein translocation process was observed upon copper treatment of cells grown in minimal medium.

Given that the amino acids histidine and cysteine are known to strongly chelate copper, we tested whether supplementation of individual amino acids in the growth medium could affect copper-induced defects in the protein translocation process. We observed that supplementation of lysine, arginine, serine, threonine, and glycine amino acids in cells grown in minimal medium suppressed the copper-induced growth repression and defects in the protein translocation process, indicating that such amino acids can modulate the cellular labile copper pool, preventing toxic effects. These results suggest that the cellular amino acid pool is maintained by the nutrients present in the growth medium and the process of autophagy and vacuole sequestration.

Since mitochondria is one of the major copper utilizing cell organelles and provides precursors for biosynthesis of amino acids, we went ahead and tested the impact of copper on protein translocation process in cells grown in non-fermentable carbon source medium, upon inhibition of respiration, in rho zero cells (lacking mitochondrial genome) and deletion mutants of *CUP2*/*ACE1*, *CCS1*, *MAC1* and *SOD1*. We hypothesized that in rho zero cells, the use of copper in the mitochondria may increase to suppress the toxic effects of accumulated free radicals, which means the demand for copper in the mitochondria may increase, leaving less copper in the cytoplasm. It is also known that the degradation of Ctr1 increases in rho zero due to lack of Sco1 which can decrease the copper import (62). Similarly, demand for copper in mitochondria may also increase to suppress the free radicals generated upon inhibition of respiration by antimycin treatment and in non-fermentable carbon source medium, causing a decrease in copper levels in the cytoplasm. Consistent with this hypothesis, copper-mediated defects in protein translocation were significantly reduced in rho zero, cells treated with antimycin, and cells grown on non-fermentable carbon sources. These findings suggest that copper trafficking between the cytoplasm and mitochondria can influence the protein translocation process. In contrast, deletion of *CUP2*/*ACE1*, a transcription factor required for *CUP1* expression, resulted in increased labile copper and significantly greater copper-induced protein translocation defects. However, the deletion of *CCS1*, *MAC1*, and SOD1 did not alter the defect in the translocation process upon copper treatment; it was found to be the same as that in wild-type cells. Probably, copper homeostasis does not change in the absence of CCS1, MAC1, and *SOD1*. Further investigation is needed to understand the role of *CCS1*, *MAC1,* and *SOD1* in copper homeostasis.

Given that mitochondrial function is regulated by fission, fusion, and ERMES, we also tested copper sensitivity and protein translocation in gene deletion mutants defective in these processes. Among several mutants, *mrs3*Δ*/*Δ and *fis1*Δ*/*Δ showed slower growth in the presence of copper, but defect in protein translocation was detected only in *fis1*Δ*/*Δ in comparison to wild-type cells. Since the balance between mitochondrial respiration and the autophagy process regulates the cellular pool of amino acids, we decided to identify the role of autophagy in copper-mediated inhibition of the protein translocation process. We screened approximately 40 *Atg* gene deletion mutants for copper sensitivity. Some mutants (*atg6*Δ*/*Δ*, atg36*Δ*/*Δ*, atg39*Δ*/*Δ, and *vps38*Δ*/*Δ) were copper-sensitive, while others (*atg15*Δ*/*Δ*, atg32*Δ*/*Δ and *atg34*Δ*/*Δ) were copper-resistant. We further went ahead and analyzed the protein translocation process in selective mutants (*atg6*Δ*/*Δ*, atg15*Δ*/*Δ, and *vps38*Δ*/*Δ). Translocation defects were more pronounced in *atg6*Δ*/*Δ and *vps38*Δ*/*Δ mutants and less severe in *atg15*Δ*/*Δ mutants compared to wild-type cells. These findings suggest that autophagy may regulate the impact of copper on protein translocation, likely by modulating the intracellular amino acid pool. As we know, peroxisome, vacuole, mitochondria, endoplasmic reticulum, and lipid droplets regulate the process of lipid metabolism, we wanted to examine the role of lipid droplets in copper homeostasis. Moreover, the biogenesis of lipid droplets has also been shown to be functionally linked with the accumulation of copper in yeast cells. To better understand the role of lipid droplets in copper homeostasis we created and tested the growth of single, double, triple and quadruple gene deletion mutants (Figure S6A) in the presence of copper in the SC media. Surprisingly, while the double and triple mutants exhibited severely reduced growth in the presence of copper, the quadruple mutant (H1246: *are1*Δ*, are2*Δ*, lro1*Δ*, dga1*Δ) did not show any growth defect upon copper treatment. Additionally, the supplementation of zinc or iron did not affect the growth of any of the mutants.

These findings suggest that the composition of lipid droplets modulates copper-induced cytotoxicity. Since the solubility of copper can be impacted by the pH, we examined the effect of copper on the protein translocation process by taking the example of Gas1. Gas1-GFP transformed cells were grown in SC growth medium at different pH levels in the absence and presence of increasing amounts of copper chloride. Interestingly, we observed a significant reduction in defects in protein translocation at pH above 6.25, which is due to the low solubility of copper, as it has already been shown by others. To further understand the role of pH on copper-mediated defects in protein translocation process, we tested the growth of a few alkaline-sensitive mutants of the CWI pathway and found one of the gene deletion mutants, *bem2*Δ*/*Δ, sensitive to copper chloride treatment. Bem2 gene encodes a Rho GTPase-activating protein (RhoGAP), involved in the regulation of actin cytoskeleton and bud formation. Although not known, we believe Bem2 can indirectly influence acidification process inside the vacuole by affecting cytoskeleton formation and vesicle trafficking. This observation further confirms that cellular pH can modulate the copper-mediated defects in the protein translocation process.

In summary, this comprehensive study for the first time provides evidence that mitochondrial activities regulate the sub-cellular trafficking of labile copper impacting Sec61 mediated translocation of proteins process. The experiments upon supplementation of individual amino acids in the growth medium has led to the identification of amino acids suppressing the copper induced toxicity and defects in protein translocation process. Lastly experimental observations upon pharmacological inhibition of mitochondrial respiration and vacuole acidification along with screening of gene deletion mutants of different pathways suggest that autophagy, lipid droplets and redox systems play important roles in the regulation of copper homeostasis and protein translocation process. Taken together, our findings indicate the involvement of metabolic pathways in the regulation of cellular copper homeostasis and the secretory pathway of proteins, with broader applications to understand the diseases of copper metabolism.

## Materials and methods

### Yeast strains, Chemicals, and growth conditions

All the yeast strains and their genotypes used throughout this study are listed in Table S1, and the primers used for confirmation of strains are listed in Table S2. The selected gene deletion mutant strains were validated by genomic PCR using primers for ORFs of respective genes (Figure S8). The yeast strains were grown in SC (synthetic complete), YPD (1% yeast extract, 2% peptone, 2% dextrose), YPE (2% Ethanol), YPG (2% Glycerol), YPEG (1% ethanol, 1% Glycerol), and SC dropout media. SC growth media composed of 0.17 % of YNB without ammonium sulphate, 1.81 g per liter amino acids mix (adenine, arginine, glutamate, glycine, aspartate, isoleucine, lysine, histidine, methionine, phenylalanine, leucine, proline, serine, threonine, tryptophan, tyrosine, uracil, and valine), 0.5 % of ammonium sulphate, and 2% of D-glucose. Yeast nitrogen base without ammonium sulphate was purchased from HI-Media, India. Glucose, ammonium sulphate, and all amino acids were purchased from Merck Sigma Aldrich. The petri dish plates containing SC solid agar growth media were prepared by adding 2% Bacto agar (BD) into the liquid growth media. The liquid cultures of yeast cells were grown at 30°C, 200 rpm in the incubator shaker (Innova 44, New Brunswick Scientific). Copper chloride dihydrate (CuCl_2_·2H2O), Antimycin A and Concanamycin A were purchased from Sigma with cat. No. 307483, A8674 and C9705 respectively.

### Growth assays

The drop test experiments were performed to examine the growth phenotype of the yeast strains, wild type and the mutants of different pathways in untreated and under different treatment conditions. Overnight grown yeast cultures were adjusted to an OD_600_ of 1.0, and ten-fold serial dilutions were prepared in deionised water. The 3 μL of each diluted cultures were spotted onto the solid growth medium, added with 2% glucose or other carbon sources (such as 2% Glycerol, 2% Ethanol and 1% Glycerol plus 1% ethanol), with/without the addition of CuCl_2_, FeSO_4_, and amino acids, as indicated in the figures and legends. Amino acid dropout plates were prepared by eliminating individual amino acids (mentioned in figures) from standard SC growth media. The petri dish plates containing SC plus agar growth media were prepared by supplementing an extra amount of individual amino acids at the concentration of 3, 5, or 10 mM. For the inhibition of respiration, antimycin-A was added at the concentration of 10, 20, 50 µM in the SC growth medium. The plates were incubated at 30°C and the growth of the cells was recorded after 72 hrs using an HP Scanjet scanner.

### Generation of rho/petite cells

Yeast strains lacking mtDNA (rho zero) were generated by treatment of the Histone H3 wild-type and an H3 mutant H3Δ(28–31) strain with Ethidium Bromide (EtBr) as described earlier (63). Briefly, overnight grown primary cultures were seeded at 0.2 OD_600,_ and after adding 10 μg/ml of EtBr, the cells were cultured in YPD growth medium at 30°C while being stirred for around 24 hours. Following a second and third treatment with 10ug/ml EtBr for 24hr, the cells were diluted (1:1000). EtBr-treated and untreated cells were plated on YPD growth medium to create single colonies incubated at 30°C for 2-3 days. The Rho mutants/petite cells were confirmed by streaking them on YPEG growth media containing plates. Ethidium bromide-treated cells, which lack mitochondrial DNA, were unable to grow on YPEG plates.

### Total RNA isolation and Semi-q PCR analysis

Saturated overnight grown cultures of yeast cells were diluted in growth media at 0.2OD_600_, were grown till mid-log phase and treated with CuCl_2_ for 20 min. Untreated and copper chloride treated yeast cultures were harvested, and total RNA was isolated from cell pellets by using a heat/freeze phenol method as described earlier (64). Normalised RNA was used to prepare cDNA using the iScript cDNA Synthesis kit (100R BioRad, cat no. 1708891) according to the manufacturer’s guidelines. The primers used for semi-q polymerase chain reaction analysis are listed in Table S2. The relative expression in fold changes was calculated using the Histone H1 (HHO1) housekeeping gene as control.

### Transformation and tagging

CPY at C-terminal end in the respective yeast strains were myc-tagged using the pFA6a-13myc plasmid. A template for tagging was created using the primers specific for CPY C-terminal tagging and pFA6a-13myc plasmid through two-step PCR. Transformation of respective yeast strain was performed by using the purified PCR product, and positive colonies were selected on SC-Trp plates (65). Briefly, mid-log phase yeast cells were taken to prepare competent cells. The cells at mid log phase were harvested and washed with autoclaved Milli-Q (AMQ) water and subsequently with 1X LioAc-TE buffer and were incubated at 30 °C for 45 min along with plate buffer (50% PEG, 1× LioAc-TE), template, and salmon ss-DNA, followed by incubation at 42 °C for 15 min. This was followed by removing the plate buffer with AMQ water and cells were subsequently plated on the SC-Trp selection plate. Western blotting was performed to confirm protein tagging using anti-Myc antibody. The competent cells of respective strains were also transformed with the pRS415-Gas1-GFP plasmid using the same above-mentioned procedure. Selection media plates (SC-Leu/SC-His) were used to maintain the Gas1-GFP plasmid-transformed cells. All the plasmids used in this study are listed in Table S3.

### Whole cell protein extract preparation

Exponentially growing yeast cultures at 0.8-1.0OD_600_ were treated with and without copper chloride and individual amino acids (Arginine, Lysine, Serine, Glycine, Threonine), Antimycin A, Concanamycin A, and cotreatment with copper chloride for 2 hr. After treatment, cells were harvested at 3000 rpm at 4°C and washed subsequently with water and 20 percent trichloro acetic acid (TCA). Whole cell protein extracts were prepared by following a standard TCA extraction method (12). Briefly, 10 ml of pelleted cell cultures were suspended in 200 µl of 20% TCA, and the suspended mixture was vortexed vigorously in the presence of glass beads to disrupt the cells. The suspension was pelleted at 10000 rpm, 5 min, and TCA was washed with 0.5 M Tris–Cl (pH 7.5). The protein pellet was boiled at 100°C in 0.5 M Tris–Cl (pH 7.5) and 6xSDS dye in a ratio of 3:1 for 10 min. Cell debris was removed by centrifugation (10,000 g, 5 min, 4 °C), and the supernatant was collected for further protein analysis.

### Immunoblotting

Proteins were resolved through SDS-PAGE and were analysed by western blotting according to the standard protocols. Briefly, after SDS-PAGE, proteins were transferred onto the 0.45 μm nitrocellulose membrane (Bio-Rad, 1620115) for 90 min at 4 °C along with a pre-stained protein ladder (Bio-Rad, 161-0374). Following the transfer, the membrane was blocked using 2.5% bovine serum albumin (catalog no.: MB083; HiMedia) and incubated in primary antibodies for about an hour, followed by subsequent washes with 1X TBST solution (Tris-buffered saline and Tween-20) containing Tris-Cl (pH 7.5), NaCl, and Tween-20. After washing, the membranes were incubated with IR-dye-tagged secondary antibodies, followed by subsequent washes with 1X TBST, and then blots were air dried and scanned by utilizing the LI-COR infrared imaging machine. The following primary antibodies were used: α-GFP (catalog no.: G1544; Sigma), α-TBP polyclonal antiserum raised in rabbit, α-Myc monoclonal primary antibody (catalog no.: MA1-980; Invitrogen), and the following secondary antibodies were used: goat anti-rabbit IgG secondary antibody (catalog no.: A32734; Invitrogen) and goat anti-mouse IgG secondary antibody (catalog no.: 926-32210; Odyssey). Molecular weight protein loading markers (10-250 kDa) were loaded (BioRad catalog no. 1610374) in each western blot. The western blotting with anti-TBP antibody was performed as a protein loading control. The protein levels in the western blots were quantified by using ImageJ software. The signals of the mature and immature form of the model proteins (Gas1 and CPY) were normalized to TBP (loading control), and the ratio of immature to mature forms were calculated. To determine the total protein level of either Gas1 or CPY, the normalized intensities were added. The percentage of immature or mature protein in relation to the total protein level was calculated using the formula (normalized intensity of immature or mature protein/intensity of total protein) ∗ 100.

## Acknowledgements

We are thankful to Prof. C.D. Allis for gifting us histone H2A mutant yeast strains; Prof. Won Ki Huh for gifting us Gas1-GFP plasmid. We acknowledge all the laboratory members for their assistance and suggestions throughout this study.

## Author’s contribution

V.A., S.A., P.T., S.A., R.P., S.S., S.S.A, R.M., D.R. and R.S. conceptualization, investigations, methodology, validation, and formal analysis; R.S.T supervision, conceptualization and writing-original draft; R.S.T., V.A., S.A., P.T., S.A., R.P., S.S. and R.S. writing – review & editing; R.S.T. project administration and funding acquisition.

## Funding and additional information

This work was supported by funds from IISER Bhopal.

