## Supplementary file for "Metabolic regulation of copper homeostasis governs the Sec61-dependent protein translocation process in *Saccharomyces cerevisiae*"

#### Supplementary information

### Figure S1

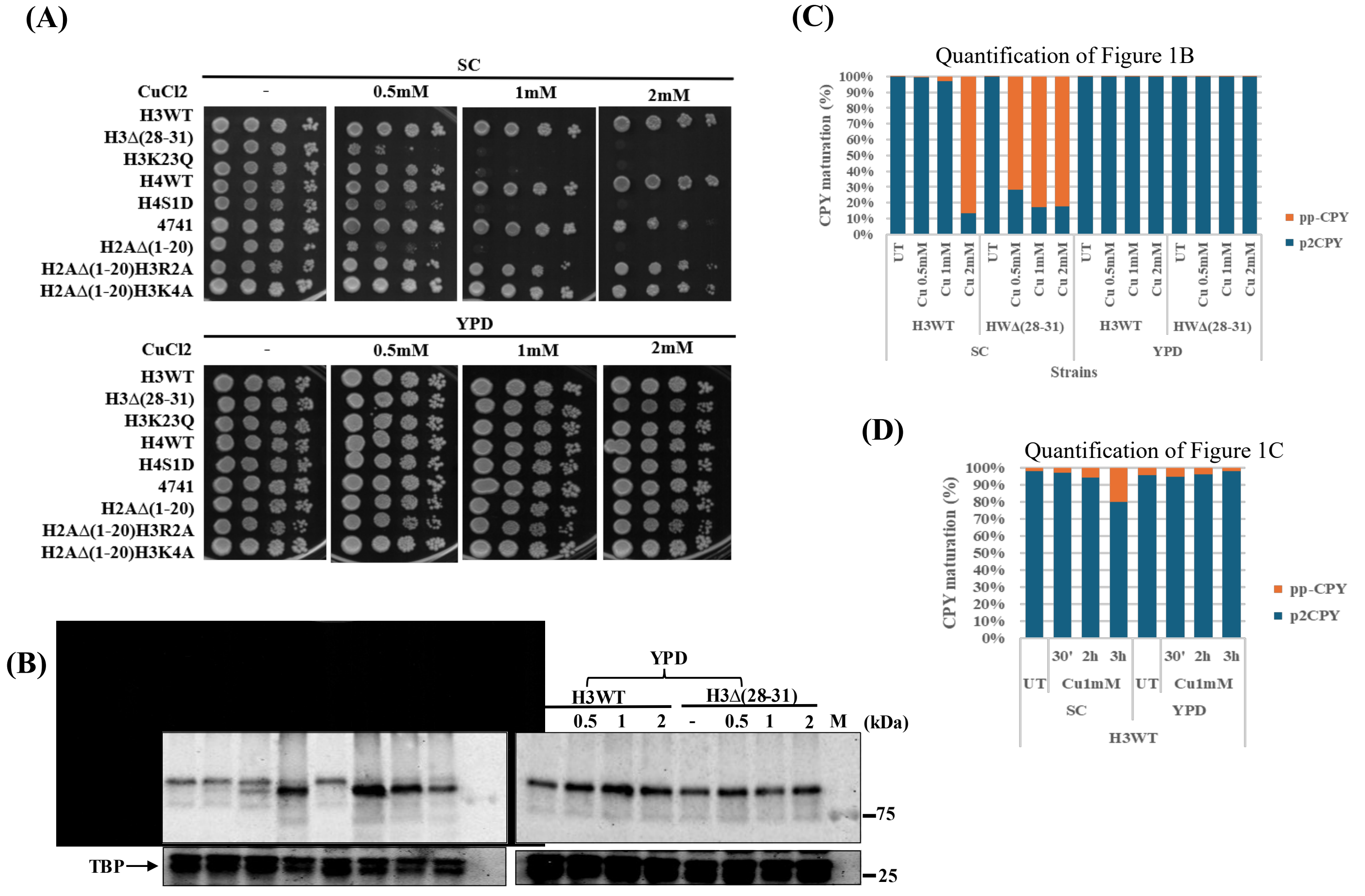


**Figure S1: Growth media regulate copper homeostasis and protein translocation process.** (A) Spot test assay of some of the copper-sensitive histone H3, H4, and H2A mutants along with their respective wild type on SC agar or YPD agar solid growth medium supplemented with and without copper chloride. Overnight grown cultures were diluted to set 1OD600 and 10-fold serially diluted cells, spotted from left to right. Plates were incubated at 30°C and images captured after 72 hours. UT or minus sign means growth of cells on SC medium without copper chloride. (B) Replicate of Western blots presented in Figure 1(B), myc-tagged CPY in a copper-sensitive histone H3 mutant and wild-type cells. Overnight grown cultures were seeded at 0.2OD600 in SC and YPD liquid medium and grown till 1OD600, allowed to grow for 2 hours in untreated and in the presence of copper chloride (0.5, 1, and 2mM). Whole cell protein extracts (WCE) were prepared for western blotting. Western blotting with TBP served as a protein loading control. SC means synthetic complete media, and YPD means yeast extract, peptone, and dextrose. TBP means TATA Binding Protein. (C-D) Quantification of western blots presented in Figure 1(B and C). Data indicates the ratio of p2CPY form and pp-CPY forms in a histone H3 mutant as compared to wild-type cells.

#### Figure S2

**(A)**

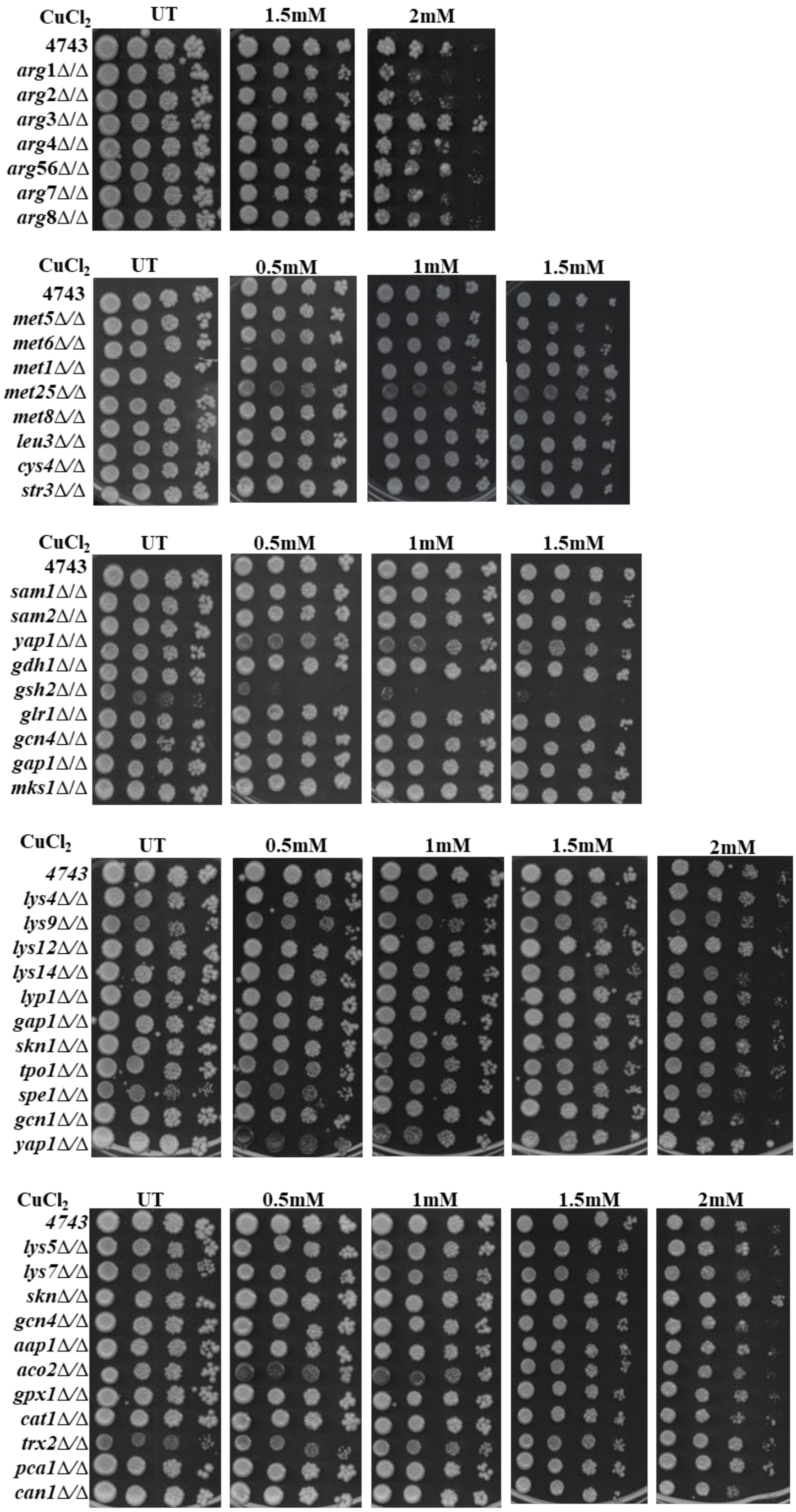
CuCl2
UT
1.5mM
2mM
4743
arg1Δ/Δ
arg2Δ/Δ
arg3Δ/Δ
arg4Δ/Δ
arg56Δ/Δ
arg7Δ/Δ
arg8Δ/Δ
CuCl2
UT
0.5mM
1mM
1.5mM
4743
met5Δ/Δ
met6Δ/Δ
met1Δ/Δ
met25Δ/Δ
met8Δ/Δ
leu3Δ/Δ
cys4Δ/Δ
str3Δ/Δ
CuCl2
UT
0.5mM
1mM
1.5mM
4743
sam1Δ/Δ
sam2Δ/Δ
yap1Δ/Δ
gdh1Δ/Δ
gsh2Δ/Δ
glr1Δ/Δ
gcn4Δ/Δ
gap1Δ/Δ
mks1Δ/Δ
CuCl2
UT
0.5mM
1mM
1.5mM
2mM
4743
lys4Δ/Δ
lys9Δ/Δ
lys12Δ/Δ
lys14Δ/Δ
lyp1Δ/Δ
gap1Δ/Δ
skn1Δ/Δ
tpo1Δ/Δ
spe1Δ/Δ
gcn1Δ/Δ
yap1Δ/Δ
CuCl2
UT
0.5mM
1mM
1.5mM
2mM
4743
lys5Δ/Δ
lys7Δ/Δ
sknΔ/Δ
gcn4Δ/Δ
aap1Δ/Δ
aco2Δ/Δ
gpx1Δ/Δ
cat1Δ/Δ
trx2Δ/Δ
pca1Δ/Δ
can1Δ/Δ

### Figure S2

(B)

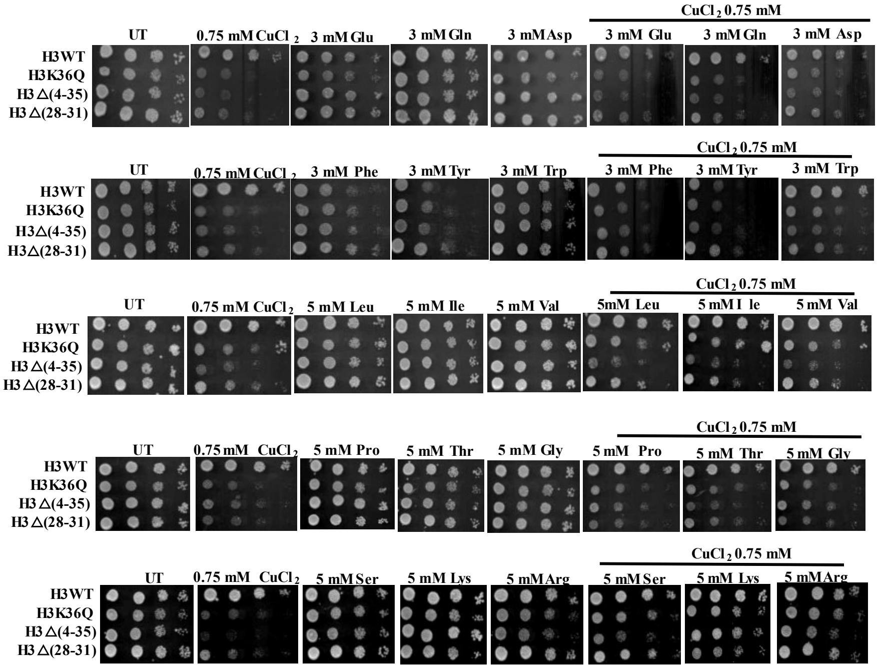
CuCl2 0.75 mM
UT
0.75 mM CuCl2
3 mM Glu
3 mM Gln
3 mM Asp
3 mM Glu
3 mM Gln
3 mM Asp
H3WT
H3K36Q
H3∆(4-35)
H3∆(28-31)
CuCl2 0.75 mM
UT
0.75 mM CuCl2
3 mM Phe
3 mM Tyr
3 mM Trp
3 mM Phe
3 mM Tyr
3 mM Trp
CuCl2 0.75 mM
UT
0.75 mM CuCl2
5 mM Leu
5 mM Ile
5 mM Val
5mM Leu
5 mM Ile
5 mM Val
CuCl2 0.75 mM
UT
0.75 mM CuCl2
5 mM Pro
5 mM Thr
5 mM Gly
5 mM Pro
5 mM Thr
5 mM Gly
CuCl2 0.75 mM
UT
0.75 mM CuCl2
5 mM Ser
5 mM Lys
5 mM Arg
5 mM Ser
5 mM Lys
5 mM Arg

(C)

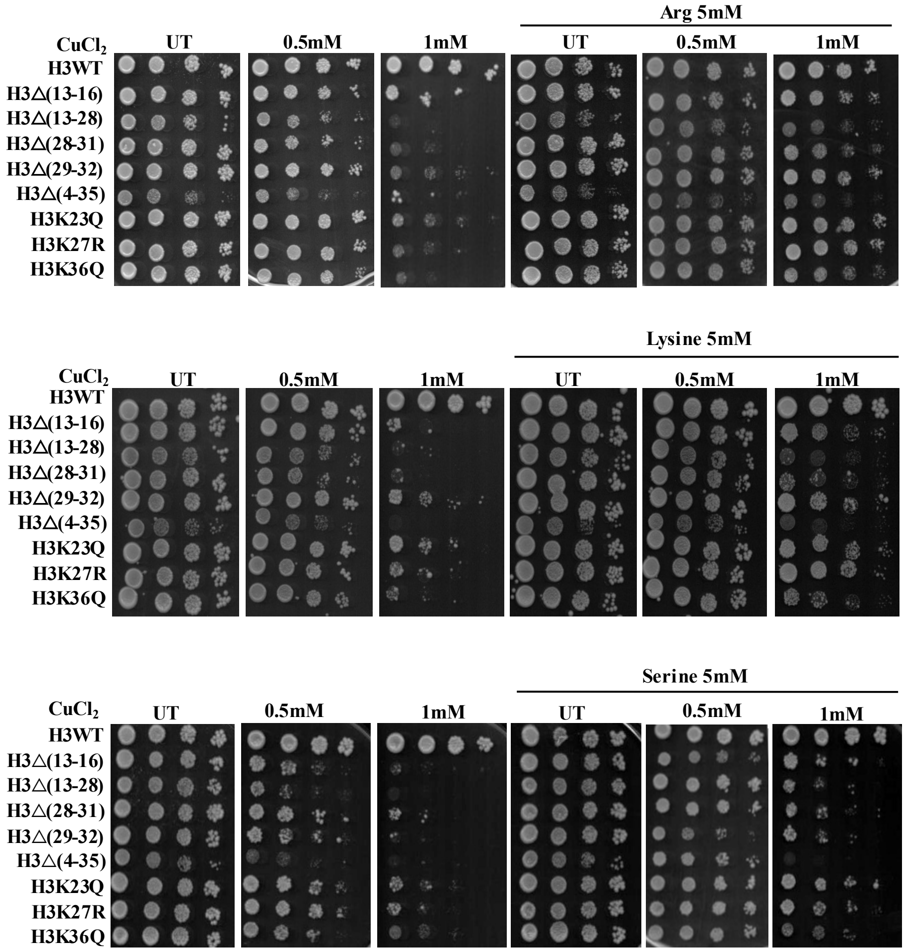
Arg 5mM
CuCl2
UT
0.5mM
1mM
UT
0.5mM
1mM
H3WT
H3∆(13-16)
H3∆(13-28)
H3∆(28-31)
H3∆(29-32)
H3∆(4-35)
H3K23Q
H3K27R
H3K36Q
Lysine 5mM
Serine 5mM

#### Figure S2

(D)

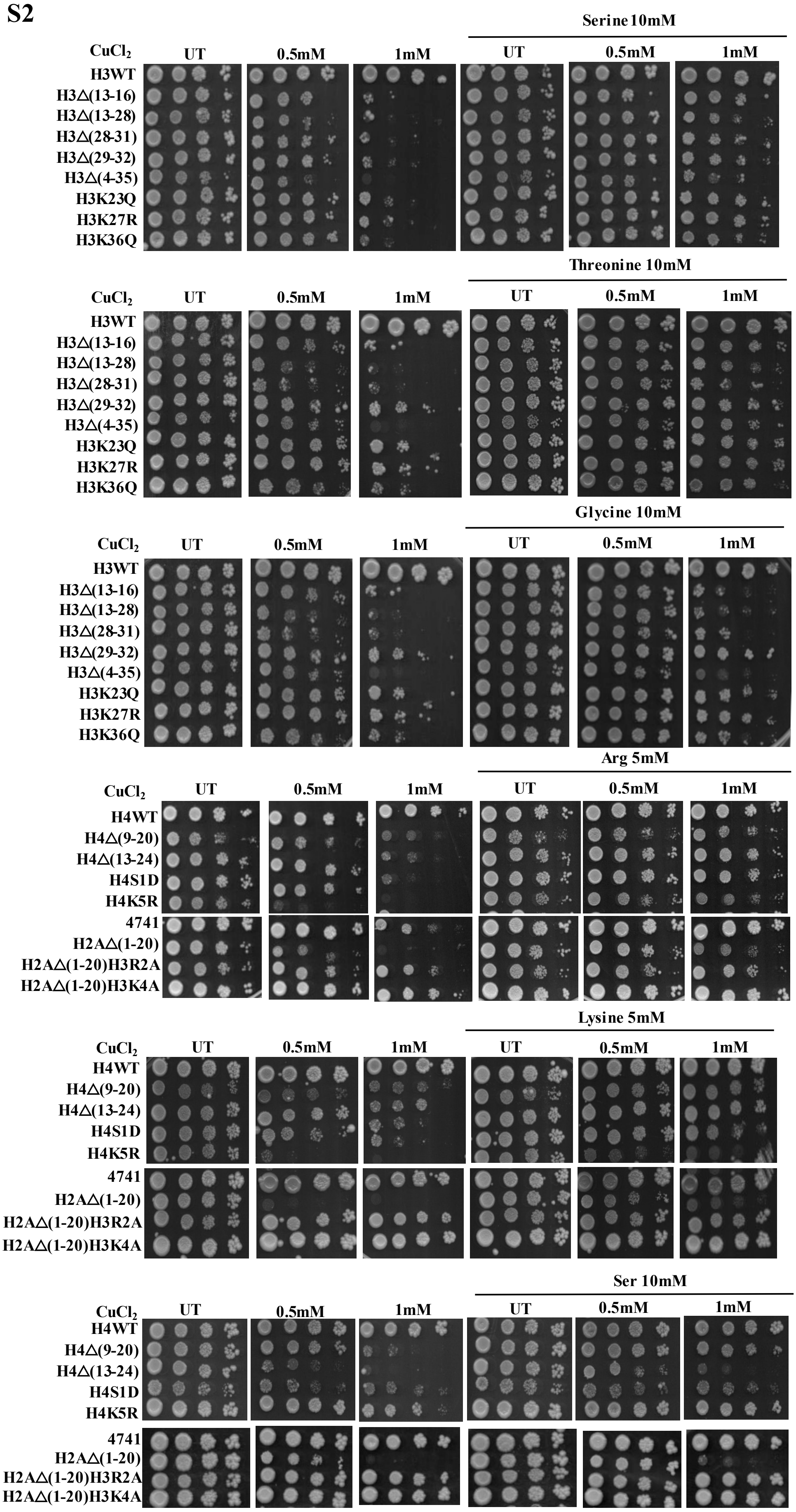
Serine 10mM
CuCl$_2$ UT 0.5mM 1mM UT 0.5mM 1mM
H3WT
H3△(13-16)
H3△(13-28)
H3△(28-31)
H3△(29-32)
H3△(4-35)
H3K23Q
H3K27R
H3K36Q
Threonine 10mM
CuCl$_2$ UT 0.5mM 1mM UT 0.5mM 1mM
H3WT
H3△(13-16)
H3△(13-28)
H3△(28-31)
H3△(29-32)
H3△(4-35)
H3K23Q
H3K27R
H3K36Q
Glycine 10mM
CuCl$_2$ UT 0.5mM 1mM UT 0.5mM 1mM
H3WT
H3△(13-16)
H3△(13-28)
H3△(28-31)
H3△(29-32)
H3△(4-35)
H3K23Q
H3K27R
H3K36Q
Arg 5mM
CuCl$_2$ UT 0.5mM 1mM UT 0.5mM 1mM
H4WT
H4△(9-20)
H4△(13-24)
H4S1D
H4K5R
4741
H2A△(1-20)
H2A△(1-20)H3R2A
H2A△(1-20)H3K4A
Lysine 5mM
CuCl$_2$ UT 0.5mM 1mM UT 0.5mM 1mM
H4WT
H4△(9-20)
H4△(13-24)
H4S1D
H4K5R
4741
H2A△(1-20)
H2A△(1-20)H3R2A
H2A△(1-20)H3K4A
Ser 10mM
CuCl$_2$ UT 0.5mM 1mM UT 0.5mM 1mM
H4WT
H4△(9-20)
H4△(13-24)
H4S1D
H4K5R
4741
H2A△(1-20)
H2A△(1-20)H3R2A
H2A△(1-20)H3K4A

**Figure S2**

**(E)**

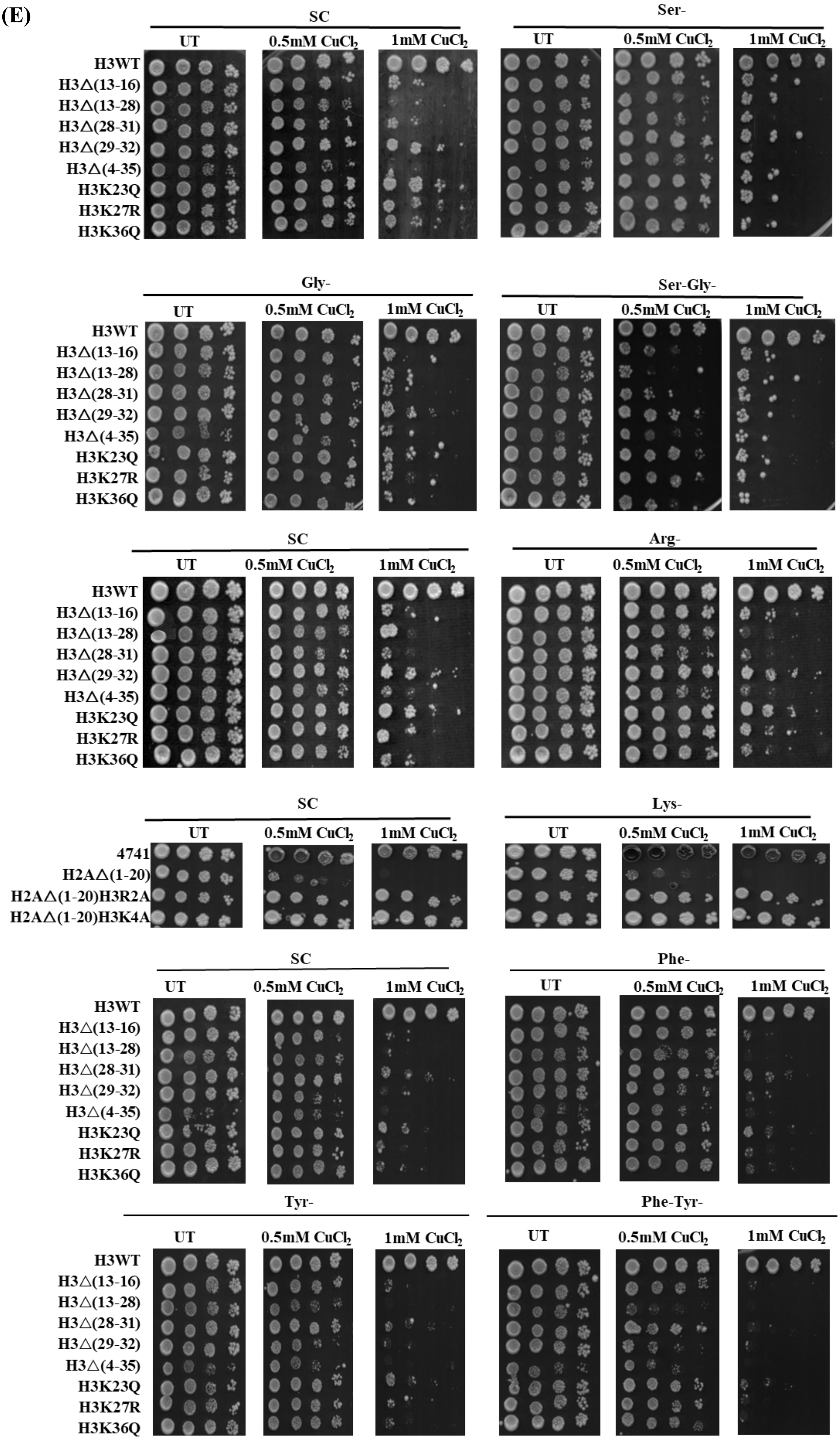
SC
UT
0.5mM $CuCl_2$
1mM $CuCl_2$
Ser-
Gly-
Ser-Gly-
Arg-
Lys-
Phe-
Tyr-
Phe-Tyr-
H3WT
H3Δ(13-16)
H3Δ(13-28)
H3Δ(28-31)
H3Δ(29-32)
H3Δ(4-35)
H3K23Q
H3K27R
H3K36Q
4741
H2AΔ(1-20)
H2AΔ(1-20)H3R2A
H2AΔ(1-20)H3K4A

### Figure S2

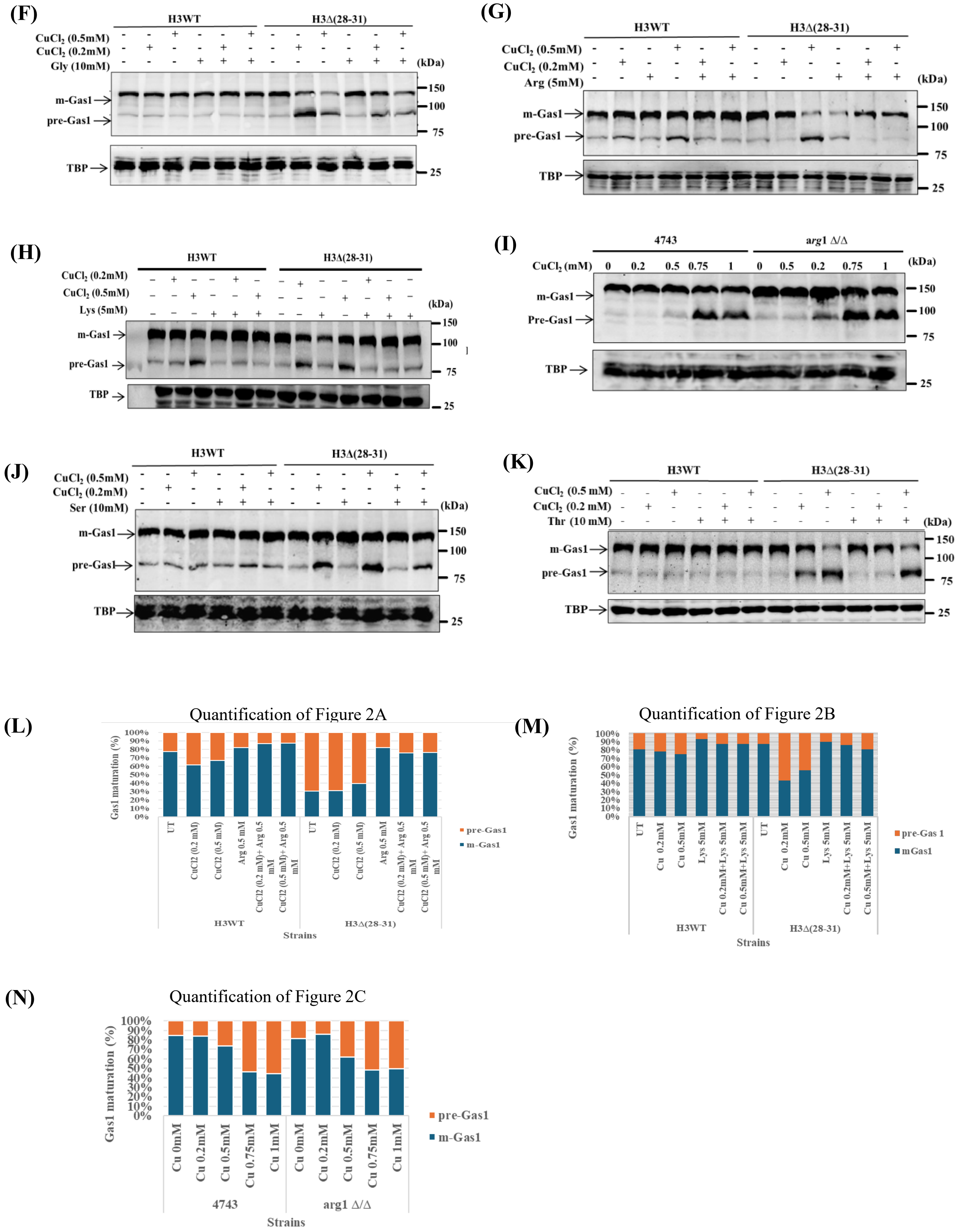
(F)
H3WT
H3Δ(28-31)
CuCl2 (0.5mM)
CuCl2 (0.2mM)
Gly (10mM)
(kDa)
m-Gas1
pre-Gas1
TBP
150
100
75
25
(G)
H3WT
H3Δ(28-31)
CuCl2 (0.5mM)
CuCl2 (0.2mM)
Arg (5mM)
(kDa)
m-Gas1
pre-Gas1
TBP
150
100
75
25
(H)
H3WT
H3Δ(28-31)
CuCl2 (0.2mM)
CuCl2 (0.5mM)
Lys (5mM)
(kDa)
m-Gas1
pre-Gas1
TBP
150
100
75
25
(I)
4743
arg1 Δ/Δ
CuCl2 (mM)
0
0.2
0.5
0.75
1
0
0.5
0.2
0.75
1
(kDa)
m-Gas1
Pre-Gas1
TBP
150
100
75
25
(J)
H3WT
H3Δ(28-31)
CuCl2 (0.5mM)
CuCl2 (0.2mM)
Ser (10mM)
(kDa)
m-Gas1
pre-Gas1
TBP
150
100
75
25
(K)
H3WT
H3Δ(28-31)
CuCl2 (0.5 mM)
CuCl2 (0.2 mM)
Thr (10 mM)
(kDa)
m-Gas1
pre-Gas1
TBP
150
100
75
25
(L)
Quantification of Figure 2A
Gas1 maturation (%)
100%
90%
80%
70%
60%
50%
40%
30%
20%
10%
0%
UT
CuCl2 (0.2 mM)
CuCl2 (0.5 mM)
Arg 0.5 mM
CuCl2 (0.2 mM)+ Arg 0.5 mM
CuCl2 (0.5 mM)+ Arg 0.5 mM
H3WT
H3Δ(28-31)
Strains
pre-Gas1
m-Gas1
(M)
Quantification of Figure 2B
Gas1 maturation (%)
UT
Cu 0.2mM
Cu 0.5mM
Lys 5mM
Cu 0.2mM+Lys 5mM
Cu 0.5mM+Lys 5mM
H3WT
H3Δ(28-31)
Strains
pre-Gas 1
mGas1
(N)
Quantification of Figure 2C
Gas1 maturation (%)
Cu 0mM
Cu 0.2mM
Cu 0.5mM
Cu 0.75mM
Cu 1mM
4743
arg1 Δ/Δ
Strains
pre-Gas1
m-Gas1

#### Figure S2

(O)

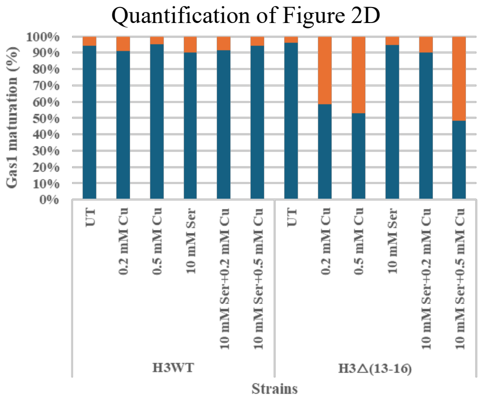


(P)

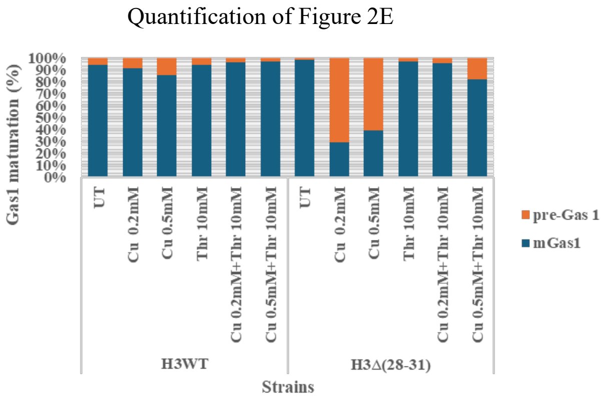


**Figure S2: Screening identifies genes and amino acids required to regulate copper homeostasis and mitigate copper-induced translocation defect.** (A) Spot test assay of double deletion mutants of amino acid metabolism genes and wild type to test their growth on SC agar medium in the absence of copper chloride and in the presence of different concentrations of copper chloride as indicated. (B) Spot test assay of copper sensitive histone H3 mutant and wild type cells on SC agar medium in absence and presence of copper chloride, individual amino acids [Glu (Glutamic acid), Gln (Glutamine), Asp (Aspartic acid), Tyr (Tyrosine), Trp (Tryptophan), Phe (Phenylalanine), Leu (Leucine), Ile (Isoleucine), Val (Valine), Pro (Proline), Thr (Threonine), Gly (Glycine), Ser (Serine), Lys (Lysine), Arg (Arginine)] and in cotreatment with copper chloride and individual amino acids. The concentrations of copper chloride and amino acids were taken as indicated. (C-D) Spot test assay of copper-sensitive histone H3, H4, and H2A histone mutants and wild-type cells on SC agar solid media supplemented with and without copper chloride and individual amino acids (Arg, Lys, Ser) and cotreatment of copper chloride and individual amino acids (concentrations as indicated). Similarly, H3 histone mutants were also spotted on SC agar solid medium in the absence and presence of copper chloride, and individual amino acids Thr (Threonine), Gly (Glycine). (E) Spot assay of copper-sensitive histone H3 mutants on SC agar and SC minus Ser/Gly/Ser and Gly/Arg/Phe/Phe and Tyr agar medium, and spot assay of copper-sensitive histone H2A mutants on SC agar and SC minus Lys agar solid medium with and without copper chloride (0.5 and 1mM). Overnight grown cultures of each strain were 10-fold serially diluted and spotted from left to right, plates were incubated at 30°C, and images captured after 72 hours. UT means untreated, growth of cells on SC media without copper or supplemented with amino acids. (F) Immunoblot of a Gas1-GFP transformed copper sensitive histone H3 mutant and wild type cells in untreated and copper chloride treatment (0.2 and 0.5mM) and Gly (10mM) and cotreatment for 2hrs. (G-K) Replicates of western blots presented in Figure 2(A-E). (G) Immunoblot of a Gas1-GFP transformed copper sensitive mutant and wild type cells treated with copper chloride (0.2 and 0.5mM) and Arg (5mM) and cotreatment for 2hrs. (H) Same as '(G)' but supplemented with 5mM of Lys instead of Arg. (I) Immunoblots of a Gas1-GFP transformed Arg1 deleted and wild type cells, untreated and treated for 2hrs with copper chloride (0.2, 0.5, 0.75 and 1mM). (J) Immunoblots of a Gas1-GFP transformed copper sensitive histone H3 mutant and wild type cells treated with copper chloride (0.2 and 0.5mM) and Ser (10mM) and cotreatment for 2hrs. (K) Immunoblots of a Gas1-GFP transformed copper sensitive histone H3 mutant and wild type cells treated with copper chloride (0.2 and 0.5mM) and Thr (10mM) and cotreatment for 2hrs. Whole cell extracts were used for immunoblotting. Western blot of TBP served as a protein loading control. (L-O) Quantification of western blots presented in Figure 2 (A-E). Data indicates the ratio of pre-Gas1 form and mature Gas1 form in mutants indicated w.r.t. wild type cells.

#### Figure S3

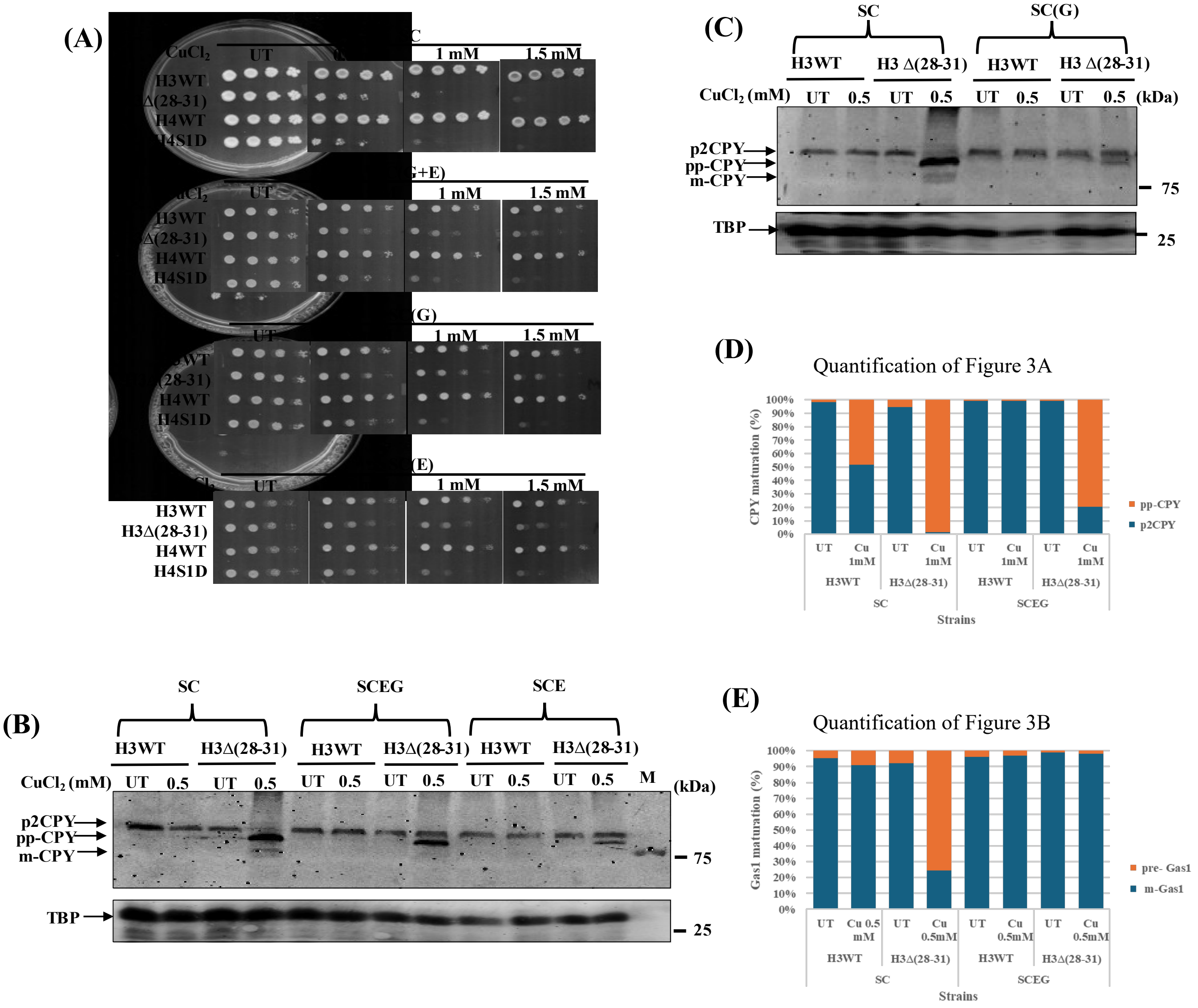


**Figure S3: Effect of different carbon sources on copper homeostasis.** (A) Spot assay of wild type and copper-sensitive histone H3 and H4 mutant, H3Δ(28-31) and H4S1D. Overnight grown cultures of cells in SC media were serially diluted (10-fold), and spotted from left to right on SC and non-fermentable carbon sources (NFC); SCEG, SCG, SCE in solid agar media supplemented with or without copper. UT means untreated. (B-C) Western blot of myc-tagged CPY in a copper-sensitive histone H3 mutant and its respective wild type cells, cells from an overnight grown culture were seeded at 0.2OD600 in SC, SC-EG, SC-E, and SC-G liquid medium, grown till 1OD600, and grown for 2 hours in untreated and in the presence of copper chloride. Whole cell extracts were prepared for the immunoblotting. The western blot analysis of TBP served as a control for protein loading. SC refers to synthetic complete media containing glucose. SC(E) indicates SC media containing ethanol in place of glucose. SC(G) denotes SC media containing glycerol instead of glucose, while SCEG refers to SC media containing glycerol and ethanol. (D) Quantification of western blots illustrated in Figure 3(A). Data indicates the ratio of p2CPY form and pp-CPY forms in a histone H3 mutant as compared to wild-type cells. (E) Quantification of western blots illustrated in Figure 3(B). Data indicates the ratio of pre-Gas1 form and mature Gas1 form in mutants as compared to wild-type cells.

### Figure S3

(F)

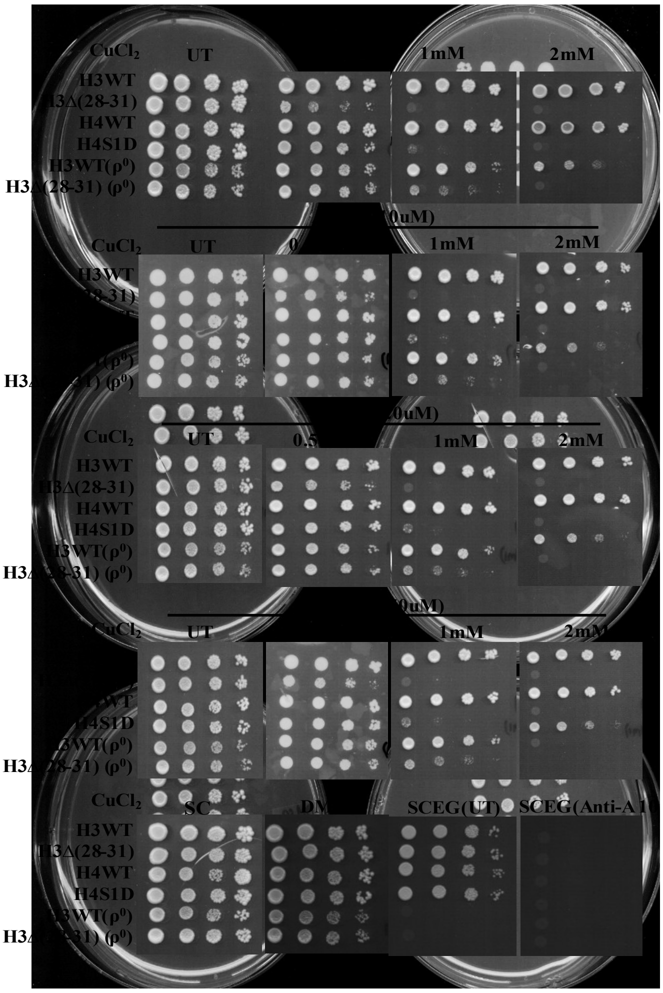


(G)

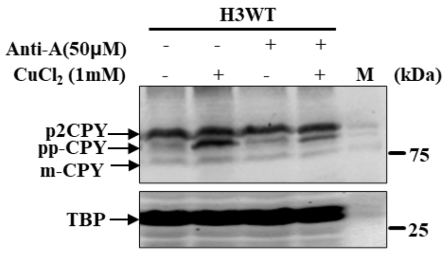


(H)

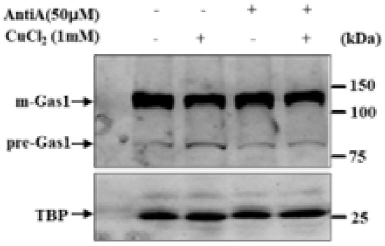


(I)

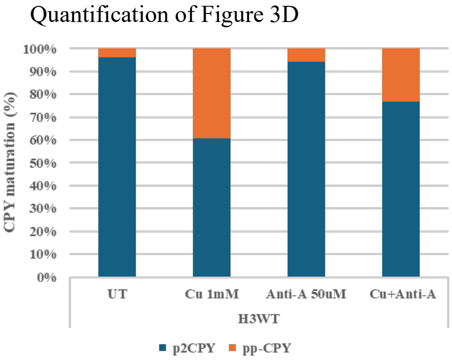


(J)

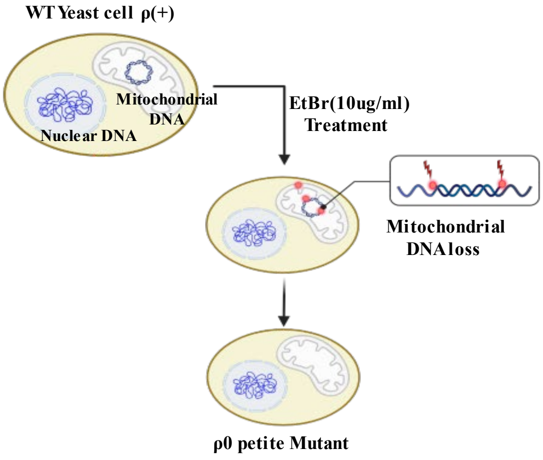

**Figure S3**

(K)

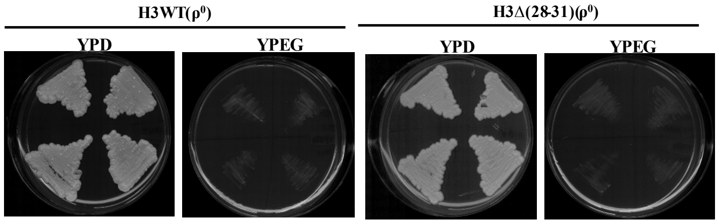


(L)

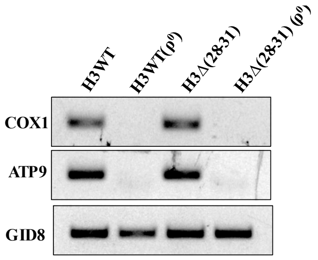


(M)

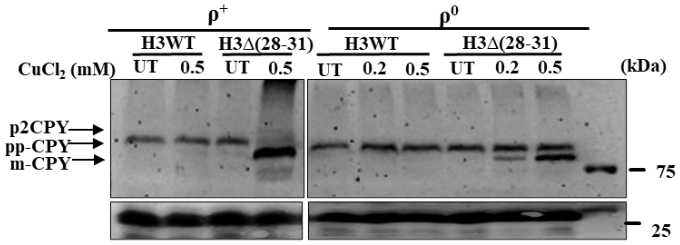


(N)

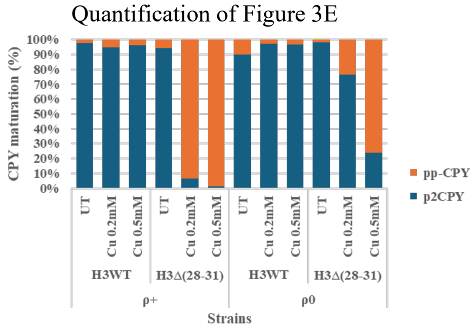


**Figure S3: Mitochondrial genome regulates copper homeostasis.** (F) Spot assay of copper-sensitive histone mutants, H3Δ(28-31) and H4S1D, along with their respective wild type and rho mutant of H3WT and H3Δ(28-31). Cells were grown overnight in SC media, serially diluted (10-fold), and spotted from left to right on SC solid agar supplemented with and without copper chloride, Antimycin-A, and in cotreatment with copper and Antimycin-A. UT means untreated. (G) Replicate of western blots presented in Figure 3(D). Western Blot of myc-tagged CPY in wild type cells, cells from an overnight grown culture were seeded at 0.2OD600 in SC liquid medium, grown till 1OD600, supplemented with and without copper chloride, Antimycin-A, and cotreatment of both copper and Antimycin-A for 2hrs, WCE used for immunoblotting. (H) Same as '(G)', but the Gas1-GFP plasmid transformed H3WT strain is used. (I) Quantification of western blots presented in Figure 3(D). Data indicates the ratio of p2CPY and pp-CPY forms in a histone H3 wild type cells. (J) Schematic representation of $\rho^0$ (Rho Zero) cells generation by EtBr (Ethidium Bromide) treatment. (K) Confirmation of $\rho^0$ (Rho Zero) cells by growing them on YPEG medium. (L) Confirmation of $\rho^0$ (Rho Zero) cells by semi-quantitative PCR using primers of mitochondrial genes. (M) Biological replicate of western blots presented in Figure 3(E). Western Blot of myc-tagged CPY in ρ+ (Rho plus) and $\rho^0$ (Rho Zero) cells of wild type and histone H3 mutant cells, from an overnight grown culture, were seeded at 0.2OD600 in SC liquid medium, grown till 1OD600, treated and untreated with copper chloride for 2hrs, WCE were prepared for immunoblotting. The western blot analysis of TBP served as a control for protein loading. SC means synthetic complete media. WCE means whole cell extract. (N) Quantification of western blots illustrated in Figure 3(E). Data indicates the ratio of p2CPY form and pp-CPY forms in a histone H3 mutant compared to wild-type cells.

#### Figure S3

(O)

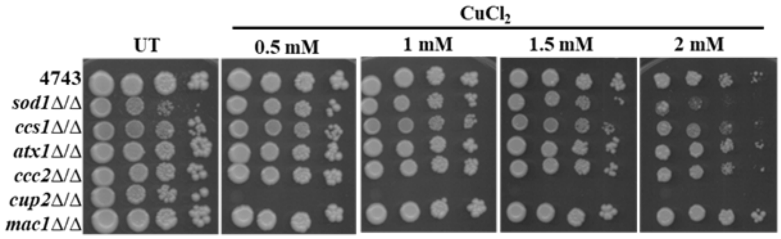


(P)

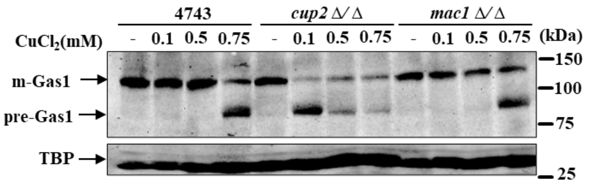


(Q)

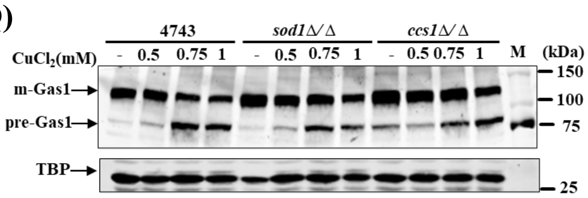


(R)

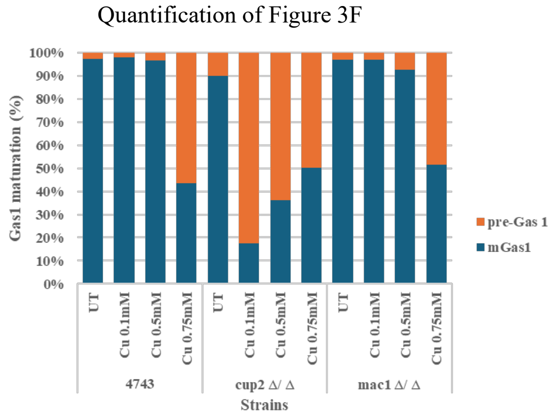


(S)

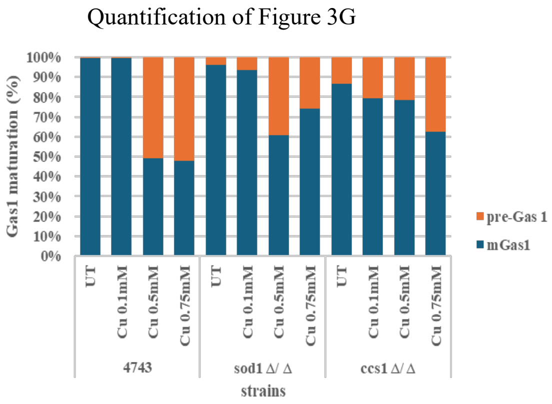


**Figure S3: Copper response genes maintain a labile pool of copper and the Sec61-mediated protein translocation process.** (O) Spot test assay of double deletion mutants of copper response genes; copper chaperones, copper regulon, and wild type to test their growth on SC agar medium in the absence and presence of copper chloride (0.5, 1, 1.5, 2mM). 10-fold serially diluted cells were spotted from left to right, UT means untreated. (P-Q) Replicate of western blots presented in Figure 3(F-G). (P) Immunoblots of Gas1-GFP transformed double deletion mutants of *cup2*Δ/Δ, *mac1*Δ/Δ and wild type, exponentially growing cells treated with and without copper chloride (0.5, 0.75 and 1mM) for 2hrs. (Q) Immunoblots of Gas1-GFP transformed double deletion mutants of *sod1*Δ/Δ, *ccs1*Δ/Δ and wild type, exponentially growing cells treated with and without copper chloride (0.5, 0.75 and 1mM) for 2hrs. The Western blot analysis of TBP served as a control for protein loading. (R-S) Quantification of western blots illustrated in Figure 3(F-G). Data indicates the ratio of pre-Gas1 form and mature Gas1 form in mutants compared to wild-type cells.

**Figure S4**

**(A)**

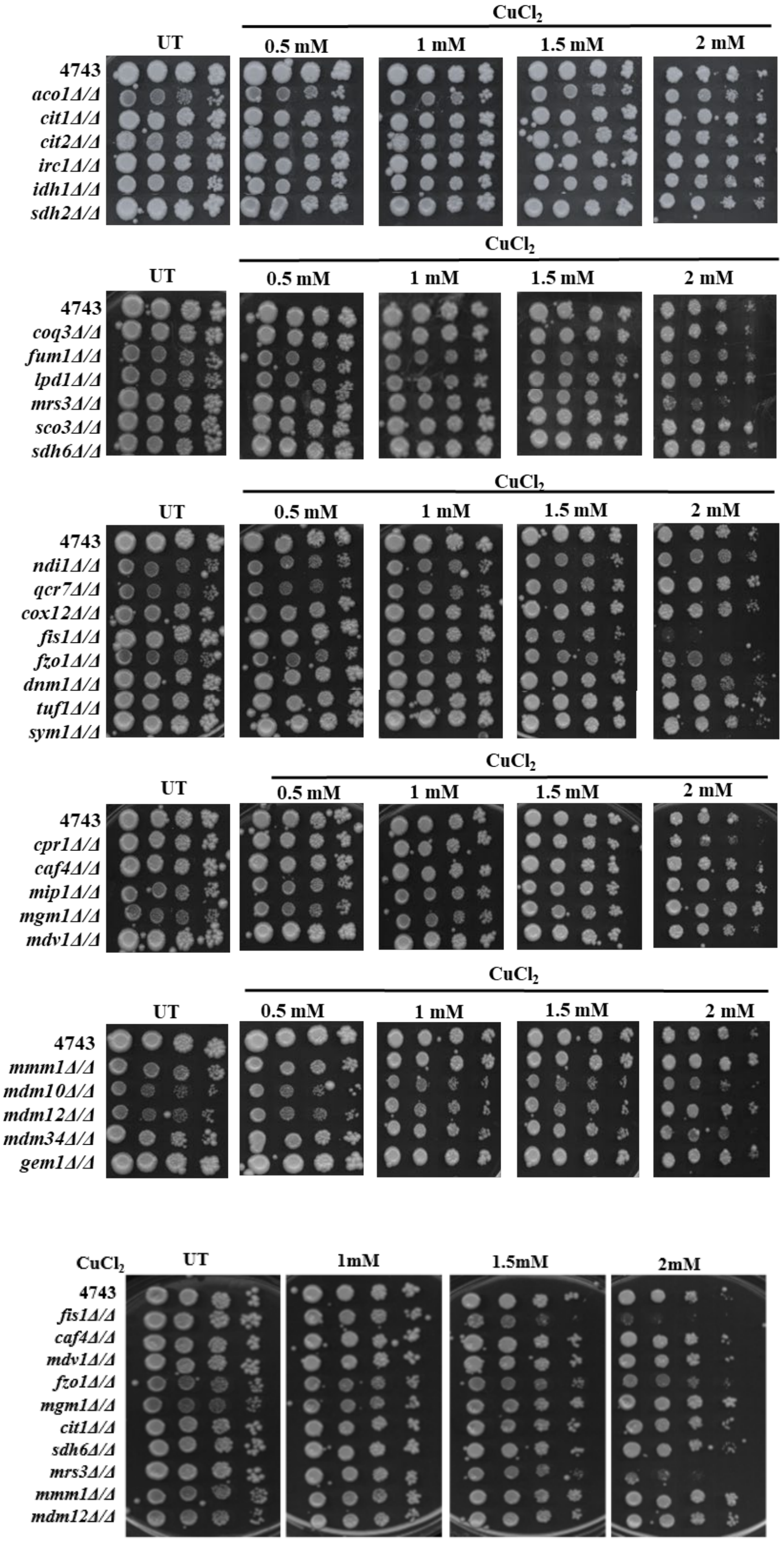

CuCl2
UT
0.5 mM
1 mM
1.5 mM
2 mM
4743
aco1Δ/Δ
cit1Δ/Δ
cit2Δ/Δ
irc1Δ/Δ
idh1Δ/Δ
sdh2Δ/Δ
CuCl2
UT
0.5 mM
1 mM
1.5 mM
2 mM
4743
coq3Δ/Δ
fum1Δ/Δ
lpd1Δ/Δ
mrs3Δ/Δ
sco3Δ/Δ
sdh6Δ/Δ
CuCl2
UT
0.5 mM
1 mM
1.5 mM
2 mM
4743
ndi1Δ/Δ
qcr7Δ/Δ
cox12Δ/Δ
fis1Δ/Δ
fzo1Δ/Δ
dnm1Δ/Δ
tuf1Δ/Δ
sym1Δ/Δ
CuCl2
UT
0.5 mM
1 mM
1.5 mM
2 mM
4743
cpr1Δ/Δ
caf4Δ/Δ
mip1Δ/Δ
mgm1Δ/Δ
mdv1Δ/Δ
CuCl2
UT
0.5 mM
1 mM
1.5 mM
2 mM
4743
mmm1Δ/Δ
mdm10Δ/Δ
mdm12Δ/Δ
mdm34Δ/Δ
gem1Δ/Δ
CuCl2
UT
1mM
1.5mM
2mM
4743
fis1Δ/Δ
caf4Δ/Δ
mdv1Δ/Δ
fzo1Δ/Δ
mgm1Δ/Δ
cit1Δ/Δ
sdh6Δ/Δ
mrs3Δ/Δ
mmm1Δ/Δ
mdm12Δ/Δ

**Figure S4**

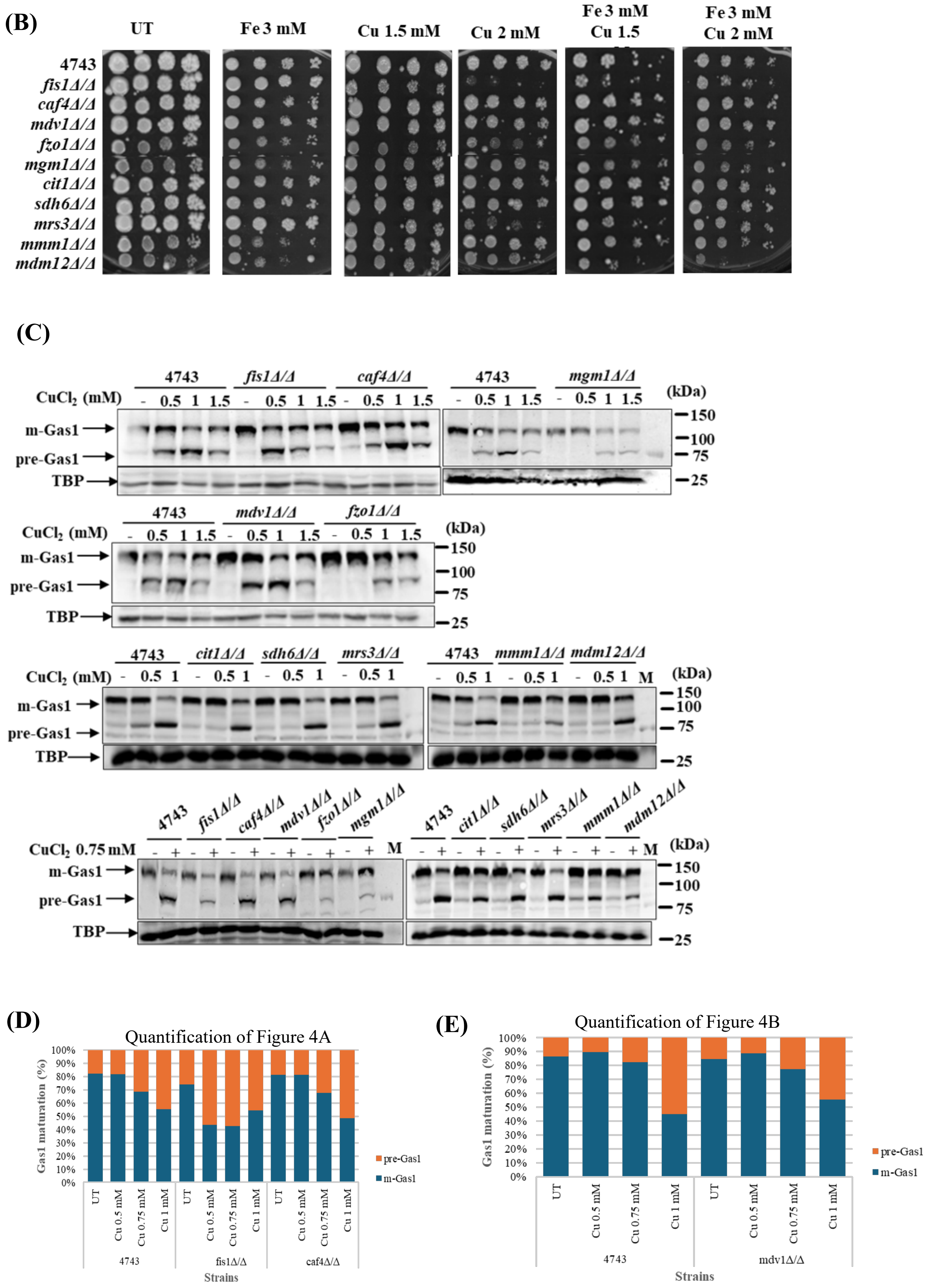

(B)
UT
Fe 3 mM
Cu 1.5 mM
Cu 2 mM
Fe 3 mM Cu 1.5
Fe 3 mM Cu 2 mM
4743
fis1Δ/Δ
caf4Δ/Δ
mdv1Δ/Δ
fzo1Δ/Δ
mgm1Δ/Δ
cit1Δ/Δ
sdh6Δ/Δ
mrs3Δ/Δ
mmm1Δ/Δ
mdm12Δ/Δ
(C)
4743
fis1Δ/Δ
caf4Δ/Δ
mgm1Δ/Δ
mdv1Δ/Δ
fzo1Δ/Δ
cit1Δ/Δ
sdh6Δ/Δ
mrs3Δ/Δ
mmm1Δ/Δ
mdm12Δ/Δ
CuCl2 (mM)
CuCl2 0.75 mM
m-Gas1
pre-Gas1
TBP
M
(kDa)
150
100
75
25
(D)
Quantification of Figure 4A
Gas1 maturation (%)
UT
Cu 0.5 mM
Cu 0.75 mM
Cu 1 mM
4743
fis1Δ/Δ
caf4Δ/Δ
Strains
pre-Gas1
m-Gas1
(E)
Quantification of Figure 4B
mdv1Δ/Δ

#### Figure S4

(F)

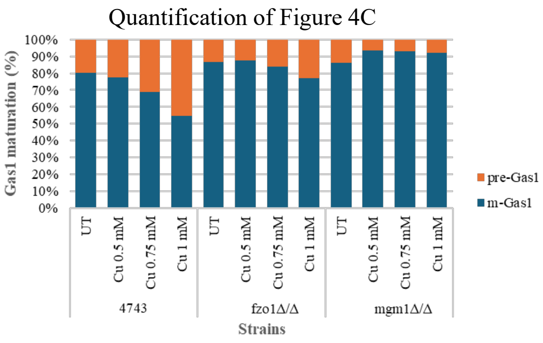


(G)

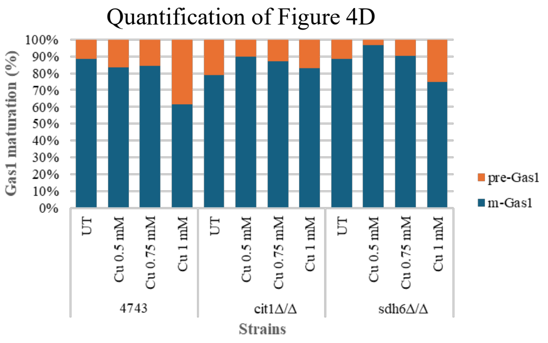


(H)

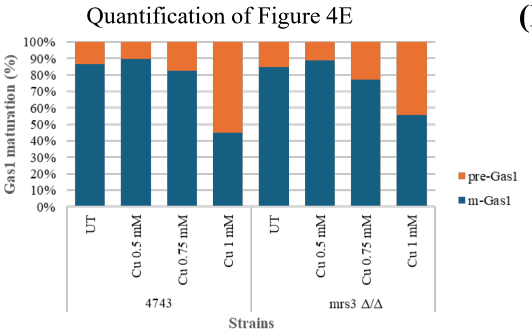


(I)

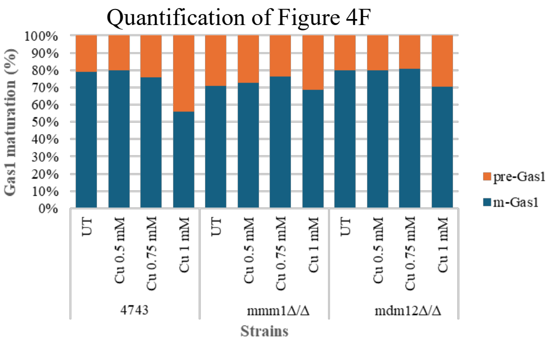


**Figure S4: Mitochondrial fission and fusion regulate copper homeostasis.** (A) Spot test assay of double deletion mutants of mitochondrial respiration, fission, fusion, and ERMES genes and wild type to test their growth on SC agar medium supplemented with and without copper chloride (0.5, 1, 1.5, 2mM). Overnight grown cultures were serially diluted (10-fold), spotted from left to right. (B) Spot test assay of a few double deletion mutants from 'A' of mitochondrial respiration, fission, fusion, and ERMES genes and wild type to test their growth on SC agar medium supplemented with and without copper chloride (1.5 and 2mM), Ferrous chloride 3mM, and cotreatment. Plates were incubated at 30°C and images captured after 72 hours. (C) Replicates of western blots presented in Figure 4(A-F). Western blots of some of the Gas1-GFP transformed double deletion mutants of mitochondrial respiration, fission, fusion, and ERMES genes and wild type cells untreated and treated with copper chloride (0.5, 0.75, and 1mM) for 2hrs. Western blot of TBP served as a protein loading control. (D-I) Quantification of western blots presented in Figure 4(A-F). Data indicates the ratio of pre-Gas1 form and mature Gas1 form in mutants indicated w.r.t. wild type cells.

### Figure S5

(A)

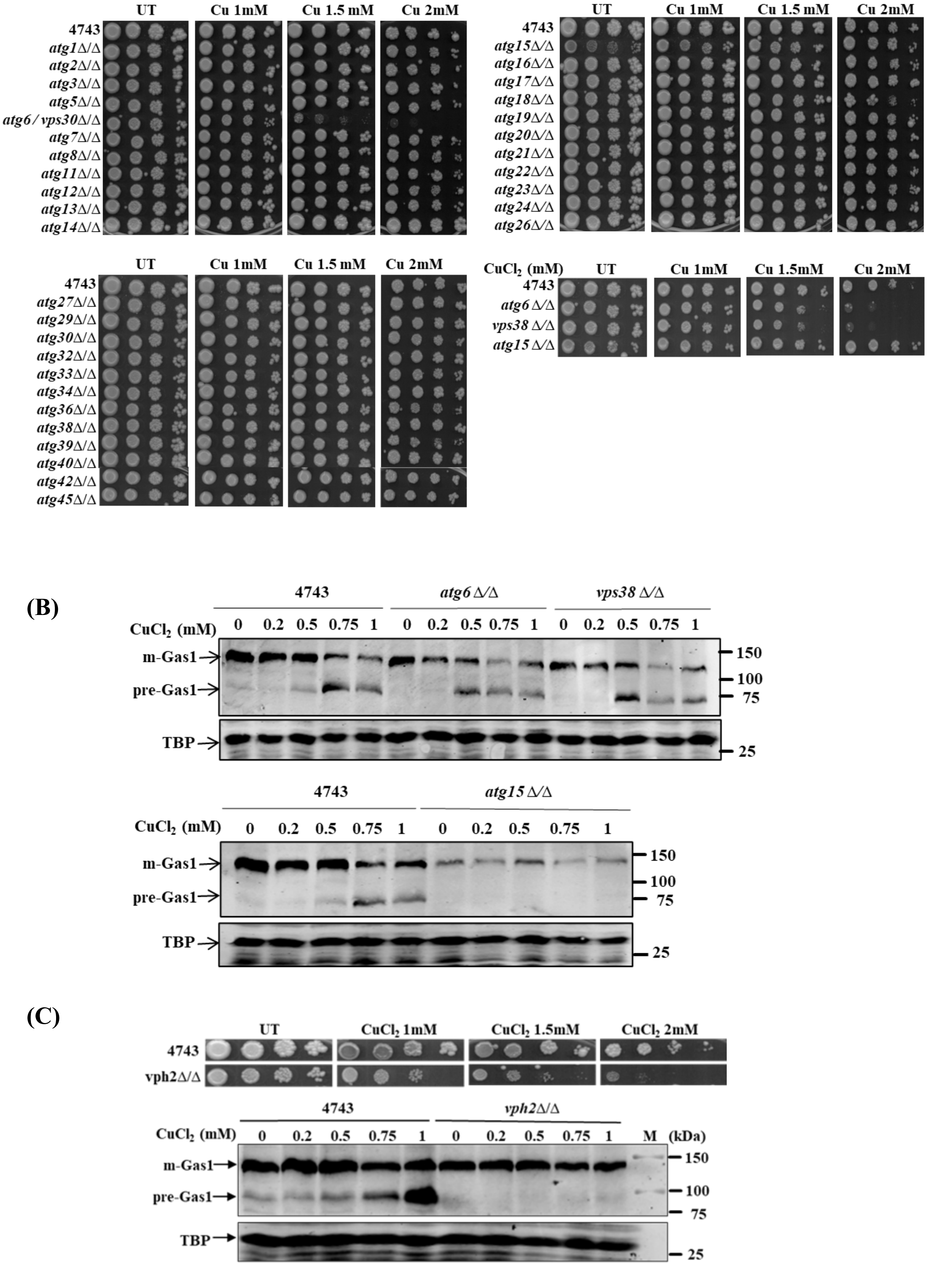

UT
Cu 1mM
Cu 1.5 mM
Cu 2mM
4743
atg1Δ/Δ
atg2Δ/Δ
atg3Δ/Δ
atg5Δ/Δ
atg6 / vps30Δ/Δ
atg7Δ/Δ
atg8Δ/Δ
atg11Δ/Δ
atg12Δ/Δ
atg13Δ/Δ
atg14Δ/Δ
atg15Δ/Δ
atg16Δ/Δ
atg17Δ/Δ
atg18Δ/Δ
atg19Δ/Δ
atg20Δ/Δ
atg21Δ/Δ
atg22Δ/Δ
atg23Δ/Δ
atg24Δ/Δ
atg26Δ/Δ
atg27Δ/Δ
atg29Δ/Δ
atg30Δ/Δ
atg32Δ/Δ
atg33Δ/Δ
atg34Δ/Δ
atg36Δ/Δ
atg38Δ/Δ
atg39Δ/Δ
atg40Δ/Δ
atg42Δ/Δ
atg45Δ/Δ
CuCl2 (mM)
Cu 1.5mM
atg6 Δ/Δ
vps38 Δ/Δ
atg15 Δ/Δ
(B)
4743
atg6 Δ/Δ
vps38 Δ/Δ
CuCl2 (mM)
0 0.2 0.5 0.75 1
m-Gas1
pre-Gas1
TBP
150
100
75
25
atg15 Δ/Δ
(C)
CuCl2 1mM
CuCl2 1.5mM
CuCl2 2mM
vph2Δ/Δ
M
(kDa)

#### Figure S5

(D)

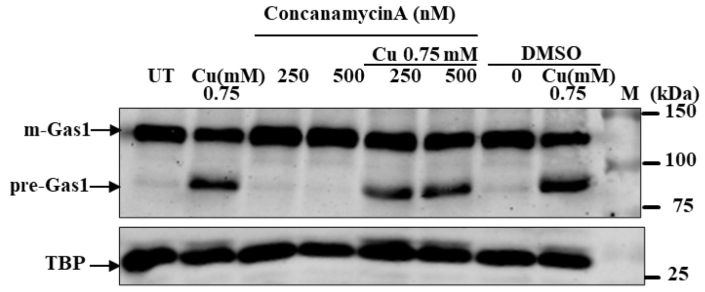


(E)

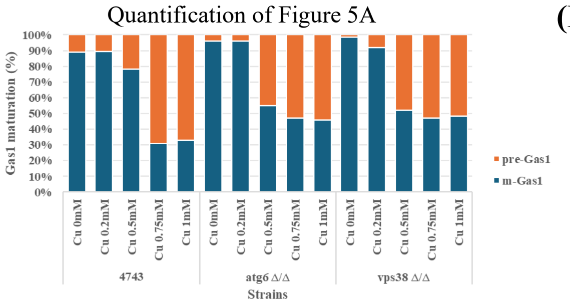


(F)

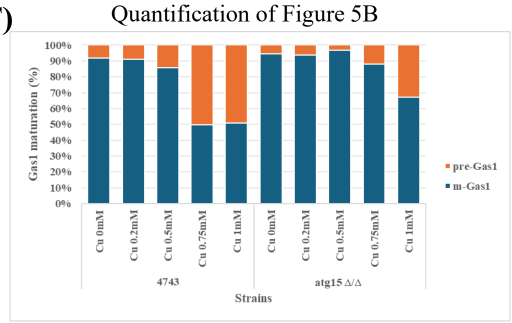


(G)

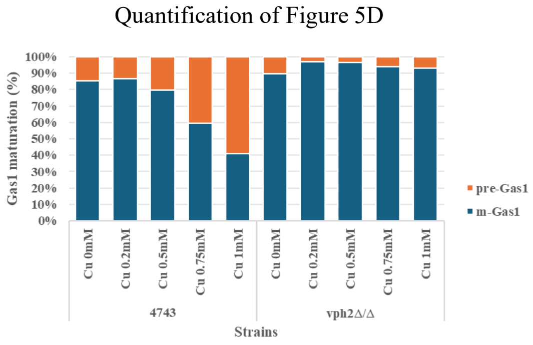


(H)

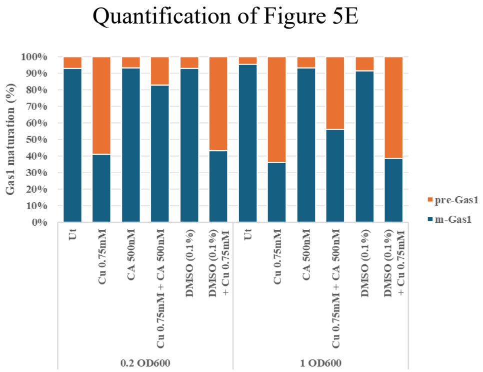


**Figure S5: Autophagy and vacuolar acidification regulate the labile pool of copper.** (A) Spot test assay of double deletion mutants of autophagy pathway genes and wild type to test their growth on SC agar medium supplemented with and without copper chloride (1, 1.5, 2mM), 10-fold serially diluted cells prepared from overnight grown cultures, spotted from left to right. Plates were incubated at 30°C and images captured after 72 hours. (B-D) Replicate of western blot presented in Figure 5(B-D), (B) western blots of Gas1-GFP transformed double deletion mutants of autophagy pathway genes *atg6*Δ/Δ, *vps38*Δ/Δ , and *atg15*Δ/Δ and wild type cells untreated and treated with copper chloride (0.5, 0.75, and 1mM) for 2hrs. (C) Spot assay and western blot of *vph2*Δ/Δ deletion mutant and compared with wild type cells, untreated and treated with copper chloride. (D) Immunoblot of Gas1-GFP transformed wild type cells, untreated and treated with copper chloride (0.75mM), concanamycin-A (250 and 500nM), and cotreated with copper and concanamycin-A for 2hrs. Whole cell extracts were used for immunoblotting. Immunoblot of TBP served as a protein loading control. (E-H) Quantification of western blots presented in Figure 5(A, B, D, E, respectively). Data indicates the ratio of pre-Gas1 form and mature Gas1 form in mutants indicated as compared to wild type cells.

### Figure S6

(A)

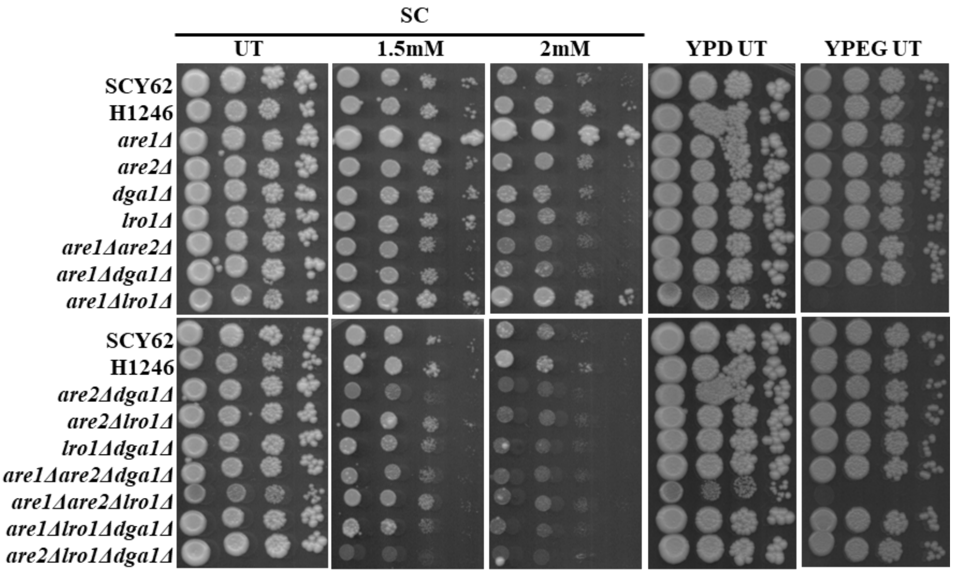


(B)

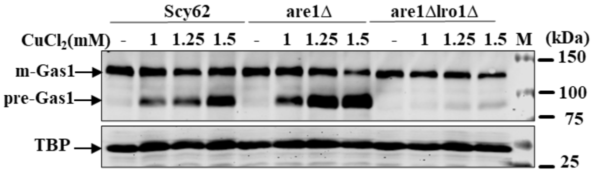


(C)

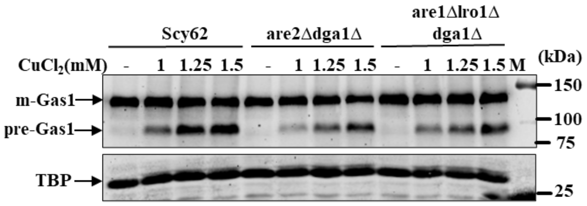


(D)

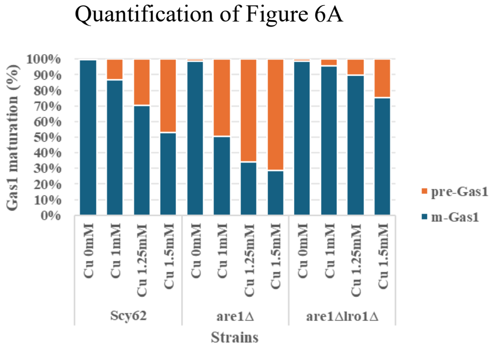


(E)

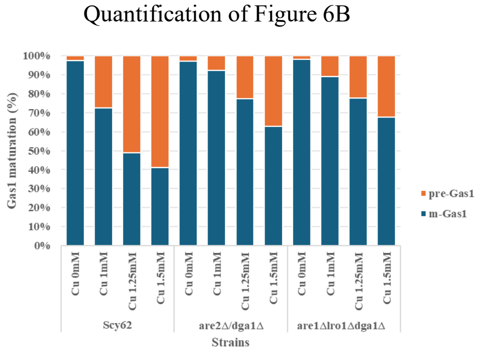

(F)

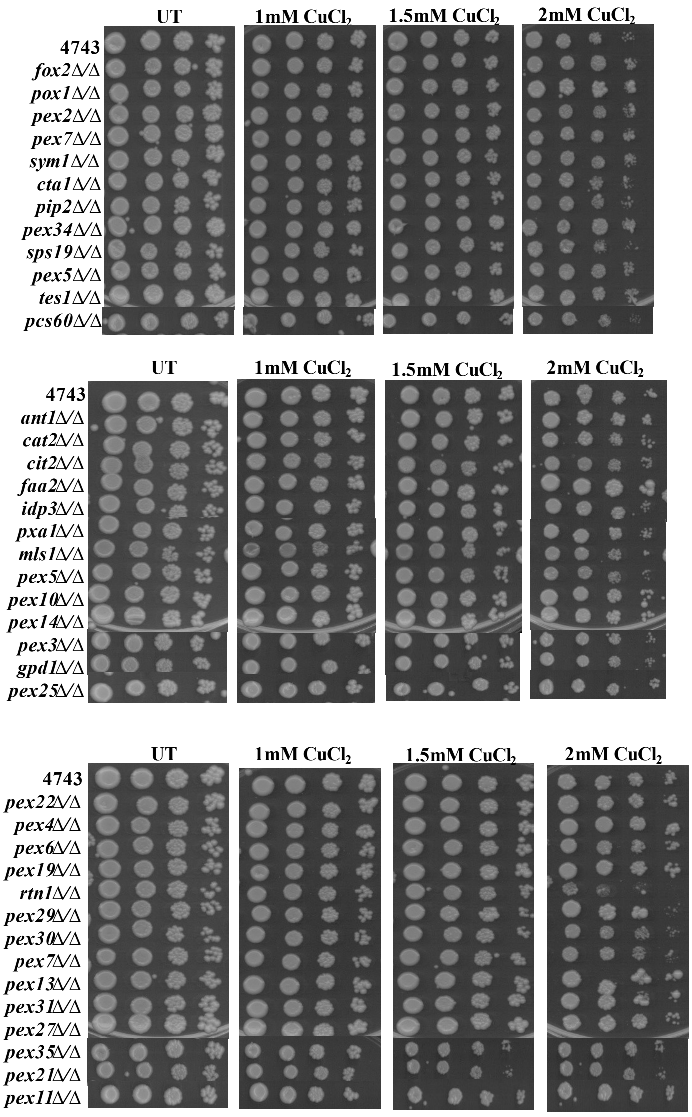
UT
1mM CuCl$_2$
1.5mM CuCl$_2$
2mM CuCl$_2$
4743
fox2Δ/Δ
pox1Δ/Δ
pex2Δ/Δ
pex7Δ/Δ
sym1Δ/Δ
cta1Δ/Δ
pip2Δ/Δ
pex34Δ/Δ
sps19Δ/Δ
pex5Δ/Δ
tes1Δ/Δ
pcs60Δ/Δ
UT
1mM CuCl$_2$
1.5mM CuCl$_2$
2mM CuCl$_2$
4743
ant1Δ/Δ
cat2Δ/Δ
cit2Δ/Δ
faa2Δ/Δ
idp3Δ/Δ
pxa1Δ/Δ
mls1Δ/Δ
pex5Δ/Δ
pex10Δ/Δ
pex14Δ/Δ
pex3Δ/Δ
gpd1Δ/Δ
pex25Δ/Δ
UT
1mM CuCl$_2$
1.5mM CuCl$_2$
2mM CuCl$_2$
4743
pex22Δ/Δ
pex4Δ/Δ
pex6Δ/Δ
pex19Δ/Δ
rtn1Δ/Δ
pex29Δ/Δ
pex30Δ/Δ
pex7Δ/Δ
pex13Δ/Δ
pex31Δ/Δ
pex27Δ/Δ
pex35Δ/Δ
pex21Δ/Δ
pex11Δ/Δ

(G)

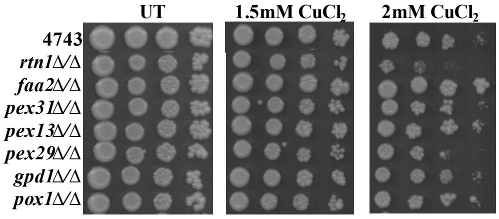
UT
1.5mM CuCl$_2$
2mM CuCl$_2$
4743
rtn1Δ/Δ
faa2Δ/Δ
pex31Δ/Δ
pex13Δ/Δ
pex29Δ/Δ
gpd1Δ/Δ
pox1Δ/Δ

(H)

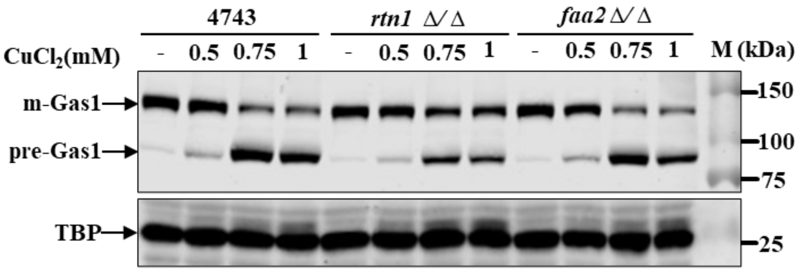


(I)

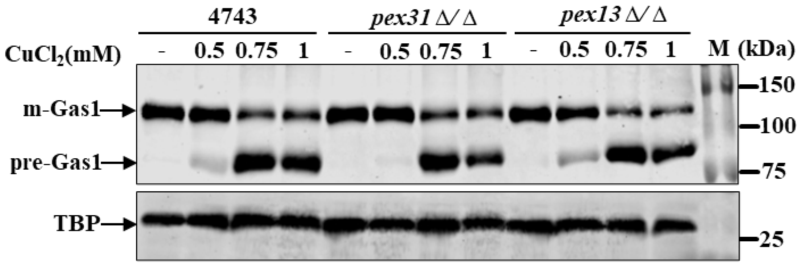


(J)

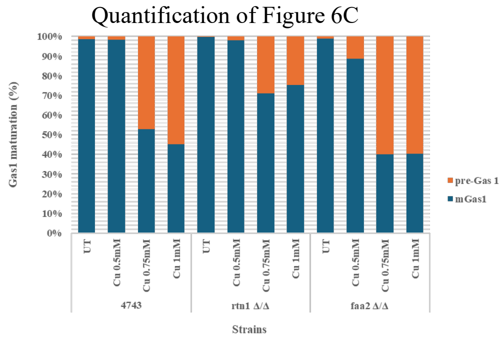


(K)

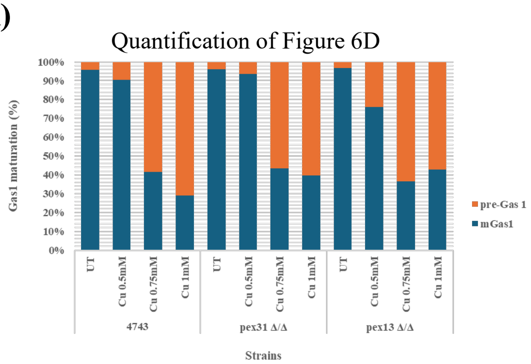


**Figure S6: Lipid droplets and peroxisomal homeostasis regulate copper homeostasis.** (A) Spot test assay of deletion mutants of lipid droplet biogenesis genes and wild type to test their growth on SC agar solid medium supplemented with and without copper chloride (1.5, 2mM) and YPD and YPEG agar solid media without copper. Overnight grown cultures of cells were 10-fold serially diluted, spotted from left to right. (B-C) Replicates of western blots presented in Figure 6(A and B). The western blotting of Gas1-GFP transformed deletion mutants of lipid droplet biogenesis genes and wild type cells grown for 2 hours in the absence and presence of copper chloride (1.0, 1.25, and 1.5mM). Western blot of TBP served as a protein loading control. (D-E) Quantification of western blots presented in Figure 6(A-B). Data indicates the ratio of pre-Gas1 and mature Gas1 forms in mutants indicated w.r.t. wild type cells. (F-G) Spot test assay of double deletion mutants of peroxisomal biogenesis genes and wild type to test their growth on SC agar medium supplemented with and without copper chloride (1.0, 1.5, 2mM), 10-fold serially diluted cells, spotted from left to right. Plates were incubated at 30°C and images were captured after 72 hours. (H-I) Biological replicate of western blot presented in Figure 6(C-D), western blotting of Gas1-GFP transformed deletion mutants of peroxisomal homeostasis gene (*rtn1*Δ/Δ, *faa2*Δ/Δ, *pex31*Δ/Δ *and pex13*Δ/Δ) and respective wild type cells grown in the absence and presence of chloride (0.5, 0.75 and 1mM) for 2hrs. (J-K) Quantification of western blots presented in Figure 6(C-D). Data indicates the ratio of pre-Gas1 and mature Gas1 forms in mutants indicated w.r.t. wild type cells.

#### Figure S7

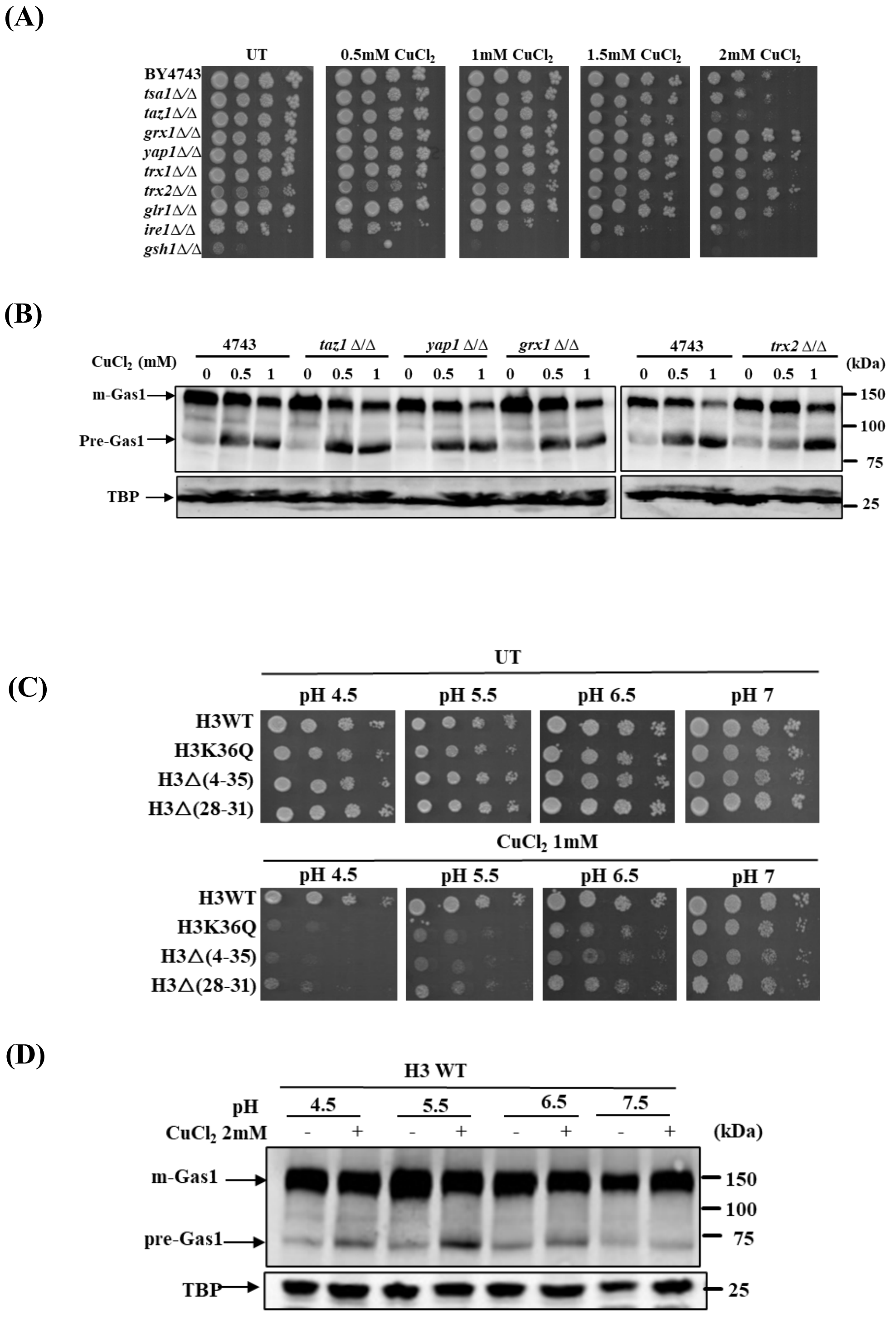

(A)
UT
0.5mM CuCl2
1mM CuCl2
1.5mM CuCl2
2mM CuCl2
BY4743
tsa1Δ/Δ
taz1Δ/Δ
grx1Δ/Δ
yap1Δ/Δ
trx1Δ/Δ
trx2Δ/Δ
glr1Δ/Δ
ire1Δ/Δ
gsh1Δ/Δ
(B)
CuCl2 (mM)
4743
taz1 Δ/Δ
yap1 Δ/Δ
grx1 Δ/Δ
4743
trx2 Δ/Δ
(kDa)
0 0.5 1 0 0.5 1 0 0.5 1 0 0.5 1 0 0.5 1 0 0.5 1
m-Gas1
Pre-Gas1
TBP
150
100
75
25
(C)
UT
pH 4.5
pH 5.5
pH 6.5
pH 7
H3WT
H3K36Q
H3Δ(4-35)
H3Δ(28-31)
CuCl2 1mM
(D)
H3 WT
pH
4.5
5.5
6.5
7.5
CuCl2 2mM
- + - + - + - +
(kDa)
m-Gas1
pre-Gas1
TBP
150
100
75
25

### Figure S7

## (E)

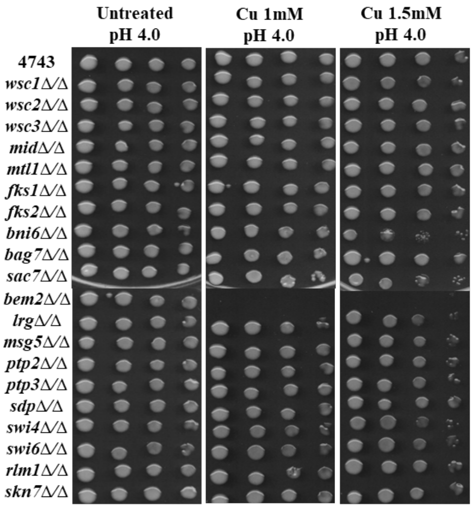


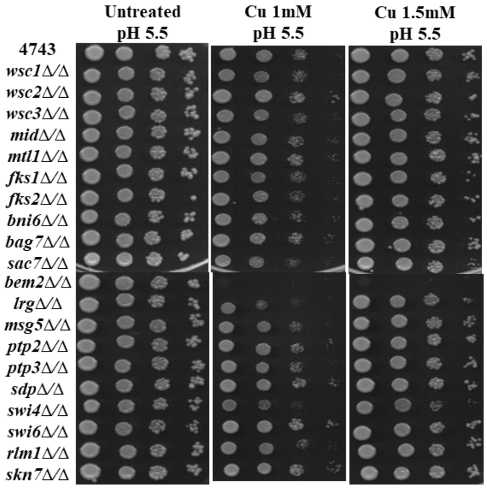


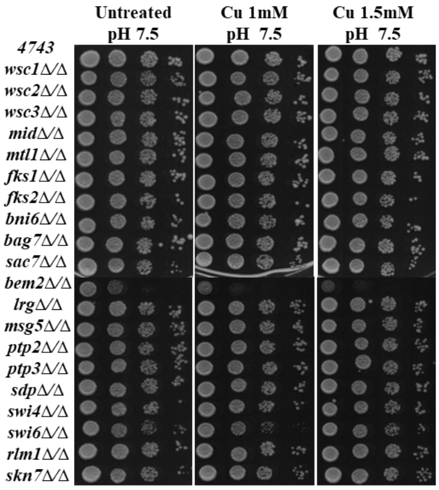


## (F)

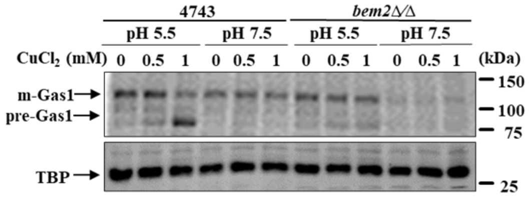


## (G)

Quantification of Figure 7A

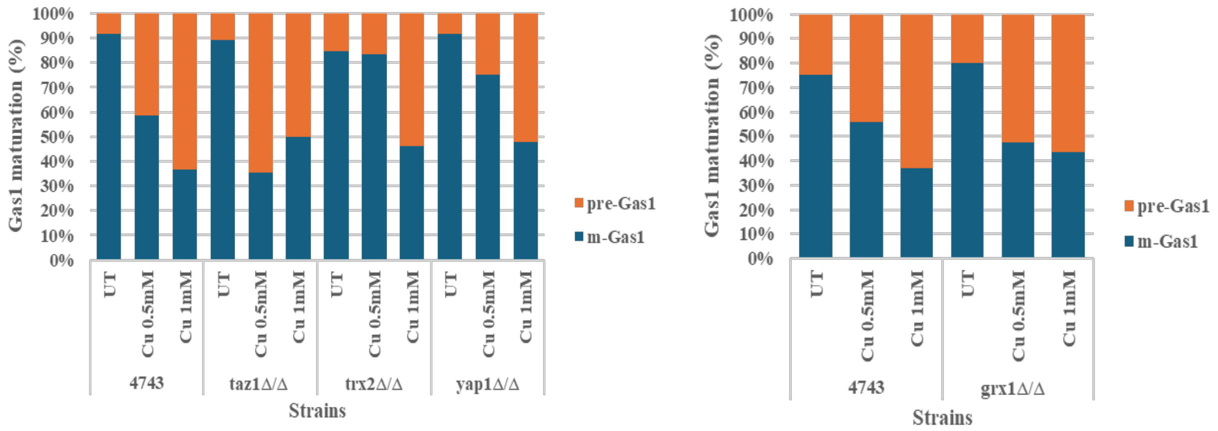

#### Figure S7

(H)

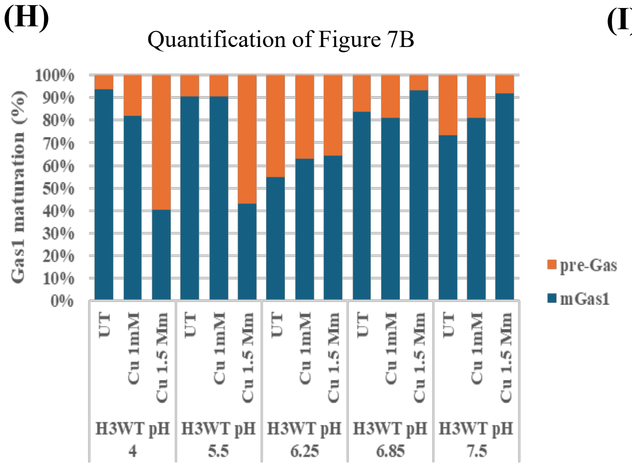


(I)

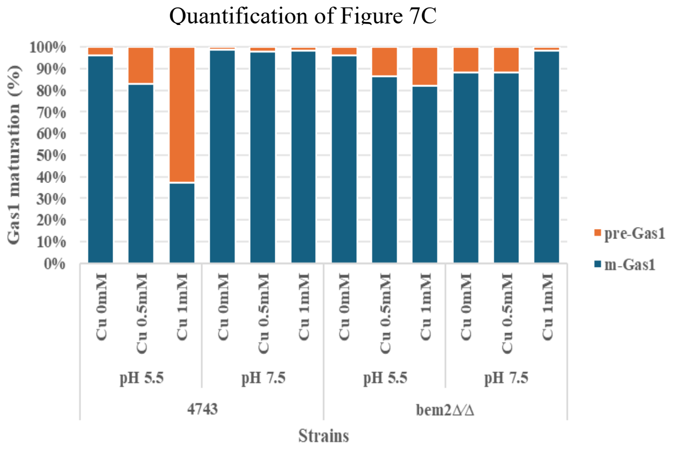


**Figure S7: Effect of redox homeostasis, pH balance, and cell wall homeostasis on copper homeostasis and copper-induced protein translocation defect.** (A) Spot test assay of double deletion mutants of redox homeostasis genes and wild type to test their growth on SC agar medium supplemented with and without copper chloride (0.5, 1, 1.5, 2mM). 10-fold serially diluted cells prepared from overnight grown cultures, spotted from left to right. (B) Biological replicate of western blot presented in Figure 7(A), western blots of Gas1-GFP transformed deletion mutants of redox homeostasis gene (*taz1*Δ/Δ, *trx2*Δ/Δ, *yap1*Δ/Δ and *grx1*Δ/Δ) and respective wild type cells treated with and without copper chloride (0.5 and 1mM) for 2hrs. (C) Spot test assay of copper-sensitive histone mutants and wild-type cells to test the growth on SC agar solid growth medium supplemented with and without copper chloride 1mM at different pH (4.5, 5.5, 6.5, 7), 10-fold serially diluted cells, spotted from left to right. (D) Western blot of Gas1-GFP transformed wild type cells, an overnight grown culture were seeded at 0.2OD600 in SC liquid medium at different pH (4.5, 5.5, 6.5 and 7) and grown till 1OD600, treated and untreated with copper chloride for 2hrs, WCE used for immunoblotting. Immunoblot of TBP served as a protein loading control. WCE means whole cell extract. (E) Spot test assay of double deletion mutants of cell wall homeostasis genes and wild type to test their growth on SC agar medium supplemented with and without copper chloride (1 and 1.5mM) at different pH (4, 5.5, and 7.5), 10-fold serially diluted cells prepared from overnight grown cultures, spotted from left to right. Plates were incubated at 30°C, and images were captured after 72 hours. (F) Biological replicate of western blot presented in Figure 7(C), western blots of Gas1-GFP transformed deletion mutant *bem2Δ/*Δ and respective wild type cells in presence and absence of copper chloride (0.5, 0.75, and 1mM) for 2hrs.(G-I) Quantification of western blots presented in Figure 7(A-C). Data indicates the ratio of pre-Gas1 form and mature Gas1 form in mutants indicated w.r.t. wild type cells.

**Figure S8**

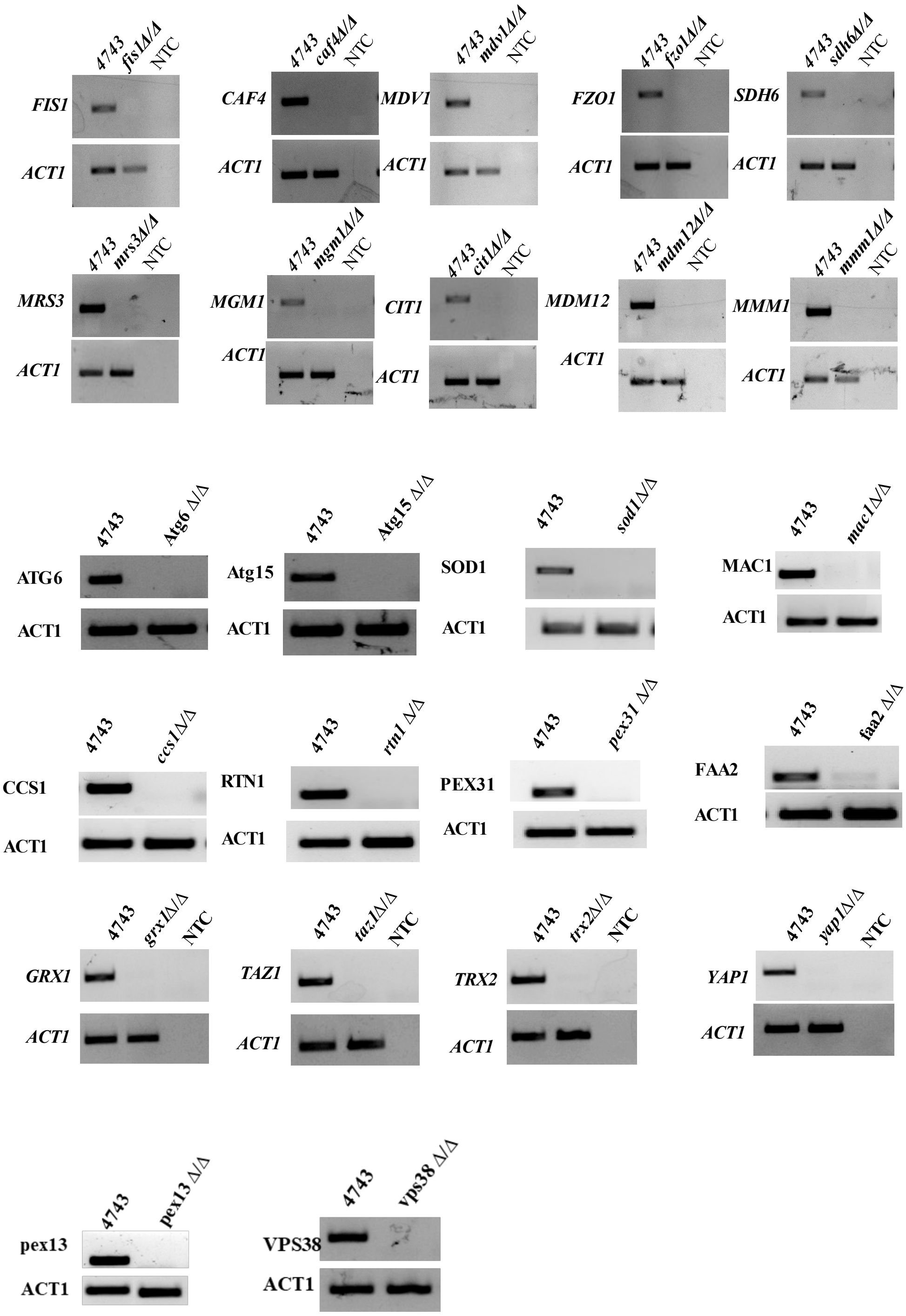


**Figure S8: Confirmation of double-deletion strains through semi-qPCR analysis.** Semi-quantitative PCR of yeast double deletion gene mutations were performed to confirm the strains. Overnight grown cultures of deletion mutants and wild type strains were seeded at 0.2OD600 and grown till 1OD600 in SC liquid media. Harvested and pellet used to isolate gDNA for semi-qPCR analysis, primers used for confirming respective strains are mentioned in Table S2. PCR to amplify the *Act1* (Actin 1) gene serves as a control.

**Table S1. List of strains:**

| Strain | Genotype | Reference |
|---|---|---|
| H3 WT | *MATa his3Δ200 leu2Δ0 lys2Δ0 trp1Δ63 ura3Δ0 met15Δ0 can1::MFA1pr-HIS3 hht1-hhf1::NatMX4 hht2-hhf2::[HHTS-HHFS]*-URA3* | Dharmacon |
| H3Δ(13-16) | Isogenic to H3 WT | Dharmacon |
| H3Δ(13-28) | Isogenic to H3 WT | Dharmacon |
| H3Δ(28-31) | Isogenic to H3 WT | Dharmacon |
| H3Δ(29-32) | Isogenic to H3 WT | Dharmacon |
| H3Δ(4-35) | Isogenic to H3 WT | Dharmacon |
| H3K23Q | Isogenic to H3 WT | Dharmacon |
| H3K27R | Isogenic to H3 WT | Dharmacon |
| H3K36Q | Isogenic to H3 WT | Dharmacon |
| H4 WT | *MATa his3Δ200 leu2Δ0 lys2Δ0 trp1Δ63 ura3Δ0 met15Δ0 can1::MFA1pr-HIS3 hht1-hhf1::NatMX4 hht2-hhf2::[HHTS-HHFS]*-URA3* | Dharmacon |
| H4S1D | Isogenic to H4 WT | Dharmacon |
| H4K5R | Isogenic to H4 WT | Dharmacon |
| H4Δ(9-20) | Isogenic to H4 WT | Dharmacon |
| H4Δ(13-24) | Isogenic to H4 WT | Dharmacon |
| BY4741 | *MATa his3Δ1 leu2Δ met15Δ ura3Δ0* | Open Biosystem |
| H2AΔ(1-20) | *MATa his3-1 leu2-0 met15-0 ura3-0 hht1-hhf1::KAN hhf-2hht2::NAT hta1-htb1::HPH hta2-htb2::NAT p[CEN LEU2 hta1Δ(1–20)-HTB1-HHT2-HHF2]* | (Kim et al., 2012) |
| H2AΔ(1-20) H3K4A | *MATa his3-1 leu2-0 met15-0 ura3-0 hht1-hhf1::KAN hhf-2hht2::NAT hta1-htb1::HPH hta2-htb2::NAT p[CEN LEU2 hta1Δ(1–20)-HTB1-hht2K4A-HHF2]* | (Kim et al., 2012) |
| H2AΔ(1-20) HR2A | *MATa his3-1 leu2-0 met15-0 ura3-0 hht1-hhf1::KAN hhf-2hht2::NAT hta1-htb1::HPH hta2-htb2::NAT p[CEN LEU2 hta1Δ(1–20)-HTB1-hht2R2A-HHF2]* | (Kim et al., 2012) |
| BY4743 | *MATa/α his3Δ1/his3Δ1 leu2Δ0/leu2Δ0 LYS2/lys2Δ0 met15Δ0/MET15 ura3Δ0/ura3Δ0)* | Open Bio system |
| *cup2Δ/ Δ* | *Isogenic to BY4743 cup2Δ/Δ::KANMX* | Open Bio system |
| *mac1Δ/ Δ* | *Isogenic to BY4743 mac1Δ/Δ::KANMX* | Open Bio system |
| *sod1Δ/ Δ* | *Isogenic to BY4743 sod1Δ/Δ::KANMX* | Open Bio system |
| *ccs1Δ/ Δ* | *Isogenic to BY4743 ccs1Δ/Δ::KANMX* | Open Bio system |
| *fis1Δ/Δ* | *Isogenic to BY4743 fis1Δ/Δ::KANMX* | Open Bio system |
| *caf4Δ/Δ* | *Isogenic to BY4743 caf4Δ/Δ::KANMX* | Open Bio system |
| *mdv1Δ/Δ* | *Isogenic to BY4743 mdv1Δ/Δ::KANMX* | Open Bio system |

| | | |
|---|---|---|
| *fzo1Δ/Δ* | *Isogenic to BY4743 fzo1Δ/Δ::KANMX* | Open Bio system |
| *mgm1Δ/Δ* | *Isogenic to BY4743 mgm1Δ/Δ::KANMX* | Open Bio system |
| *cit1Δ/Δ* | *Isogenic to BY4743 cit1Δ/Δ::KANMX* | Open Bio system |
| *sdh6Δ/Δ* | *Isogenic to BY4743 sdh6Δ/Δ::KANMX* | Open Bio system |
| *mrs3Δ/Δ* | *Isogenic to BY4743 mrs3Δ/Δ::KANMX* | Open Bio system |
| *mmm1Δ/Δ* | *Isogenic to BY4743 mmm1Δ/Δ::KANMX* | Open Bio system |
| *mdm12Δ/Δ* | *Isogenic to BY4743 mdm1Δ/Δ::KANMX* | Open Bio system |
| *atg6Δ/Δ* | *Isogenic to BY4743 atg6Δ/Δ::KANMX* | Open Bio system |
| *vps38Δ/Δ* | *Isogenic to BY4743 vps38Δ/Δ::KANMX* | Open Bio system |
| *atg15Δ/Δ* | *Isogenic to BY4743 atg15Δ/Δ::KANMX* | Open Bio system |
| *atg18Δ/Δ* | *Isogenic to BY4743 atg18Δ/Δ::KANMX* | Open Bio system |
| *atg32Δ/Δ* | *Isogenic to BY4743 atg32Δ/Δ::KANMX* | Open Bio system |
| *atg34Δ/Δ* | *Isogenic to BY4743 atg34Δ/Δ::KANMX* | Open Bio system |
| *atg36Δ/Δ* | *Isogenic to BY4743 atg36Δ/Δ::KANMX* | Open Bio system |
| *atg39Δ/Δ* | *Isogenic to BY4743 atg36Δ/Δ::KANMX* | Open Bio system |
| *rtn1Δ/Δ* | *Isogenic to BY4743 rtn1Δ/Δ::KANMX* | Open Bio system |
| *faa2Δ/Δ* | *Isogenic to BY4743 faa2Δ/Δ::KANMX* | Open Bio system |
| *pex31Δ/Δ* | *Isogenic to BY4743 pex31Δ/Δ::KANMX* | Open Bio system |
| *pex29Δ/Δ* | *Isogenic to BY4743 pex29Δ/Δ::KANMX* | Open Bio system |
| *pex13Δ/Δ* | *Isogenic to BY4743 pex13Δ/Δ::KANMX* | Open Bio system |
| *taz1Δ/Δ* | *Isogenic to BY4743 taz1Δ/Δ::KANMX* | Open Bio system |
| *yap1Δ/Δ* | *Isogenic to BY4743 yap1Δ/Δ::KANMX* | Open Bio system |
| *grx1Δ/Δ* | *Isogenic to BY4743 grx1Δ/Δ::KANMX* | Open Bio system |
| *trx2Δ/Δ* | *Isogenic to BY4743 trx2Δ/Δ::KANMX* | Open Bio system |
| *bem2Δ/Δ* | *Isogenic to BY4743 bem2Δ/Δ::KANMX* | Open Bio system |
| Scy62 | *MATa his3-11,15; leu2-3,112; ura3-1; trp1-1; can1-100 ADE2* | (Sandager et al., 2002) |
| *are1Δ* | *Isogenic to* Scy62 *are1Δ::HIS3* | This study |
| *are1Δlro1Δ* | *Isogenic to* Scy62 *are1Δ::HIS3;lro1Δ::TRP1* | This study |
| *are2Δdga1Δ* | *Isogenic to* Scy62 *are2Δ::HIS3;dga1Δ::URA3* | This study |
| *are2Δlro1Δdga1Δ* | *Isogenic to* Scy62 *are2Δ::HIS3;lro1Δ::TRP1;dga1Δ::URA3* | This study |

**Table S2. List of primers:**

| Genes | Primer sequence (5'-3') | References |
|---|---|---|
| *ACT1* | FP: TCGTCGGTAGACCAAGACAC | This study |
| | RP: TTCTTCTGGGGCAACTCTCA | This study |
| *CAT8* | FP: TATCCAAGGGGGAGAACGCA | This study |
| | RP: ATTGGCATCCGTGGCATCTG | This study |
| *HHO1* | FP: AGCCTGCAACCAGCAAAGG | This study |
| | RP: GGATCCGACGATCGGGTAGT | This study |
| *FBP1* | FP: CGCCTCAAAAGGCCATCTACT | This study |
| | RP: GTCGCAAGGGTATGCGAAAAG | This study |
| *SOD1* | FP: GTGTCTCTGCTGGTCCTCAC | This study |
| | RP: GGATAACGACGCTTCTGCCT | This study |
| *CCS1* | FP: AGGGCGTGGAATCTACTGGT | This study |
| | RP: CTTAACGGAGGAAGGCTCGTT | This study |
| *MAC1* | FP: GGCATCGCTCTTCAACATGC | This study |
| | RP: GGCACGTACAAACAGTATTGGC | This study |
| *FIS1* | FP: TTAACTACGCATGGGGGCTG | This study |
| | RP: TACTCTTCAAAGCGCCCACC | This study |
| *CAF4* | FP: GGCCCGAAAGTACAAAAGAT | This study |
| | RP: CTGTTGAATGATGGAAACGG | This study |
| *MDV1* | FP: GCCTAGATTTTGATGCGCCC | This study |
| | RP: AAAGCCGCATCTCTACCACC | This study |
| *FZO1* | FP: GAGGATGATTTGTTGCCCT | This study |
| | RP: TTTTCCCGTTCATCATTTGC | This study |
| *MGM1* | FP: TGAGAGCAGAATTGGATCAG | This study |
| | RP: TGCATTGGTAGTAAGCATTG | This study |
| *CIT1* | FP: CTTTTGGGCTACGAAAACAA | This study |
| | RP: AAAACTTCTTGATTGGCACG | This study |
| *SDH6* | FP: TTACACCTGTATAGGGCTTC | This study |
| | RP: TGTATATTCGTCAGTTCAGGG | This study |

| | | |
|---|---|---|
| *MRS3* | FP: TCTGTGTGGCAGTATCAGCG | This study |
| | RP: TCCTTGGTTTCCAACCTCTCC | This study |
| *MMM1* | FP: GGTTTCAGGCGTGTTTGACC | This study |
| | RP: GACACTGCCGATCTTGGGAA | This study |
| *MDM12* | FP: TACGATTGGTGCCGACCTTG | This study |
| | RP: TACTGAGCCCTGTCCCTGAT | This study |
| *ATG6* | FP: TCAGCAACAAGGGCATCGAA | This study |
| | RP: ACGCTGGTCCTCTCTGCTATAA | This study |
| *ATG15* | FP: CTTTTGGATTGCCTGCGGTC | This study |
| | RP: CCCACCAGTGAGCAACTTGA | This study |
| *VPS38* | FP: ATTACAGGCACAACGCAC | This study |
| | RP: TCCTCATGGACTTGGGAAGA | This study |
| *RTN1* | FP: CAGCGAGGGTCAGTTATCAACA | This study |
| | RP: CATCTTCCATCAAGCCGCCA | This study |
| *FAA2* | FP: TACCACTGACCAGCTATCCCAG | This study |
| | RP: ACGTTGAGGCCATCTCCTTG | This study |
| *PEX31* | FP: GGCTTGGGAATGGCACGATA | This study |
| | RP: CAGTCCTGATCCACCTCCTTCT | This study |
| *PEX29* | FP: CTCGCATCCGTCCAATCGT | This study |
| | RP: CAACTGGGCGCCATTCTTTG | This study |
| *PEX13* | FP: CGACCTAAACCTTGGGAGACC | This study |
| | RP: GATTCGCCGTAGGTACCAGAG | This study |
| *TRX2* | FP: AGCATCTGGCGACAAGTTAGT | This study |
| | RP: CGCCCTTGTAGAAGATTAGGGT | This study |
| *TAZ1* | FP: TGGAATGGACCCCTCACTCT | This study |
| | RP: AATGGGCGGCTTTGTTGC | This study |
| *GRX1* | FP: ACGTACTGTCCATACTGCCA | This study |
| | RP: AAGTCGTCGTTGCCTCCAAT | This study |
| *YAP1* | FP: TACACGTGATGGCGAGGATA | This study |
| | RP: CCACTTCATTTTGCTGCTGA | This study |

FP: forward primer, RP: Reverse primer

**Table S3. List of plasmids used in this study: -**

| Plasmid name | References |
|---|---|
| pFA-6a-13myc | This study |
| pRS415-Gas1-GFP | (Ha et al., 2014) |
